# Calpain DEK1 acts as a developmental switch gatekeeping cell fate transitions

**DOI:** 10.1101/2021.08.25.457637

**Authors:** Viktor Demko, Tatiana Belova, Maxim Messerer, Torgeir R. Hvidsten, Pierre-François Perroud, Ako Eugene Ako, Wenche Johansen, Klaus F.X. Mayer, Odd-Arne Olsen, Daniel Lang

## Abstract

Calpains are cysteine proteases that control cell fate transitions. Although calpains are viewed as modulatory proteases displaying severe, pleiotropic phenotypes in eukaryotes, human calpain targets are also directed to the N-end rule degradatory pathway. Several of these destabilized targets are transcription factors, hinting at a gene regulatory role. Here, we analyze the gene regulatory networks of *Physcomitrium patens* and characterize the regulons that are deregulated in *DEK1* calpain mutants. Predicted cleavage patterns of regulatory hierarchies in the five DEK1-controlled subnetworks are consistent with the gene’s pleiotropy and the regulatory role in cell fate transitions targeting a broad spectrum of functions. Network structure suggests DEK1-gated sequential transition between cell fates in 2D to 3D development. We anticipate that both our method combining phenotyping, transcriptomics and data science to dissect phenotypic traits and our model explaining the calpain’s role as a switch gatekeeping cell fate transitions will inform biology beyond plant development.

## Introduction

Multicellular organisms are dependent on the establishment of distinct cellular identities to implement individual tissues, cell types and functions. In the course of development, cells acquire specific cellular fates resulting in either division, differentiation or death. A cell fate is the product of a specific expression state, leading to a distinct cellular complement of RNAs and proteins that provide a specific competence to respond to environmental signals and developmental patterning cues (Pierre-Jerome et al., 2018; Casey et al., 2020). The transition between different cellular fates requires both the reprogramming of the gene expression state as well as the modulation or exchange of the cell’s protein complement. In many instances, cell fate transitions are the result of asymmetric cell division. In plants, cell fate is dependent on spatial localization and intercellular communication via direct protein-protein interactions or mobile signals like phytohormones, peptides or transcription factors (Pierre-Jerome et al., 2018; Gundu et al., 2020; Hata and Kyozuka, 2021). This process must be highly coordinated in order to synchronously implement the changes on an epigenetic, transcriptional, post-transcriptional, translational and post-translational level.

Asymmetric, formative cell divisions mostly involve a reorientation of the division plane and depend on both intrinsic and extrinsic factors (Shao and Dong, 2016; de Keijzer et al., 2021). This divisional asymmetry requires direct modification or exchange of cytoskeletal components or other proteins and protein complexes affecting cell polarity. Thus, plant cell fate transitions depend on pathways implementing the physical asymmetry of cell division as well as the regulatory (re)programming of divergent cellular identities. Dependence on positional and other extrinsic signals necessitates the tight integration of all physical and regulatory layers determining plant cell fate. How are the different physical and regulatory layers coordinated and synchronized to a new cell fate and how is the integration of potentially conflicting extrinsic signals and intrinsic factors implemented?

The endosperm aleurone as well as the protodermal or the derived epidermal L1 layers of flowering plants are prime examples for cell fate specifications that depend on these processes. Both cell types implement the outer layers of the respective flowering plant tissues, emerge by asymmetric cell division and are specified by positional information (Takada and Iida, 2014; Olsen, 2020). Remarkably, these cell fate specifications fail in deletion or loss of function mutants of the *DEFECTIVE KERNEL 1* (*DEK1*) gene in mono-(Becraft et al., 2002; Lid et al., 2002) and dicotyledonous (Lid et al., 2005; Johnson et al., 2005) plants. The failed embryonal cell fate transition has tremendous impact on further plant development – it is arrested at the globular embryo stage, because the L1 layer is crucial to the regulation of shoot stem cell maintenance and organ growth (Takada and Iida, 2014). *DEK1* mutants in various organisms display pleiotropic phenotypes and are always severely impeded in further development (Ahn et al., 2004; Lid et al., 2005; Hibara et al., 2009; Liang et al., 2015; Amanda et al., 2017). DEK1 is essential for flowering plant seed, embryo, epidermis, trichome, leaf and shoot apical meristem development (Olsen et al., 2015). Consistent with the assessment of DEK1 as one of the key components in land plant meristem evolution (Harrison, 2015), studies in the bryophyte model system *Physcomitrium patens* (Physcomitrella; Rensing et al., 2020) demonstrated a vital role as a developmental regulator controlling cell fate decisions in the moss’ simplex meristems involved in the 2D to 3D transition (Perroud et al., 2014; Demko et al., 2014).

At the molecular level, DEK1 seems to affect both physical and regulatory layers governing cell fate transitions. In terms of the physical, asymmetry determining factors, *DEK1* null mutants display disorganization of division planes resulting in defective division patterns and cell shapes (Lid et al., 2005; Perroud et al., 2014). These phenotypes are caused by defects in microtubule-organized orientation (Liang et al., 2015), the deposition and remodeling of the cell wall as well as cell adhesion (Amanda et al., 2016). Changes in gene expression levels and patterns hint towards a role also in the regulatory layers. These comprise the deregulation of several cell type-specific markers like the HD-ZIP family IV transcription factors (TFs: ML1, PDF2, HDG11, HDG12, and HDG2; Galletti et al., 2015), as well as other embryo- and post-embryo developmental regulators (e.g. CLV3, STM, WOX2, WUS, PIN4; Liang et al., 2015) and potential downstream target genes involved in cell wall biosynthesis and remodeling and similar functions related to the differentiation of the respective cell type (XTH19, XTH31, PME35, GAUT1, CGR2, EXP11; Amanda et al., 2016; Amanda et al., 2017).

DEK1 is a 240 kDa multi-pass transmembrane (TM) protein with a cytosolic calpain cysteine protease (*CysPc* and *C2L* domains) as the effector (Fig. 1a). Other structural features include a long cytosolic *loop* region between TM domains as well as an *LG3*-like domain in the cytosolic *linker* connecting the membrane domain (*MEM*) with the calpain protease domain (*CysPc-C2L;* Demko et al., 2014; Johansen et al., 2016). The DEK1 calpains comprise a subfamily of a deeply conserved, multi-domain superfamily of calpains sharing a common *CysPc* domain that likely evolved in eubacteria (Zhao et al., 2012). The clade of the animal-specific, so-called classical calpains are mechanistically the best characterized and have been described as the third proteolytic system besides the lysosomal and proteasomal systems (Spinozzi et al., 2021). Contrary to the latter two degradatory systems, the Ca^2+^-dependent calpains are often described as modulatory proteases that cleave proteins at a limited number of specific sites, generating fragments or neo-proteins with novel functions (Zhao et al., 2012; Cardoso et al., 2019). These e.g. comprise maturation from an inactive preprotein, modulation of stability and association of multiprotein complexes or modification of subcellular localizations. Like plant DEK1 calpains, classical calpains are pivotal to animal development and cell fate transitions with roles in early embryonic patterning, muscle and neural development and homeostasis, cell migration, Wnt signalling during tissue morphogenesis and axial patterning via the NFκB/IκB pathway (Araujo et al., 2018). The functional roles in the regulation of cell division and the cell cycle seems to be the ancestral state of the entire superfamily (Zhao et al., 2012; Vieira et al., 2017). Consistent with the broad functional importance of the calpain system as well as the pleiotropy and severity of loss of function mutants in various eukaryotic model systems, human calpains have been found as aggravating factors in many pathophysiological conditions and illnesses including various cancers and hereditary diseases like muscular dystrophy (Storr et al., 2011; Ono and Sorimachi, 2012; Ono et al., 2016; Araujo et al., 2018).

**Figure 1.**
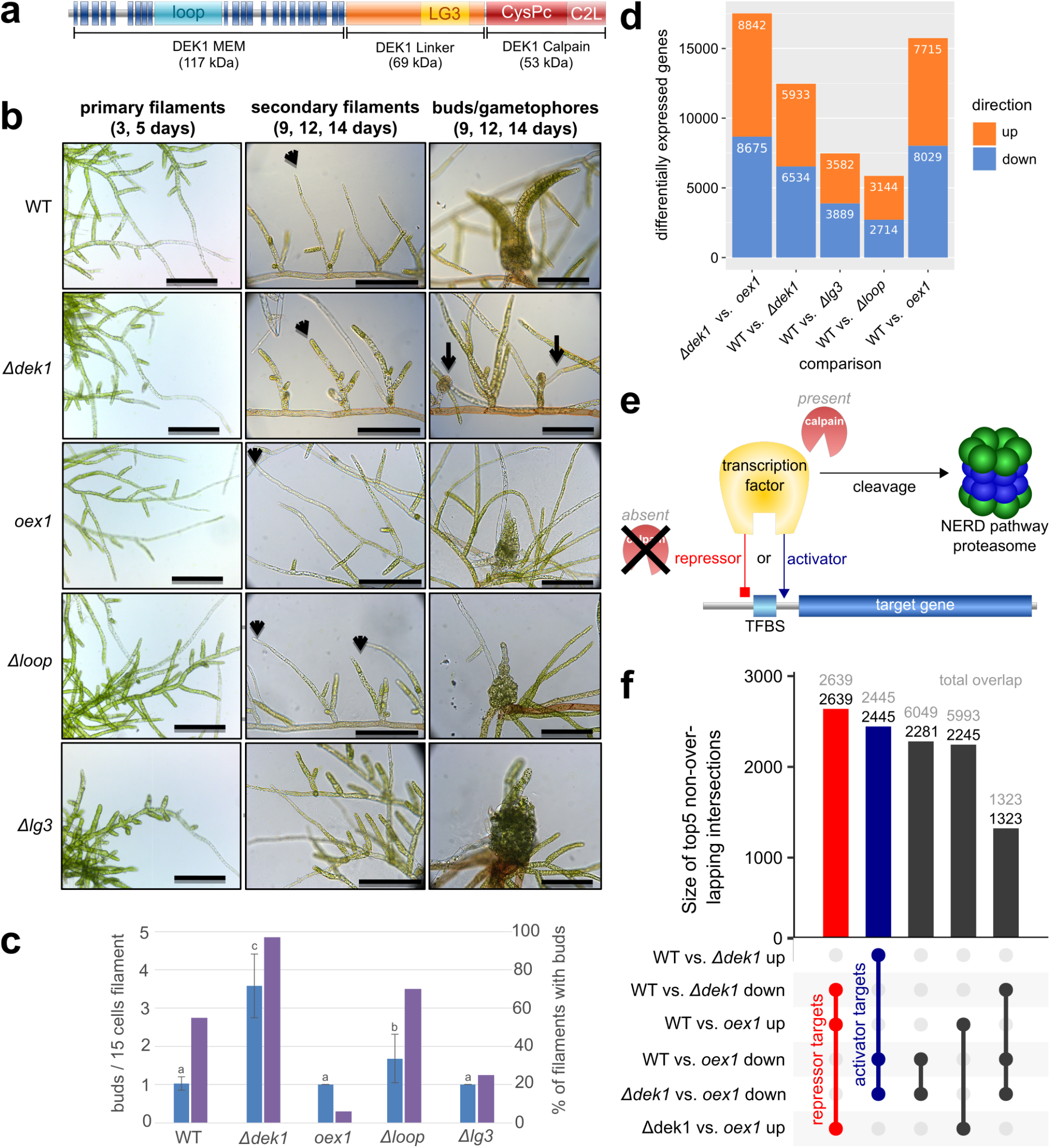
Phenotypes and transcriptome of *DEK1* mutant lines in *P. patens*. **a.** DEK1 protein domain structure. **b.** Time series analysis of *P. patens* juvenile gametophyte development in wild type (<u>WT</u>), a DEK1-Calpain domain over-expressor (*<u>oex1</u>*), a complete deletion of the DEK1 gene (*<u>Δdek1</u>*; Perroud et al. 2014) as well as two partial deletion lines lacking the loop (*<u>Δloop</u>*; Demko et al., 2014) and the LG3 domain (*<u>Δlg3</u>*; Johansen et al. 2016), respectively. Microscopic images (scale bars: 200μm) depict primary filaments in early stages of protonemata development (left column), secondary filaments (middle column, arrowheads point to apical cells of individual secondary filaments), buds and gametophores (right column, arrows point to arrested buds in *Δdek1*). **c.** Quantitative analysis of gametophore apical meristem (bud) formation in *DEK1* mutants. Frequency of meristem initiation expressed as <u>number of buds per 15 cell long filament</u> (blue bars; left x-axis) and <u>percentage of filaments forming buds</u> (purple bars; right x-axis). Statistical significance at 95% confidence is indicated for mean values (a, b, c). Analysis of variance (ANOVA) and least significant difference (LSD) was performed in multiple sample comparisons. **d.** Pairwise, differential time-series gene expression analysis of *DEK1* mutants at 3, 5, 9, 12 and 14 days. Stacked bar chart of significantly <u>differentially expressed genes</u> (DEGs) at false discovery rate (FDR) < 0.1. Fill colors correspond to <u>up</u> (orange) and <u>down</u>-regulated (lightblue) genes. **e.** Model hypothesis for the gene regulatory role of DEK1/calpains. Model depicts two states: When the active calpain is <u>present</u>: TF (transcription factor) is cleaved and targeted to the N-end rule degradatory (NERD) pathway, resulting in lack of gene regulation. In the <u>absence</u> of an active calpain the TF regulates target gene expression either as an <u>activator</u> (blue) or <u>repressor</u> (red). **f.** Size of the top 5 intersections between the sets of DEGs in Panel 1d. Numbers above the bars depict the part that is unique to the given set (black) as well as the total (grey) size of each intersection. The two largest unique intersections correspond to expression profiles consistent with a reciprocal deregulation in *Δdek1* and *oex1*. The largest set (red) comprises 2,639 genes which are down-regulated in *Δdek1* and up-regulated in *oex1*. This profile is consistent with the hypothesis that these genes are targets of DEK1-controlled <u>repressors</u>. The second largest set (blue) comprises 2,445 genes that are up-regulated in *Δdek1* and down-regulated in *oex1.* These genes are likely controlled by DEK1-targeted <u>activators</u>. These sets represent the most conservative lists, as e.g. the third intersection certainly contains additional *activator targets* with weak FDR support in the comparison of *Δdek1* and the WT.

The *CysPc-C2L* protease domain of DEK1 calpains is functionally conserved among land plants (Liang et al., 2013). It comprises the catalytic sites also found in the classical calpains and is a functional cysteine protease (Wang et al., 2003). Although *in vitro* catalytic activity does not depend on Ca^2+^, the activity increases in the presence of added mM levels of Ca^2+^ (Wang et al., 2003). Like classical calpains, the full DEK1 protein demonstrates autoproteolytic activity releasing several soluble neo-proteins with cytosolic localization (Johnson et al., 2008). The membrane-localized full-length protein thus likely represents the inactive calpain that might be activated by phosphorylation (Moody, 2019), mechanosensing (Tran et al., 2017) or other sources emitting local Ca^2+^ pulses similar to the classical calpains (Araujo et a., 2018). All of this, taken together with the phylogeny, phenotypic data and the observed broad functionality that is expressed by conserved functions in basic biological processes and convergently evolved roles in development, points to a deep conservation of the *CysPc-C2L* protease domain’s molecular function. The alternate domain architectures of the superfamily evolved by domain shuffling and likely accommodate independent, but convergent ways to modulate, specify or restrict the catalytic activity of the calpain protease (Zhao et al., 2012).

Calpains display a fuzzy target specificity that is less reliant on the primary sequence of the substrate and possibly depends on higher order factors like 2D/3D protein structure or cofactors (Ono and Sorimachi, 2012; Araujo et al., 2018). Coupled with the modulatory role of the protease and the broadness of the involved biological processes, the seeming lack of specificity has been a limiting factor for the mechanistic exploration and systematic identification of calpain targets (Ono et al., 2016). As a result, so far there are a few hundred confirmed calpain substrates, most of which have been reported in mammals. In plants, so far no direct substrates of the DEK1 calpain have been identified. Generally, it has been extremely difficult to mechanistically align the view of a non-degradatory, modulatory but relatively unspecific protease with the diversity, ubiquity, severity and impact of calpain defects and mutations.

*In vitro* and *in vivo* degradation assays of confirmed mammalian calpain targets (Piatkov et al., 2014) revealed them as short-lived substrates for the N-end rule degradation or N-degron pathway (NERD; Majovsky et al., 2014; Nguyen et al., 2018; Holdsworth et al., 2020). Natural C-terminal fragments of a variety of mammalian proteins, including transcriptional regulators, bear N-terminal residues, so-called N-degrons, that attract and activate the components of the NERD pathway and ultimately lead to their degradation by the ubiquitin-proteasome system (Piatkov et al., 2014). This also explains the short half-life of auto-proteolytic cleavages in mouse and fruit fly calpains resulting in N-degron fragments (Farkas et al., 2004; Piatkov et al., 2014). Thus, in these instances calpain cleavage leads to destabilization or inhibition of biological functions, providing a plausible explanation to align the limited target specificity with the broad biological impact of the cysteine proteases.

Considering that the targets of such cleavages also comprise TFs and other transcriptional regulators, a more direct route to the control of both the physical and regulatory layers of cell fate transitions emerges. In this case, calpains act as post-translational regulators of gene functions either a) by directly modulating the proteins’ function, b) by indirectly affecting the target protein’s stability marking them for the NERD pathway or c) by indirectly controlling the stability of the gene’s transcriptional regulators (Fig. 1e). While a) represents a positive control over a gene’s function, b) an inhibitory, negative control, the outcome of c) depends on whether the targeted TF or regulator represents a transcriptional activator or repressor and thus allows a bidirectional control of gene functions.

Due to its subcellular localization in the plasma membrane and its role in the establishment of cell division plane orientation, so far studies of the plant calpain have been designed with a scenario in mind where DEK1 only acts as a modulatory protease (a) targeting a limited number of specific targets e.g. in or around the cell division plane apparatus. Combining the pleiotropy and developmental essentiality in *DEK1* mutants, the fuzzy target specificity and the broad range of functions with the inhibitory role in targeting proteins to the NERD pathway, we postulate a second scenario (*dual role scenario*): the calpain directly modulates specific protein functions (a) and indirectly controls the half-life of potentially many proteins (b). These include transcriptional regulators and thereby affect the expression of a large number of target genes (c).

The model moss *P. patens* provides an ideal system to evaluate these two scenarios. Generally, its evolutionary position as a representative of the earliest diverging clade of land plants, the available reference genome and high-quality annotation resources (Lang et al., 2018), the well-established molecular toolbox and the comprehensive transcriptomics resources covering the entire life cycle (e.g. Frank and Scanlon, 2015; Perroud et al., 2018; Kirbis et al., 2020) make it an attractive model of choice for land plant evo-devo studies like ours (Rensing et al, 2020; Hata and Kyozuka, 2021; Keijzer et al., 2021). Specifically, the simple morphological structures in the early development of the moss, comprising mostly single-cell-layered tissues, is not impeded in the *DEK1* null mutants (Perroud et al., 2014; Demko et al., 2014; Johansen et al., 2016) and thus facilitate transcriptional and phenotypic profiling along a longer developmental time course including the 2D to 3D transition of the moss’ simplex shoot meristem. The NERD pathway is conserved among land plants and *P. patens* is ideally suited to assess the calpain’s potential role as a post-translation gene regulator feeding into this pathway, because mutants of the moss’ NERD pathway display a phenotype that is highly similar to DEK1 (Hoernstein et al., 2016; Schuessele et al., 2016).

We set out to elucidate DEK1’s role in plant development and gene regulation by combining phenotypic and transcriptional profiling of wild-type and mutant moss lines with a comprehensive, genome-wide, integrative, multi-scale data mining approach to analyze the mutant deregulation profiles of the global gene regulatory networks. Our analysis sheds light on the molecular nature underlying the pleiotropic developmental phenotypes caused by *DEK1*. The predicted gene regulatory network allows to connect the calpain’s biochemical function and regulatory consequences supporting its role as a post-translational gene regulator. We combined phenotyping, transcriptomics and data science to dissect phenotypic traits. The annotated, genome-wide gene regulatory network is a valuable resource facilitating next generation plant evo-devo research. The proposed model explains the calpain’s role as a developmental switch gatekeeping cell fate transitions.

## Results

### DEK1 dramatically impacts moss development

Previous work demonstrated that mutations in the *DEK1* gene dramatically affect *P. patens* development (Olsen et al., 2015). Here, we characterize and quantify phenotypes and global gene expression profiles of five moss lines. In addition to wild-type (WT), a *Δdek1* deletion strain (Perroud et al., 2014), a strain overexpressing the DEK1 linker and calpain domains (Fig. 1a) under the maize ubiquitin promoter (*oex1*) and two partial domain-deletion lines, *Δloop* (Demko et at., 2014) and *Δlg3* (Johansen et al., 2016), were studied (Fig.1a). These lines were sampled at five time points that comprise the early and intermediate stages of *P. patens* development, including the transition from 2D tip-growth in the filamentous protonema to 3D apical growth in leafy shoots (gametophores; Fig. 1b). While the early stages of protonema development are largely unaltered in the *Δdek1* and *Δloop* mutants (Fig. 1b), the later stages as well as the formation of gametophores are significantly disturbed in all mutant strains.

Comparison of the developmental defects in the deletion and overexpression lines, highlights systematic, diametrically opposing effects of the two mutation types as compared to WT. These diametrical phenotypes include reduced (*Δdek1*) vs. enhanced (*oex1*) secondary filament extension, higher vs. lower percentage of filaments that form buds (*Δdek1*/*oex1*) and a four times higher gametophore bud initiation rate per filament in *Δdek1* (Fig. 1b,c).

The partial *Δloop* deletion line shows milder phenotypic changes than *Δdek1*, except for bud development which, instead of stopping, continues to proliferate forming naked stems without initiation of phyllids. *Δlg3* displays unique phenotypes including severely affected protonema differentiation and branching resulting in reduced plant size, as well as aberrant gametophore formation. Juvenile *Δlg3* plants display stunted phyllids.

How can altering one membrane-bound protease have such variable, complex and drastic effects on plant development? The clear, diametrical behaviour of the deletion and overexpression lines, as well as the intermediate phenotypes of the partial deletions, provide the opportunity to shed light into this question by evaluating the global, molecular consequences of *DEK1* mutation using differential transcriptomics.

### Many genes and functions are deregulated in *DEK1* mutants

We performed differential gene expression (DGE) analysis based on triplicate RNAseq libraries generated for the developmental time course described above. Expression levels for all non-redundant transcripts of protein- and non-protein coding genes were quantified, condensed per gene and subsequently tested for DGE along the time course using kallisto/sleuth (Bray et al., 2016; Pimentel et al., 2017). Sets of up- and down-regulated genes in the mutants were identified at 10% FDR (Fig 1d; see Table ST1 for statistics with FDR 10% and 1%). The impact of *DEK1* mutation on overall gene regulation can be defined as the total sum of differentially expressed or up- and down-regulated genes, i.e. the total number of deregulated genes in each mutant line. Consistent with the dramatic phenotypic consequences in *Δdek1*, *oex1*, *Δloop* and *Δlg3*, the DGE analysis revealed a substantial number of deregulated genes in the mutants. The largest number of deregulated genes is observed in the *Δdek1* (35% of all genes) and *oex1* (44% in comparison to wild type and 49% relative to *Δdek1* levels) lines. Whereas in the partial deletion lines *Δloop* and *Δlg3* about 7% of all genes are deregulated. Overall, we observe balanced directionality of deregulation in all studied contrasts, i.e. similar number of up and down-regulated genes.

Overall, the extent of deregulated genes supports a scenario where the DEK1 calpain controls the half-life of transcriptional regulators and thereby indirectly affects the expression of a large number of target genes (*dual role scenario*). This also allows us to explain the observed balanced directionality of deregulated genes that is consistent with a uniform distribution of direct DEK1 cleavage targets, comprising equal proportions of gene activators and repressors. The lack of a few, specific, modulatory calpain cleavages alone unlikely leads to such pronounced transcriptional changes. Such specific action likely would result in only specific functions to be deregulated. To delineate the functional consequences of mutating *DEK1*, we utilized Gene and Plant Ontology annotations to assess the percentages of terms and genes that are deregulated in the mutants (Fig. SF1; Table ST2). Consistent with a broad target prevalence of DEK1 (*dual role scenario*) and the observed complex, pleiotropic phenotype, the analysis demonstrates that 85% of molecular functions, 88% of biological processes, 90% of cellular components, 92% anatomical entities and 94% of all developmental stages in GO and PO are deregulated in *Δdek1*.

### Potential indirect DEK1 targets display consistent mutant deregulation patterns

Given the *dual role scenario*, DEK1 acts as a post-translational regulator (Fig. 1e) of ubiquitous gene functions. This should be reflected in the global expression patterns of the mutants: Down-stream genes of repressors that are cleaved by DEK1 at the protein-level, will be up-regulated in the over-expressor and down-regulated in the null mutant (referred to as *repressor targets* below). Down-stream genes of DEK1-controlled activators should respond in a diametrically opposite fashion (*activator targets*).

To assess this, we performed multiple set analysis of the differentially expressed genes to identify dominant deregulation patterns (Fig. 1f). Indeed, the two top emerging patterns comprise consistently deregulated, diametrically responding genes. The largest set (red bar; Fig. 1f) comprises 2,639 genes which are down-regulated in *Δdek1* and up-regulated in *oex1*. This profile is consistent with the hypothesis that these genes are targets of DEK1-controlled repressors (Fig. 1e). The second largest set (blue bar; Fig. 1f) comprises 2,445 genes that are up-regulated in *Δdek1* and down-regulated in *oex1*. These genes are potentially controlled by DEK1-targeted activators (Fig. 1e). The third and fourth most frequent patterns are subsets of these two patterns. Both gene sets also likely comprise indirect targets of DEK1. Thus, the dominant, global expression patterns of deregulated genes are in line with the *dual role scenario* and all of these deregulated genes can be considered as indirect DEK1 targets. In the subsequent analyses, we focused on the two major sets of consistently deregulated genes as potential indirect DEK1 *repressor* and *activator targets* (Fig. 1f).

As a next step, we profiled gene expression during wild type gametophyte development and detected 71% and 75% of the putative DEK1-controlled *repressor* and *activator targets* respectively displaying significant expression level changes. The two sets substantially differ in their expression patterns along the time course. For example, while 765 of the *repressor targets* are more active during the early phase (days 3-5, Fig. 1b), 794 of the *activator targets* are up-regulated during the later phases of development (days 9-14). This might indicate that some of these potential DEK1-targets are involved in the developmental transitions occurring during this time interval. Taken together, the evidence from the deregulated transcriptomes of *DEK1* mutants and the wild type is consistent with the role of the plant calpain acting as a post-translational regulator with a broad range of direct or indirect targets and encoded gene functions whose timed expression is consistent with the observed, severe and pleiotropic mutant phenotype.

### 5 out of 11 GRN subnetworks are enriched for putative DEK1 targets

To further evaluate the *dual role scenario*, we set out to look for patterns of consistent deregulation in the moss’ gene regulatory networks (GRNs). As a basis, we predicted genome-wide regulatory interactions based on 374 public and novel, paired-end RNASeq libraries and 1,736 novel annotated regulators using the Random Forest predictor of GENIE3 (Huynh-Thu et al., 2010). The overall directionality of the regulatory interactions, i.e. positive (activating) or negative (repressive) regulation, was determined by the correlation coefficient of regulator and target gene expression levels along the developmental time course. Subsequent community detection of the top10 regulatory interactions for 35,706 genes resulted in 11 robust subnetworks (Fig. SF2a). The global analysis and functional composition of the subnetworks is beyond the scope of this study (manuscript in preparation). Here, we focus on the predicted regulatory interactions and subnetworks as tools to assess the putative gene regulatory role of DEK1.

We employed network enrichment analysis of candidate gene sets inferred from the DGE analyses in the subnetwork graphs (Signorelli et al., 2016). This revealed significant enrichment of regulatory relationships for DEK1-controlled *repressor* and *activator targets* in subnetworks II, V VIII, IX and X (FDR << 0.01; rows 1 and 3 of Fig. 2a; Fig. SF2b). Based on peak activities of gene expression during wild type gametophyte development, subnetwork V is enriched for *repressor targets* active during the early phase comprising mostly chloronema filaments (3-5d; row 2 Fig. 2a). Subnetworks II and X are enriched for *activator targets* which are expressed during the 2D to 3D transition (9-14d; row 5 Fig. 2a). By tracing the regulatory links of deregulated genes, the GRN allowed us to identify upstream TFs representing potential direct cleavage targets of the DEK1 calpain with unaffected expression in the mutants. Consequently, we tested subnetworks for enrichment of such upstream TFs predicted to directly control any of the deregulated genes. While this analysis again highlighted subnetworks II, VIII, and X, it also suggested potential direct cleavage targets in three other subnetworks (I, III and XI; row 4 in Fig. 2a). As the deregulated targets of these TFs predominantly also fall in the five DEK1-controlled subnetworks, the latter group of regulators might serve as DEK1-controlled interfaces to other regulatory circuits.

**Figure 2.**
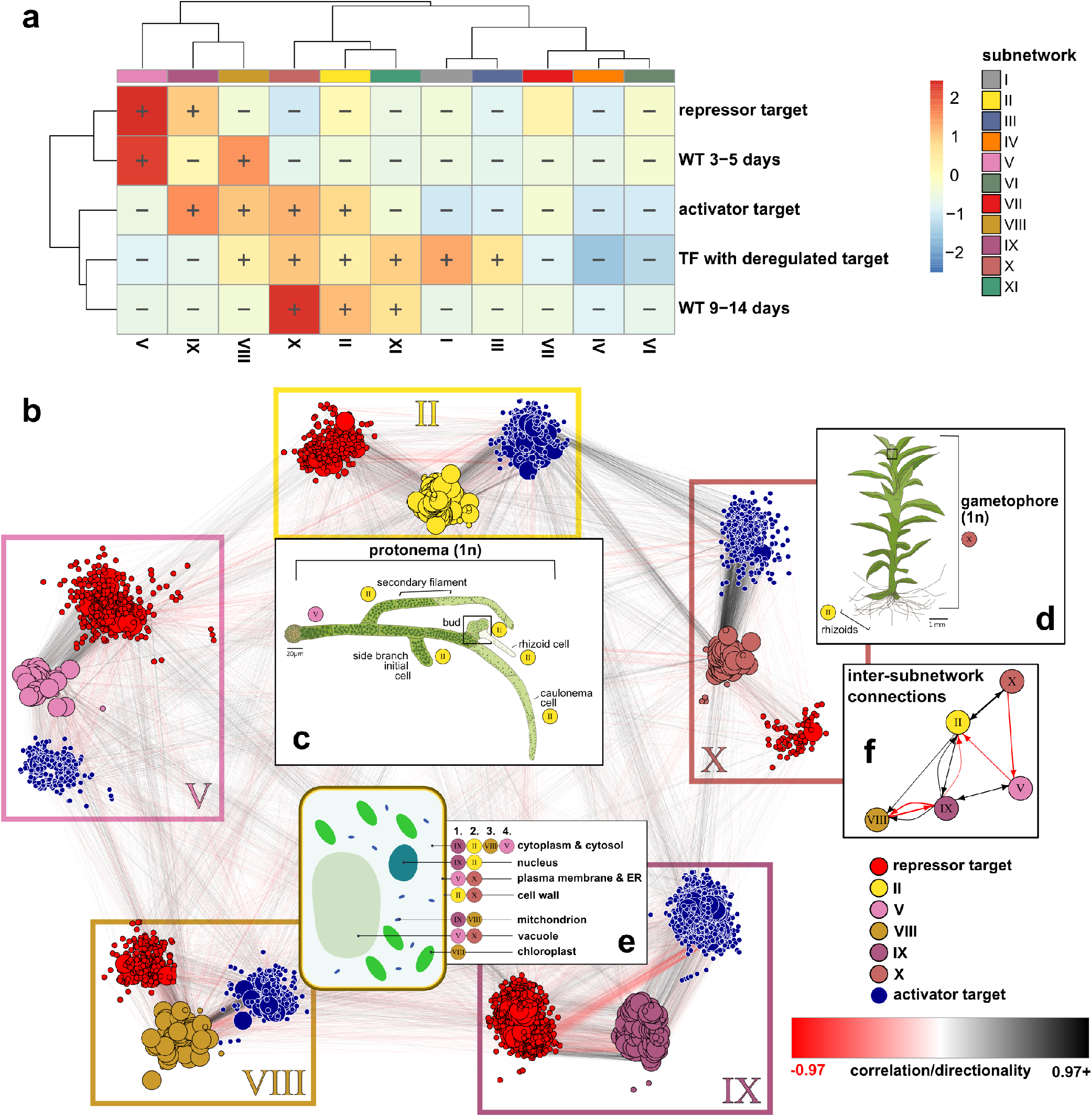
Tracing DEK1-deregulated genes and their upstream regulators in the predicted *P. patens* gene regulatory network (GRN) highlights the subnetworks that are encoding the developmental transitions governed by the calpain. Prediction of regulatory interactions with subsequent community clustering resulted in 11 subnetworks (Fig. SF2a). **a.** Network enrichment analysis highlights specific subnetworks as over-represented among DEK1-deregulated gene sets (<u>activator</u> and <u>repressor targets</u>) and their upstream regulators (<u>TF with deregulated target</u>) as well as the distinct phases of wild type development (<u>WT 3-5</u> and <u>9-14 days</u>). Heatmap depicting ratio of observed vs expected sizes of specific candidate gene sets among the identified subnetworks. Significant (FDR < 0.01) enrichment (+) or depletion (-) is presented by respective heat map cell annotations. Ratios were clustered for rows and columns using the ward.D2 method. **b.** Network graph of the five DEK1-deregulated subnetworks: For each enriched subnetwork (II, V, VIII, IX and X), all genes with an <u>activator</u> (blue nodes) or <u>repressor</u> (red nodes) deregulation pattern in the *DEK1* mutants (Fig. 1f) are depicted as nodes together with <u>unchanged, direct upstream TFs</u> (using subnetwork color code) as a triangular subgraph in subnetwork-colour-framed boxes. Node sizes are scaled by local-reaching centrality, i.e. the fraction of the total subnetwork that can be reached via regulator→target connections. Edges, representing the predicted regulatory interaction between a TF and its target, are colored according to putative <u>directionality</u>. According to this color scheme, negative, repressive interactions are red and positive, activative regulatory interactions are black. Inlets **c.** and **d.** depict significantly enriched developmental stages and tissue or cell types. **c.** Drawing of the predicted roles of subnetworks V and II in the different cell fates comprising the haploid <u>protonema</u> stage. **d.** Drawing of the haploid, leafy, juvenile <u>gametophore</u> which, except for the filamentous rhizoids that are encoded by subnetwork II, is predominantly implemented by subnetwork X. **e.** Drawing of a <u>plant cell</u> depicting the significantly enriched <u>intracellular localizations</u> of the DEK1-controlled subnetworks. In the accompanying text box, subnetworks are ranked (1.-4.) according to the percentages of genes with terms affiliated with the respective compartment. **f.** Small network plot depicting the major, significantly enriched inter-subnetwork connections (Pearson residuals > 4; Figure SF4). Drawings of the *Physcomitrella* protonema (c.) and gametophore (d.) stage adapted from Wu et al. 2018. Drawing of a plant cell (e.) adapted from Wikimedia Commons User *Domdomegg*. Subnetwork assignments to developmental stages and cell types (c. and d.) are based on network enrichment of stage-specific DGE sets inferred from *Physcomitrella* gene atlas data (Perroud et al. 2018). Ranked subnetwork assignment of subcellular localizations (e.) is based on ontology enrichment analysis of GO cellular component terms (Figure SF3 and Tables ST3).

Jointly these analyses suggest that the set of DEK1-deregulated genes can be traced to five predominantly enriched subnetworks. Fig. 2b depicts the overall network structure of the putative indirect DEK1 targets (color-coded by deregulation pattern) including direct upstream TFs in these five subnetworks (color-coded and arranged by subnetwork).

### Transition from 2D to 3D development is encoded by the DEK1-controlled subnetworks

The subnetworks were functionally characterized via network enrichment analysis (Fig.SF3a) using the developmental stages from the *P. patens* Gene Atlas data (Perroud et al. 2018), PO and GO enrichment analysis (Figs. SF3b-i and Table ST3) as well as manual assessment of experimentally characterized *P. patens* genes (Table ST4). These analyses suggest that subnetworks V (chloronema), II (caulonema, rhizoids, side branch initiation and bud formation) and X (vegetative gametophore shoot and leaf cells) encode the cell types, tissues and developmental transitions most relevant to decipher the *DEK1* phenotypes and the 2D to 3D transition (Figs. 2b-d). Subnetwork IX encodes housekeeping gene functions including primary gene regulation, transcription, translation, constitutive epigenetic regulation as well as light-independent mitochondrial and cytosolic metabolic pathways (Figs.2 b,e). Subnetwork VIII harbours the light-dependent and -responsive pathways - in particular photosynthesis, plastid-morphogenesis/ regulation and generally plastid-localized pathways (Figs.2 b,e).

### Network structure suggests DEK1-gated sequential transition between cell fates

The majority of regulatory interactions are found within subnetworks (Fig. SF2a). However, as evident from the enrichment of inter-subnetwork connections (Fig. SF2a), the putative indirect DEK1 targets are predicted to be also controlled by regulators from other subnetworks. In order to determine whether the respective TFs act as positive (activators) or negative (repressors) regulators of potential DEK1-controlled gene functions, we studied the directionality of the inter-subnetwork connections based on the sign of the expression profiles’ correlation coefficients (black [**+**] vs. red [**-**] colored edges in Fig. 2b). Subsequently, we compared the relative proportions of intra- and inter-connections among the enriched subnetworks split according to positive and negative interactions (Figs. 2b,f and SF4).

Strikingly, we found more than expected negative links between V ⊣ II and X ⊣ V (i.e. TFs from V repressing targets in II and X TFs as repressors of V targets) and more positive, activating regulatory interactions between II → X and X → II (i.e. TFs from II activate genes in X and vice versa; Fig. 2f). In our interpretation, this chained pattern potentially reflects the developmental transition between different cell fate identities including primary filament cells differentiating to side branches and gametophore buds. Furthermore, we also observe a biased distribution of *activator* and *repressor* targets among the subnetworks (Fig. 2a). While subnetwork IX is equally enriched for both patterns, V harbors more *repressor targets* and II, VIII and X harbor more *activator targets*. It therefore seems that the enriched subnetworks respond in a specific fashion to the mutation of the plant calpain *DEK1*.

Taken together, the preferential developmental timing and the biased directionality chain suggest the presence of an inherent directionality of DEK1 action on the regulatory circuitry of these subnetworks. These findings may hint towards a mechanism in which DEK1 affects distinct cellular identities as displayed in the *DEK1* mutants.

### DEK1 is part of the APB-controlled subnetwork II guarding the 2D-to-3D transition

The directed edges of the GRN graph can be utilized to reconstruct a regulatory hierarchy by ranking the regulators according to network centrality measures like the fraction of a subnetwork that is directly or indirectly downstream and reachable from a given regulator. Applying this *local reaching centrality criterion*, our analysis identifies the AP2/ERF ANT subfamily transcription factor APB4 as the master regulator at the top of the regulatory hierarchy in subnetwork II (rank 1; reaching > 99% of the subnetwork). The eponymous flowering plant members of the ANT subfamily AINTEGUMENTA, PLETHORA and BABY BOOM (APB) are involved in various developmental processes including the formation of the stem cell niche in the Arabidopsis shoot apical meristem (Hata and Kyozuka, 2021; Scheres and Krizek, 2018; Horstman et al., 2014). Consistent with the proposed role as master regulators of gametophore apical stem cell formation, timed tissue and cell-type specific expression patterns and the additive, but distinct phenotypic severity of single, double, triple and quadruple knockout mutants of the four moss *APBs* (Aoyama et al., 2012), the outparalogous copies *APB2* (rank 35) and *APB3* (rank 41) are localized downstream of APB4 in the regulatory hierarchy of subnetwork II. The inparalogous *APB1* is localized in subnetwork VII, but is also an indirect target of APB4 (Fig. SF5a). *DEK1* is predicted to be localized downstream of APB2 and APB1 in subnetwork II (Fig. SF5b).

The upstream regulatory context suggests that *DEK1* is positively regulated by subnetwork V TFs, activated early in development and subsequently negatively controlled by an auxin/ent-kaurene responsive cascade encoded by subnetwork II as well as a hierarchical cascade of SBP TFs in subnetwork X which already has been shown to be involved in bud formation (Cho et al., 2012). In light of the biased directionality chain identified in the DEK1-controlled inter-subnetwork connections (Fig. 2f), this might represent a negative feedback loop buffering the 2D-to-3D transition at the transcriptional level.

### Deregulation of GRNs is consistent with DEK1 role as a post-translational regulator

Employing the network structure to trace the upstream regulatory contexts of each target gene in the five DEK1-controlled subnetworks, we further evaluated the validity of the *dual role scenario*. Mammalian calpains direct proteins towards the Arg/N-End Rule Degradatory (NERD) pathway (Piatkov et al., 2014). Thus, the potential direct targets of DEK1 should bear N-terminal amino acid residues marking them for ubiquitylation and subsequent degradation by the proteasome (Fig. 3a). The existence and activity of the NERD pathway has recently been demonstrated in *P. patens* and mutants of a key component are also arrested at the 2D-to-3D transition (Hoernstein et al., 2016; Schuessele et al., 2016). Assuming that DEK1 is acting as a post-translational regulator controlling the half-life of transcriptional regulators via the NERD pathway, the majority of deregulated genes in the mutants are expected to be downstream of TFs that are either also deregulated or direct targets of DEK1 calpain cleavage.

**Figure 3.**
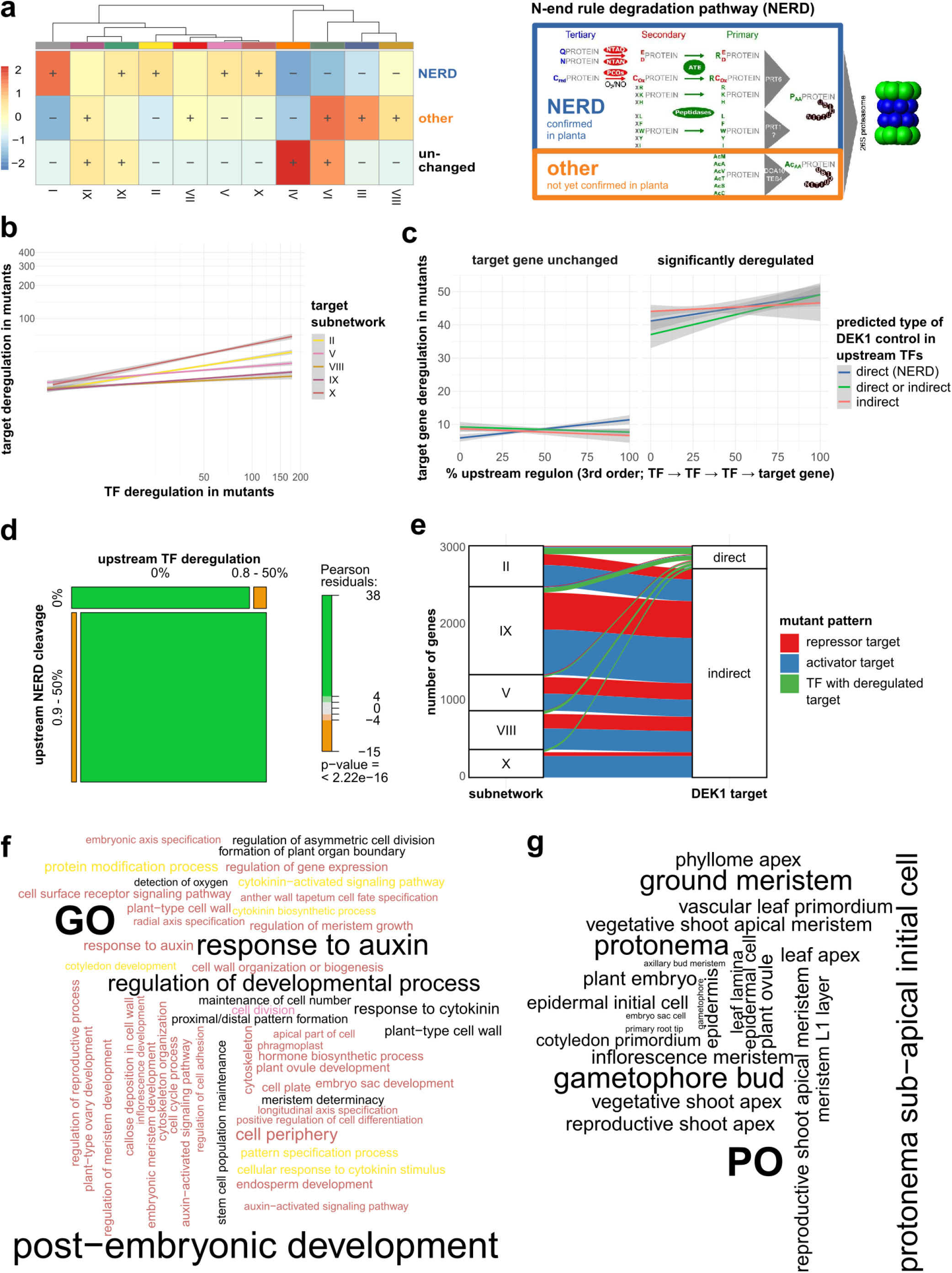
Mutation of DEK1 causes global deregulation of the moss GRN consistent with the pleiotropic phenotype and a post-translational role of the plant calpain in the regulation of protein stability affecting TF↔target interactions. **a.** Network Enrichment Analysis (NEAT) of the prevalence of three specific N-terminal amino acid signature types in predicted calpain cleavage sites of encoded moss proteins. Two of these signature types have been identified in mammals to activate the N-end rule degradation pathway (NERD) resulting in ubiquitylation and subsequent degradation by the 26S proteasome (extended Figure legend on the right adapted from Hoernstein et al., 2016). While the first route has been confirmed to be active in the moss and other plants (<u>NERD</u>), the second route (<u>other</u>) via acetylation of N-terminal residues has not yet been demonstrated *in planta*. The third class represents proteins with no cleavages or sites with N-terminal residues that would not attract the NERD pathway (<u>unchanged</u>). The heatmap depicts the ratio of observed vs expected sizes of specific candidate gene sets encoding for proteins enriched for these types of cleavages among the identified subnetworks. Significant (FDR < 0.01) enrichment (+) or depletion (-) is presented by respective cell annotations. Ratios were clustered for rows and columns using the `ward.D2` method. **b.** Significant <u>target gene deregulation</u> in *DEK1* mutants is positively dependent on deregulation of direct upstream TFs. Linear relationship of target gene and TF deregulation in *DEK1* mutants in the five most affected subnetworks. Deregulation of both types of genes is again depicted as the cumulative effect size of the LRTs in each gene. Lines depict the result of generalized linear regression of the cumulative deregulation of TFs (x-axis) and their target genes (y-axis) for each of the five subnetworks. Grey areas depict 95% confidence intervals. **c.** Target gene deregulation displays positive, linear dependency relative to the fraction of <u>directly and indirectly DEK1 calpain-controlled upstream TFs</u>. Linear regression analysis of the cumulative deregulation of target genes (y-axis; sum of LRT effect sizes) and the percentage of the upstream TFs for each gene where TFs are either directly NERD-targeted by DEK1 (blue line; i.e. classified as NERD-type cleavage Panel 3a, indirectly DEK1 targeted (orange line; i.e. significantly deregulated; contained in gene sets displayed in Panel 3b and Fig. 1f) or either of the two types (green line) for unchanged (left plot) and significantly deregulated (right plot; FDR < 0.1) target genes. Upstream regulon for each target gene in subnetworks II, V and X was evaluated up to 3rd order relationships. Grey areas depict 95% confidence intervals. **d.** Consistent, DEK1 calpain-dependent deregulation in the three subnetworks implementing the 2D-to-3D transition: The deregulated genes in subnetworks II, V and X display significant enrichment of putative NERD-type calpain cleavages and deregulation of upstream TFs. Mosaic plot depicting the relative proportions of significantly deregulated genes in *DEK1* mutants depending on the binary status of their upstream regulon with respect to predicted levels of DEK1 control (x-axis: predicted <u>NERD-type calpain cleavages</u> i.e. direct DEK1 targets; y-axis <u>indirect DEK1 targets</u>). Binary status defines whether the regulon comprises TFs predicted direct (x) or indirect (y) DEK1 targets (> 0% of the TFs) or not (= 0% TFs). Boxes are colored based on Pearson residuals from a significant χ^2^ test of the cross-table comparing the proportions of both binary classes. **e.** Alluvial plot depicting the distribution of the filtered, predicted direct and indirect <u>DEK1 targets</u> among the five predominantly controlled <u>subnetworks</u>. Color-coding of bands reflects directionality of <u>deregulation patterns in the mutant lines</u> (see Fig. 1f for details). The green bands represent unaffected upstream TFs predicted to control the significantly deregulated target genes. **f.** Significantly enriched <u>Gene Ontology (GO)</u> terms associated with direct and indirect <u>DEK1 target genes</u> are overrepresented with concepts related to observed DEK1 phenotypes in flowering plants and the moss. Word cloud depicting filtered, significantly enriched Gene Ontology (GO) *biological process* and *cellular component* concepts (FDR < 0.1). Terms were filtered for overall enriched terms among targets in subnetwork II, V, and X. Furthermore, we curated terms for connection to DEK1 phenotypes. Text size scaled by number of genes per GO term and gene set. Results combined for overall enrichment among DEK1 targets and subnetwork interconnection sets. Color code depicts subnetwork identity of (indirect) target gene. Black indicates overall enrichment among target genes. **g.** Overrepresented tissue and cell type localizations consistent with DEK1 phenotypes and expression patterns. Based on enrichment analysis using <u>Plant Ontology (PO)</u> term annotations for moss genes or their flowering plant orthologs. Word cloud of selected *plant anatomical entity* PO terms displaying an overall enrichment among direct and indirect DEK1 target genes (FDR < 0.1). Filtering and selection of terms was carried out as described for GO terms in Panel 3f.

To test this, we predicted calpain cleavage sites using GPS-CCD (Liu et al., 2011) and subsequently classified the respective proteins based on overall abundance of putative DEK1 cleavages and prevalence of NERD signatures in the resulting, novel N-termini (Figs. SF6a-f). Strikingly, the three DEK1-controlled subnetworks encoding the 2D-to-3D transition (V→ II→ X) are among the five subnetworks that are enriched for such NERD-like calpain cleavage patterns (Fig. 3a). These three subnetworks also display the highest levels of overall gene deregulation among the five DEK1-controlled subnetworks (X<II<V<IX<VIII; Fig. SF6g). Targets of deregulated TFs are more likely to be deregulated than genes downstream of non-deregulated TFs. While all five subnetworks display significant, positive correlation of deregulation levels between target genes and their direct upstream TFs (Kendall’s rank and Pearson correlation tests; p < 2e-16), the individual trends for the subnetworks mirror that of the overall gene deregulation levels and confirm the notion that subnetworks X, II and V are most affected by mutation of *DEK1* (Fig. 3b). Targets of potentially DEK1/NERD-controlled TFs are consistently more deregulated. As the three subnetworks are also enriched for NERD-like calpain cleavages, we investigated the dependency of these target gene deregulation patterns on putative DEK1 cleavages in their upstream regulons. Indeed, the deregulation levels of the indirect DEK1 target genes in subnetworks II, V and X are positively correlated with the percentage of putative, direct DEK1-targets among their upstream TF cascades (Fig. 3c). The upstream regulons of deregulated genes are significantly enriched for TFs that are indirect and direct targets of DEK1 (87%; Pearson residuals >> 4; χ^2^ test p < 2e-16 Fig. 3d). The regulatory cascades demonstrate consistent deregulation patterns. The direct upstream regulons (first order: TF→target) are mostly deregulated themselves. I.e. they are either indirect DEK1 targets because their upstream TF is controlled by DEK1 (Fig. 3d left lower box) or are both direct and indirect DEK1 targets. These TFs are directly cleaved by DEK1 and deregulated in the mutants because an upstream, higher order TF is a direct DEK1 target (Fig. 3d lower right box). Consistently, second (TF→TF→target) and third (TF→TF→TF→target) order regulons of deregulated genes are enriched for predicted direct cleavage by the calpain (Fig. SF6h).

Filtering of the DEK1-controlled regulons to first order interactions results in 531 TFs that are predicted to be direct calpain cleavage targets (Figs. 3e and SF7a). These TFs are predicted to directly regulate the expression of 3,679 significantly deregulated target genes, i.e. genes that are under indirect control of DEK1 (Table ST6). 85 TFs fall in both categories, i.e. are potential direct and indirect DEK1 targets. These predicted first order DEK1 targets form a highly interconnected network, comprising a total of 10,120 network edges (Fig. SF7b). The majority of these genes (74%) are encoded by the five major DEK1-controlled subnetworks (Figs. 3e and SF7a). 73% of the inherent 4,082 inter-subnetwork connections target one of the five subnetworks (Fig. SF7c). More than half of these target genes in the three subnetworks are involved in the 2D-to-3D transition (V, II and X). This is also confirmed by results of an ontology analysis that suggests clear functional delineation of biological processes implemented by the DEK1-controlled repressive and activating intra- and inter-subnetwork regulatory interactions (Figs. 3f-g and SF8a-e).

Taken together, by tracing the upstream regulatory cascades of deregulated target genes, we observed consistent TF-target deregulation patterns and found an enrichment of predicted calpain cleavages at the starting points of such deregulated cascades. These findings are in line with the *dual role scenario*, i.e. support the role of DEK1 as a post-translational regulator controlling the half lives of many genes via the regulatory layers especially in subnetworks II, V and X and thereby gating the sequential transition between the encoded cell fates. Further functional characterization of the predicted first order DEK1 target genes should therefore shed light on the molecular factors underlying the pleiotropic DEK1 phenotype and role in plant development.

### Candidate targets suggest deep conservation of DEK1 control over plant development

Target genes from subnetworks X, II and V positioned downstream of DEK1-controlled activators are enriched for biological processes, cellular components and plant anatomical entities (color text; Figs. 3e, f, g and SF8a) which can be directly linked to the observed *DEK1* mutant phenotypes and tissue and cell-type specific expression profiles of *DEK1* in flowering plants and the moss (Tian et al. 2007; Perroud et al., 2014; Demko et al., 2014; Amanda et al., 2016; Johansen et al., 2016; Perroud et al., 2020).

The predicted DEK1-controlled genes regulate or comprise components determining cell polarity, cell axis, cell number, division, division plane and cell fate. These are involved in biological processes including *regulation of asymmetric cell division* (e.g. by *STRUBBELIG* orthologs; Chevalier et al., 2005), *callose deposition in cell walls* (Panteris et al., 2021) and defined cellular components including the *phragmoplast* (e.g. orthologs of *AUG6*, Lee et al., 2017 and *TANGLED1*, Martinez et al., 2020) and the *cell plate* (e.g. *STRUBBELIG* orthologs or *CLAVATA1*/*PpCLV1b*, Whitewoods et al., 2018).

Besides these specific processes and compartments, the predominant pattern is consistent with the role of DEK1 as regulator governing developmental processes. *DEK1* mutants are disturbed in processes involving general cell fate determination or transition and stem cell, meristem or primordium identity and initiation (e.g. shoot, flower, root and axillary bud meristems; development of endosperm, ovule and embryonic meristems; Fig. 3f). Also, enrichment of anatomical entities among the DEK1 target genes from subnetworks X, II and V include components expected to be affected in *DEK1* mutants (Fig. 3g). These include cell types such as the *leaf lamina* (Hibara et al., 2009), *epidermal initial cell* (Galletti et al., 2015) or the *meristem L1 layer* (Olsen et al., 2015). The latter provides two striking examples where orthologs of the predicted, indirect DEK1 targets have been confirmed to be also deregulated in flowering plant *DEK1* mutants (*HD-Zip-IV TF MERISTEM LAYER 1* Galetti et al., 2015; *CLV3* Liang et al., 2015).

Experimental evidence from *P. patens* contained in Plant Ontology annotations, points to an enrichment of the cell types *protonema sub-apical initial cell* and *gametophore bud*. Both are the two major, sequentially occuring cell types related to the 2D-to-3D transition (i.e. gametophore bud formation). The former is the site for the positioning and initiation of a new apical stem cell inside a growing, primary filament that can either develop into a secondary protonemal side branch or give rise to a three-faced gametophore bud initial, which is the apical stem cell developing into the 3D gametophore. Both cell types have been extensively studied and have been genetically linked to central, conserved developmental regulators acting in both flowering plants and the moss, like e.g. the *APB* and *CLAVATA* genes (Aoyama et al., 2012; Whitewoods et al., 2018). This supports a deep evolutionary conservation of the underlying developmental processes that seem to be controlled by DEK1.

### Tracing the deregulated molecular actors of the complex, pleiotropic DEK1 phenotype in the GRN

To understand the role of the predicted DEK1 targets in these conserved developmental processes and their role in the pleiotropic *DEK1* phenotype, we established a protocol to predict genes underlying specific phenotypic characteristics of the *DEK1* mutant strains. The developed Factorial Differential Gene Expression Network Enrichment Analysis (FDGENEA) method utilizes phenotypic traits encoded as binary factors (Fig. SF10a) for differential gene expression analysis. The resulting differentially expressed genes displaying significantly altered transcript abundances in association with one of the 17 phenotypic factors (Table ST8), were then traced in the GRN to identify the predominantly affected subnetworks (Fig. SF10b,e).

Again, the observed, significant network associations demonstrate the importance of subnetworks X, II and V to explain the *DEK1* phenotype (FDR < 0.01; Fig. SF10b). The resulting FDGENEA sets overlap, but also display substantial portions of genes that are specifically deregulated in response to a single trait (Fig. SF10f). The assessed phenotypic traits clearly separate into two classes that are enriched for either DEK1-controlled *activator* or *repressor targets* (Fig. SF10g; comprising 2,048 indirect DEK1 targets).

### Ectopic gametophore initiation in the mutants can be traced to the deregulation of DEK1-controlled genes with bud cell-specific expression profiles in subnetworks II and X

The earliest step of the 2D-to-3D transition that appears to be disrupted in the *DEK1* mutants, is the initiation of gametophore apical stem cells along the protonema filaments. Phenotypically, this is particularly pronounced in the number of buds per filament and the percentage of filaments with buds (Fig. 1c). While the *oex1* line forms fewer, the *Δdek1* and *Δloop* lines develop significantly more buds per filament than the wild type (ANOVA with post-hoc LSD, p < 0.05). This latter *overbudding* phenotype (Fig. 4a) is consistent with a disrupted control of gametophore initiation, leading to ectopic formation of 3D apical stem cells. It shows the largest unique set of deregulated genes in the FDGENEA of this group of traits (Figs. SF10h; Table ST8) and is enriched for genes from subnetworks II, V, IX, X and XI (Figs. 4a, b and SF10c, e).

**Figure 4.**
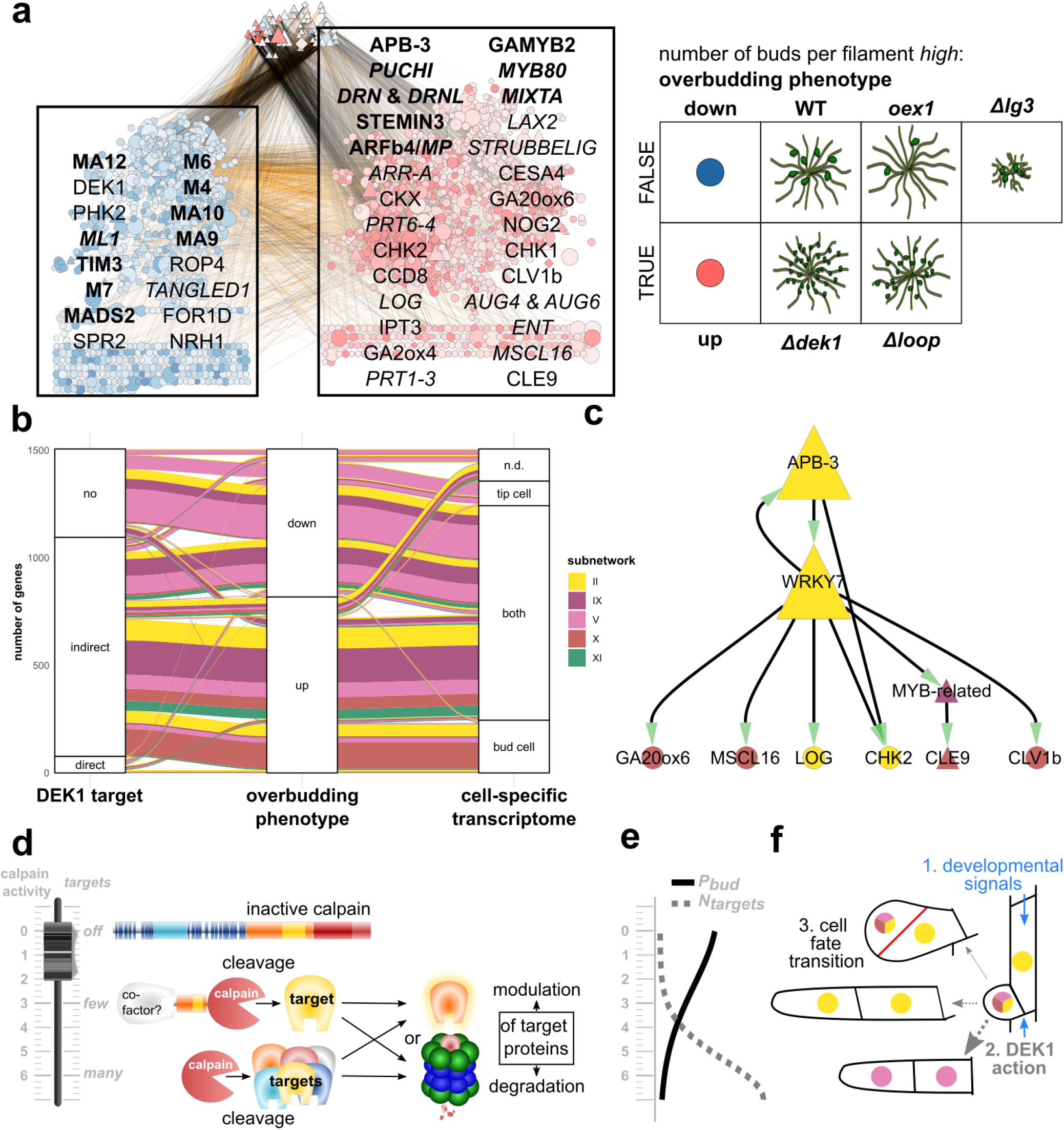
Tracing the mutants’ *overbudding* phenotype to deeply conserved, DEK1-guarded, meristematic regulons controlling the 2D-to-3D transition. **a.** Factorial Differential Gene Expression Network Enrichment Analysis (FDGENEA) of the *overbudding* phenotype reveals enriched subnetworks and upstream regulators associated with high number of buds per filament that comprise key actors in plant meristematic and primordial cell fate control. Network plot of genes with significant association to *overbudding* (left and right node groups in background of overlaid text boxes) and direct, upstream regulators without significant association (top node group). Foreground text boxes display exemplary, predicted DEK1 targets from the *overbudding* up-(right) and down-regulated (left) gene sets with experimental evidence in flowering plants or the moss. Bold font indicates TFs. Italics is used for names of *Arabidopsis* orthologs. Nodes are color-coded by the kind and strength of a gene’s association with the *overbudding* trait (colour intensity gradient relative to log-fold change in DGE analysis; <u>up</u> = positive = red; <u>down</u> = negative = blue). Node sizes relative to the cumulative, absolute deregulation fold-change of the respective gene and any predicted downstream target gene in the mutants. Node shapes: triangles-TFs, diamonds-TRs, inverted triangles-miRNAs, circles-targets. Edge color and intensity: correlation coefficient of connected nodes in DEK1 RNASeq data (black = positive; orange = negative). Panel on the right depicts used genotypes and respective phenotypic character state of the *overbudding* trait (number of buds per filament *high*: FALSE ⇔ TRUE) in wild type and *DEK1* mutant lines. **b.** *Overbudding* up-regulated DEK1 targets from subnetworks II and X are enriched in the previously identified bud cell transcriptome (Frank and Scanlon 2015). Alluvial diagram depicting the proportional distribution of subnetworks shared between three categorical sets (from left to right): <u>DEK1 target</u>: predicted direct and indirect DEK1 targets; <u>overbudding phenotype</u>: genes with significant association with the *overbudding* phenotype (up⇔ down); <u>cell-specific transcriptomics</u> : <u>n.d.</u> not detected; specific to protonemal <u>tip cell</u>; detected but no significant difference between <u>both</u> cell types; specific to gametophore <u>bud cell</u>. Band colouring is based on subnetworks **c.** Key components of the moss CLAVATA peptide (CLE9) and receptor-like kinase (CLV1b) pathway are predicted to be downstream of an *overbudding* up-regulated, DEK1-controlled regulon that comprises a 2D-to-3D master regulator (APB-3) and encodes the integration of several developmental signals (e.g. gibberellin/kaurene (GA20ox6), cytokinin (LOG, CHK2), mechanical stress (MSCL16), peptide (CLE9). Network graph depicting the immediate regulatory context of CLV1b. Full regulatory context is shown in Fig. SF10i. Node sizes are relative to the overall local reaching centrality (fraction of downstream nodes in the global network). Node colouring based on subnetwork affiliation. Triangular nodes are predicted to represent direct cleavage targets of the DEK1 calpain. All predicted regulatory interactions are positive i.e. show positive correlation in wild type and mutants along the RNASeq time course. **d.-f.** Proposed role of DEK1 as a fine-tunable, developmental switch gatekeeping cell fate transitions. Model drawings depicting the proposed relationship between the level of free calpain activity, the number of direct and indirect targets and the developmental consequences in three panels with a shared y-axis (calpain activity). **d.** Model describing three primary DEK1 calpain states (top to bottom): <u>off:</u> immobile, *inactive calpain* in full-length DEK1 protein in plasma membrane. <u>few:</u> mobile, *constrained calpain –* released by auto-catalytic cleavage at several possible locations in the Linker-LG3 domain (Supplementary File). Level of calpain activity, localization, half life and number/kind of targets might be dependent on co-factor interaction. <u>many</u>: mobile, *unconstrained calpain* - pure calpain released by auto-catalytic cleavage directly before or in the CysPc domain. Highest level of calpain activity with a presumably short half life of individual calpain molecules. Potential action as a reset switch resulting in turnover of the entire or large parts of the cells’ protein complement. Depending on the exposed N-terminal signatures (Fig. 3a), cleavage of targets by both types of mobile calpain can either result in <u>modulation</u> of the target or <u>degradation</u> by the NERD pathway. **e.** Proposed relationship between the number of DEK1 calpain targets (*N_targets_* = dashed curve) and the probability of a protonemal cell gaining the bud initial cell fate i.e. gametophore apical stem cell (*P_bud_* = solid curve). **f.** Schematic drawing of the cellular fate transitions affected in the *overbudding* phenotype in three steps. 1. After the perception of <u>developmental signals</u> in the primary filament (blue arrows), a subapical divides and the daughter cell dedifferentiates into a pluripotent, side branch initial. 2. The nature and intensity of the developmental signal (e.g. mechanical stress, phytohormone and peptide gradients) modulates the level of <u>DEK1 action</u> i.e. free calpain activity as indicated in Panel 4d. 3. <u>Cell fate transition</u>: The number and kind of DEK1 calpain cleavage targets and impact on protein half live (i.e. NERD) or activation (e.g. CLE peptides) determines one of four outcomes in the following divisions of the side branch initial cell (nuclei are colored by predominant subnetwork): It either differentiates and continues division as a secondary chloronema filament (predominantly subnetwork V; lower two-cells) or a secondary caulonema filament (subnetwork II; middle) or it divides asymmetrically (red cell wall) to form a gametophore bud (top) maintaining a pluripotent gametophore apical stem cell that will give rise to leaf and stem initials (subnetworks V, II, X) and a basal cell (subnetwork II) that will form rhizoid filaments at the base of the gametophore.

The affected parts of the GRN (Figs. 4a and SF10c, d) can be partitioned into three groups of nodes. Two groups correspond to genes that are either positively associated with a high number of buds (*overbudding-up*; i.e. up-regulated in *Δdek1* and *Δloop*; right group in Fig. 4a and SF10c, d) or those that display a negative association (*overbudding-down*; i.e. down-regulated in *Δdek1* and *Δloop*, but up-regulated in WT, *oex1* and *Δlg3*; left group in Fig. 4a and SF10c, d). The third group is composed of their direct upstream regulators without significant change in expression with respect to this phenotype (top group in Fig. 4a and SF10c, d). The clustering reveals also a trend in the type of connections between the first two groups that also harbor negative regulatory interactions (orange edge color; Fig. 4a and SF10c, d). Overall, while subnetwork V dominates the *overbudding-down* group, the *overbudding-up* assemblage is more diverse and consists of subnetworks X, IX and XI (Fig. 4b and SF10c). Subnetwork II is prominent in both groups. Negative inter-subnetwork links predominantly involve nodes between subnetwork V and either II or X. Subnetworks II and X as well as IX and XI seem to act in conjunction, i.e. share many positive edges (Fig. SF10c). These patterns are consistent with the global network structure discussed above (Fig. 2f) and the sequential transition between the encoded cell fates (Fig. 4f) from primary filament cells (V) redifferentiating to pluripotent side branch initials, that give rise to either secondary chloronemal (V) or caulonemal filaments (II) or gametophore buds (X).

The set of genes up-regulated in filaments displaying the *overbudding* phenotype is dominated by indirect and direct DEK1 targets from subnetworks II, IX and X (Fig. 4b). Down-regulated genes are either not targeted by DEK1 or encoded by subnetwork V or IX. There is a significantly larger proportion of DEK1-controlled regulatory interactions for *overbudding* associated genes in subnetworks II and X (Table ST9). Subnetwork V displays more non-DEK1 controlled interactions, most being negatively associated with *overbudding*. While these regulatory interactions likely represent the side-branch initials redifferentiation into secondary chloronema (Fig. 4f; lower row), the *overbudding* up-regulated, predominantly DEK1-controlled interactions in subnetworks II and X likely encode the cell fate transitions required to establish the gametophore apical meristem (buds).

Consistently, the set with positive association to *overbudding* (*up* in Fig 4b) is enriched for DEK1-controlled *activator targets* from subnetwork II and X which have been previously identified to be specific to gametophore bud cells, while down-regulated genes from V are predominant in the protonemal tip cell transcriptome (Frank and Scanlon 2015). *Overbudding*-associated genes from subnetworks IX and XI are more likely to be found in both transcriptomes, hinting at their more ubiquitous expression profiles or housekeeping function. The bud-specific portion of *overbudding*-*up* DEK1 targets reveals 248 genes (Table ST10). The majority (73%) is encoded by subnetworks II (62; 25%) and X (118; 48%). Thus, the DEK1-controlled, *overbudding* up-regulated *activator targets* from subnetworks II and X represent prime suspects to harbour the developmental regulons acting in the cell fate transitions involved in gametophore apical stem cell initiation which is so pivotal to the 2D-to-3D transition.

### Predicted *overbudding* up-regulated DEK1 targets comprise an interconnected regulon with many known molecular actors of meristematic cell fate specification in moss and flowering plants

Network analysis reveals that around 51% of the 901 *overbudding-up* DEK1 *activator targets* form an interconnected regulon (Fig. SF10i). Given the post-translational role of DEK1, a direct cleavage target will not be deregulated in the mutant context unless an upstream regulator is also a direct cleavage target. Thus, a regulon solely inferred based on the *overbudding-up* genes will be an underprediction. Indeed, when we extend the regulatory context of these genes to include the up to five highest ranking DEK1-controlled upstream TFs, 100% of the *overbudding-up* genes are interconnected (Fig. SF10j).

The *overbudding-up* components of the regulon reveals that the majority is controlled by two regulatory circuits from subnetworks II and X (Fig. SF10i). The first circuit is dominated by TFs from the AP2 superfamily in subnetwork II and comprises several well-characterized key players in meristematic cell fate regulation of flowering plants and *P. patens* (*DRN*, *DRNL* and *PUCHI* Chandler and Werr 2017, *STEMIN3* Ishikawa et al. 2019, *APB-3* Aoyama et al., 2012). The second circuit is dominated by subnetwork X encoded, gibberellin-responsive MYB TFs that have been shown to orchestrate reproductive organ development as well as the production of extracellular hydrophobic barriers like the cuticle and sporopollenin of flowering plants and the moss (*GAMYB2* Aya et al., 2011; *MYB80* Xu et al., 2014; *MIXTA* Oshima et al., 2013). While the two circuits are mostly insulated, some of the target genes overlap (17% of *overbudding-up* only- and 35% of the extended regulon; Figs SF10l, m). At the regulatory level, this insulation might be unidirectional in that one of the subnetwork X MIXTA orthologs is predicted to positively regulate a *DRN* and a *DRNL* ortholog in subnetwork II. This might represent a positive feedback mechanism.

Target genes of both circuits are involved in reorientation of the division plane, modulation of the cell wall, the cytoskeleton and the phragmoplast (e.g. Fig. 4a). Moreover, they encompass a notable accumulation of components required for generation, transduction and perception of local queues like mechanical stress (*MSCL16* Wilson et al., 2013) and longer distance, gradient-forming, developmental signals like plant peptide hormones (*CLV1b*, *CLE9* Whitewoods et al., 2018) or the phytohormones auxin, gibberellin, strigolactone and cytokinin. The latter is especially noteworthy. While the other three phytohormone pathways are represented by one or two components involving either transport (*LAX2* Swarup and Bhosale 2019), biosynthesis (*GA2ox4* Miyazaki et al., 2018; *CCD8* Proust et al., 2011) or activation (*GA20ox6* Miyazaki et al., 2018), in the case of cytokinin, all major aspects are covered. The regulon comprises several genes encoding biosynthesis (*IPT3* Lindner et al., 2014), activation (*LOG* Kuroha et al., 2009), degradation (*CKX* Schwartzenberg et al., 2007), transport (*ENT*, *ABCG* Borghi et al., 2015) perception (*CHK1*, *CHK2* Schwartzenberg et al., 2016) and transduction (*ARR* To and Kieber 2008) of cytokinins. In addition, the MYB circuit of the regulon also is predicted to induce cytokinin-responsive genes like *NO GAMETOPHORES 2* (*NOG2* Moody et al., 2021), whose loss-of-function mutant similar to *DEK1* also displays an *overbudding* phenotype. Consistently, the exogenous application of cytokinin (Schwartzenberg et al., 2007) and cytokinin-overproducing mutants (Schulz et al., 2000) also result in an *overbudding* phenotype. These findings clearly demonstrate the conservation of hormonal control in stem cell initiation and cell fate specification in land plants (Lee et al., 2019).

As mentioned above, the regulon also comprises essential components of the CLAVATA (CLV) peptide and receptor-like kinase pathway that has been shown to control cell fates and division planes of land plant apical stem cells (Fletcher et al., 1999; Whitewoods et al., 2018; Hirakawa et al., 2019) via CLV3/EMBRYO SURROUNDING REGION-Related (CLE Yamaguchi et al., 2016) peptide hormones which are perceived and transmitted to downstream signalling cascades via CLV1-type receptor-like kinases (Hazak and Hardtke 2016). Several studies in both Arabidopsis (Johnson et al., 2008) and Physcomitrella (Whitewoods et al., 2018; Moody et al., 2018; Moody et al., 2021) already have identified parallels in mutant phenotypes and expression patterns and have proposed models locating the CLV signalling pathway somewhere downstream of DEK1 and the moss APBs. The predicted *overbudding-up* regulon identified here, now provides us with a robust explanation for these connections. Our predictions indicate that *CLV1b* and *CLE9* are an integral part of the regulon that is downstream of both of the above DEK1-controlled circuits (Figs. SF10i,k) - in particular downstream of APB-3 (Fig 4c). APB-3 is predicted to coordinate its control over these genes with a calmodulin-binding WRKY group II transcription factor (WRK7 Park et al., 2005), that could act to integrate a possible Ca^2+^ signal emerging in response to the swelling of the gametophore initial cell (Tang et al., 2020). Furthermore, our calpain cleavage predictions indicate cleavage sites that would allow the maturation of CLE peptides from their respective preproteins encoded by the *P. patens* genome (Whitewoods et al., 2018, Supplementary File `PpCLEs.ccd.all`). This could represent another potential feedback layer of the regulon in that CLEs are both positively (maturation/activation) and negatively (indirect *activator targets*) controlled by DEK1. The second order regulatory context of CLV1b (Figs 4c and SF10k) suggests that all positional and developmental queues discussed above are co-regulated in one DEK1-controlled *CLAVATA regulon*. Consistent with the findings from two recent studies (Nemec Venza et al., 2021; Cammarata et al., 2021), our data suggests that this DEK1-controlled, cytokinin-mediated pathway governs stem-cell homeostasis acting separately from the cytokinin-independent pathway involving the RECEPTOR-LIKE PROTEIN KINASE2 (RPK2; subnetwork VIII).

The *overbudding-down* part of the GRN is controlled predominantly by MADS box TFs, contains several correctly predicted, negative regulators of bud and gametophore formation (e.g. *DEK1* Demko et al., 2014; *PHK2* Ryo et al., 2018) and is enriched for cytoskeletal components involved with polarized tip growth of protonemal filaments (e.g. *FOR1D* Vidali et al., 2009; *SPR2* Leong et al., 2018). Plant Rho GTPases (ROP) are key regulators of cellular polarization and are involved in several symmetry breaking mechanisms (Eklund et al. 2010; Muñoz-Nortes et al. 2014). Activated ROP binds effector proteins e.g. to initiate remodelling of the cell wall (Cheng et al. 2020) or the cytoskeleton (Muñoz-Nortes et al. 2014). Sometimes they act as transducers for receptor-like kinases (Jose et al., 2020). The *P. patens* ROP4 is localized at the tip of a growing protonema filament and relocalizes prior to protonemal branching to the future site of side branch formation (Cheng et al. 2020). *ROP4* is predicted to be an *overbudding-down* DEK1 *repressor target*. Rho GTPase-dependent signalling by ROPs is tightly controlled at the protein level (Eklund et al. 2010). ROPs are activated by RhoGEFs, while RhoGDIs and RhoGAPs provide independent means of ROP inactivation. Our analysis detected a representative of both ROP-regulator types as *overbudding*-associated, indirect, DEK1-targets with opposing regulatory patterns (DEK1 *activator target*, *overbudding*-*up*, part of the *CLAVATA regulon*: *ROP-GEF*, Pp3c10_9910; DEK1 *repressor targets*, *overbudding*-*down*: *RhoGAP*, Pp3c3_5940 and *RhoGDI*, Pp3c10_19650). This observation is consistent with their proposed antagonistic role in controlling ROP signalling. It provides a compelling example of how DEK1 might post-translationally control asymmetric and other types of formative cell division by remodelling of cell walls and the cytoskeleton.

## Discussion

### The plant calpain DEK1 acts as a post-translational regulator of protein stability and gene expression

In our study we contrasted two scenarios concerning the functional role of the calpain DEK1 in the developmental regulation of plant cell fate transitions. In the first scenario DEK1 solely acts as a modulatory protease that alters the activity of a limited number of specific proteins in and around the cell wall, cytoskeleton and cell division apparatus thereby primarily acting on the physical layers of cell fate transitions. In the second scenario proteolytic cleavage by DEK1 can have both modulatory and destabilizing consequences for a larger set of target proteins in both the physical and the regulatory layers of cell fate transitions. The destabilizing DEK1 cleavages contain signatures targeting the fragments to the NERD pathway and ultimately towards degradation by the proteasome. As this post-translational regulation also comprises TFs and other gene regulators, DEK1 indirectly controls the expression of many target genes.

By tracing the deregulation profiles of *DEK1* null and over-expressor lines in the gene regulatory networks of the moss *P. patens*, we identified at least 3,679 consistently deregulated genes whose expression is controlled by 531 TFs containing such destabilizing calpain cleavage sites. These TFs are thus direct targets of the plant calpain that in turn acts as an indirect regulator of downstream target genes. The observed deregulation is not limited to the target genes but can be traced through the regulatory hierarchy revealing consistent predicted cleavage sites and mutant deregulation levels of TFs including the master regulators of the respective subnetworks. Individual master regulators but also more downstream TFs and many of the target effector genes have already been experimentally tied to specific *DEK1* phenotypes in *P. patens* and flowering plants. Integrating over all observations, we conclude a dual role of DEK1 as a modulatory and destabilizing protease acting on the physical and regulatory layers of cell fate transitions and thereby indirectly controlling many gene functions.

The role in post-translational gene regulation and predicted list of DEK1 targets provides a consistent explanation for the calpain’s essentiality, pleiotropy and broad effects observed in the moss and also many of the reported phenotypes and genetic interactions in other plants and eukaryotes. Considering that these predictions do not take secondary and tertiary structure, and thus site accessibility, into consideration, the raw number of predicted calpain cleavage sites seem like an overestimate. In light of the multiple layers of regulation and presumed preferential cleavage within unstructured, inter-domain regions reported for metazoan calpains (Ono and Sorimachi, 2012), the true number of *in vivo* cleavage sites must be lower also *in planta*.

We based our predictions on the mutant expression profiles in combination with the inferred gene regulatory networks. The stringently filtered, NERD-classified, predicted calpain cleavage sites were utilized for confirmation. Nevertheless, the detected broadness of those filtered cleavage sites, especially in transcription factors and other gene regulators, is consistent with the observed broad transcriptional, functional and phenotypic responses in *P. patens*.

### DEK1 controls developmental cell fate transitions implemented by deeply conserved land plant regulons

Individual examples like ROP signalling or the above described *CLAVATA regulon* may help to bridge the gap between the well-established image of DEK1 as a developmental regulator that is affecting cell fates and division plane reorientation, and the role as a post-translational regulator proposed here. Our analyses suggest the existence of deeply conserved, orthologous, hormone-guided regulons governing land plant meristems and stem cells that are post-translationally controlled by DEK1 acting as a fine-tunable reset switch implementing or guarding the transition between different cellular identities. The documented, high level of conservation again highlights the utility of the model plant *P. patens* to elucidate embryophyte development and stem cell regulation. In case of the *overbudding* phenotype i.e. the initiation of gametophore apical stem cells, DEK1 is a negative regulator of cell fate transitions. It thus acts as a repressor of the 2D-to-3D transition. Combined with the gene regulatory networks and functional predictions, the *DEK1* mutants thus provide a framework to identify and mechanistically characterize the players involved in this crucial developmental transition.

### Model of a fine-tunable, developmental switch gatekeeping cell fate transitions

In our model (Fig. 4d-f), the level of active calpain is tightly controlled and fine-tunable on multiple levels. DEK1 integrates multiple developmental signals (e.g. phytohormones, peptides, light, mechanical stress) and acts as a gatekeeper in the transition between distinct cellular fates (Fig. 4f). The transcriptional profiling of the four distinct *DEK1* mutants enabled us to monitor three extreme points in the distribution of calpain activity i.e. the number of direct and indirect targets (*N_targets_*; *off = Δdek1*, *Δloop*; *few = Δlg3*; *many = oex1*; Figs. 4d,e). With their intermediate phenotypes and deregulation patterns, the two partial deletion lines indicate how the distinct functional regions of the encoded DEK1 protein might be involved in fine-tuning free calpain activity. In our model for gametophore bud formation, the level of calpain activity is proportional to the probability (*P_bud_*; Fig 4e) of a side branch initial developing into a gametophore initial cell (Fig. 4f).

The immobile, inactive, full-length DEK1 protein (*off*; Fig. 4d) resides in the plasma membrane (Tian et al., 2007; Johnson et al., 2008; Perroud et al., 2020) and can potentially be phosphorylated at several sites (Nühse et al., 2004; Nakagami et al., 2010), probably resulting in conformational changes and (de)activation. While animal calpain activity depends on Ca^2+^ binding (Moldoveanu et al., 2002), it is currently unclear to what extent Ca^2+^ activation is required for the DEK1 calpain’s *CysPc-C2L* protease domain (Wang et al., 2003; Tran et al., 2017). The autocatalytic activity of DEK1 (Johnson et al., 2008; Supplementary File `DEK1.jvp`) likely results in a short half-life of the mobile, unconstrained calpain that would target *many* proteins potentially acting as a reset switch of a cell’s protein complement (*many*; Fig. 4d).

However not all potential cleavages bear destabilizing N-terminal residues targeting a protein towards degradation by the proteasome via the NERD pathway (Fig. 3a). Depending on the amino acid signature of the new N-terminus, the resulting polypeptide is either NERD-directed or stable and may represent the activated or mature form of the protein or peptide (e.g. CLEs). This seems also true for DEK1 itself. Our data suggest the existence of at least three stable calpain variants that potentially arise by auto-catalytic cleavage in the Linker-LG3 domain (Supplementary File `DEK1.jvp`). These are similar to the sizes of experimentally confirmed forms in Arabidopsis (Johnson et al., 2008). The varying N-terminal regions resulting from such cleavages might either lead to different half-lifes or enable binding of different cofactors potentially modifying specificity or the calpain’s target range (*few*; Fig. 4d).

We also found components of the NERD pathway (e.g. orthologs to crucial N-recognins PRT1 and PRT6; Fig. 3a; Holdsworth et al., 2020) among the indirect DEK1 targets. These potentially represent yet another regulatory layer, allowing to switch off degradation or fine-tune protein stability and balance post-cleavage protein fates towards the modulator activity i.e. activation or maturation (Fig. 4d).

### Dual role as a modulatory and destabilizing protease

Mechanistically, calpain research so far has largely focused on the role as a non-processive, modulatory protease (Storr et al., 2011; Sorimachi and Ono, 2012; Ono and Sorimachi, 2012). Whereas its destabilizing characteristics, observable e.g. in the coactivation with the ubiquitin-proteasome system (Freitas et al., 2016) and the generation of short-lived substrates for the NERD pathway (Piatkov et al., 2014), have not yet received much attention. At the same time, many of the experimentally characterized calpain targets (Shinkai-Ouchi et al., 2016), especially those with confirmed NERD degrons (Piatkov et al., 2014), are involved in transcriptional or other forms of gene regulation. The functional implications of this have so far been under-investigated.

In plants, it has been difficult to align the observed directionality (activation vs. inactivation of biological functions), impact, pleiotropy and severity of *DEK1* phenotypes with the role as a modulator protease. This changes by extending our view to also include the role as upstream component of the ubiquitin-proteasome system (Varshavsky, 2012) that directs proteins via the route of the NERD pathway. Destabilizing calpain cleavages in TFs effectively can inhibit the expression of all downstream target genes and thus indirectly regulate gene expression. The substantial changes in gene expression observed in the *DEK1* mutant lines are most parsimoniously explained by the model that proposes DEK1 to act as a post-translational regulator by indirectly controlling the half-life of transcriptional regulators. Nevertheless, DEK1 like other calpains remains a non-processive and modulatory protease. Our predictions do hint at the importance of DEK1 in protein maturation and activation (e.g. DEK1 and CLEs). Thus, we propose a duality of outcomes for proteolysis by calpains (Fig. 4d). Our data in *P. patens* indicate that in most cases the final outcome of such cleavages will result in degradation by the proteasome. Given that calpains frequently are characterized as deleterious, aggravating factors of human pathological conditions (Storr et al., 2011; Sorimachi and Ono, 2012) and the ratio of confirmed NERD-targets among calpain substrates (Piatkov et al., 2014), this might also be a viable and resilient hypothesis for mammalian calpains.

Calpains are involved in a vast spectrum of biological processes and are controlled at multiple levels (Storr et al., 2011; Sorimachi and Ono, 2012; Ono and Sorimachi, 2012). On the evolutionary timescale, the emergence of a novel calpain cleavage site can be considered a random event with either a positive outcome (i.e. a functional neo-protein) or more frequently a negative outcome (i.e. a detrimental fragment). Without the tight regulatory constraints, the fuzzy target specificity and the ubiquitous expression of calpains likely would result in a myriad of fragments and neo-proteins. This necessitated mechanisms that favour functional neo-proteins while simultaneously avoiding or controlling deleterious fragments. The proposed dual role as a modulatory and destabilizing protease acting in the modulation of a fraction of protein functions on the one side, while directing the majority of detrimental cleavage fragments towards the NERD pathway on the other, provides the most parsimonious explanation for what we observe in extant organisms. The gene regulatory consequences of the resulting NERD-control over calpain-targeted TFs subsequently allowed to establish this as a regulatory mechanism in form of a post-translational gatekeeper of cell fates. The fact that *DEK1* is found as a single copy gene in most land plants argues for a crucial and dosage-sensitive role of the plant calpain (Zhao et al., 2012; Demko et al., 2014; Harrison J., 2015).

### New approach to genotype-phenotype mapping and the mechanistic exploration of calpains

The systematic and large-scale analysis or discovery of calpain targets so far has been hindered by the limited target specificity, involvement in a broad spectrum of biological processes and complexity of regulatory mechanisms. Piatkov et al. (2014) even go as far as to state: “Natural calpain substrates were identified thus far largely by serendipity, as distinguished from proteome-scale surveys”. Considering the confirmed calpain targeted human gene regulators and the various reports of gene deregulation in metazoan calpain mutants and human pathologies (Miyazaki et al., 2016; Oliver et al., 2018; Tian et al., 2020), a gene regulatory role might also be true for metazoan calpains. The approach chosen here, i.e. tracing calpain mutant or pathological deregulation profiles in GRNs to identify indirect and direct targets, might thus help to elucidate this yet under-explored aspect of calpain biology in general. The developed FDGENEA method can also prove to be invaluable to other sorts of genotype-phenotype mappings. The outcome of the approach is particularly fruitful for calpain research. The obtained genome-wide and unbiased target candidate gene lists serve as valuable starting points to gain further momentum in the mechanistic exploration of this enigmatic and major proteolytic system with important regulatory and developmental implications in all eukaryotes.

## Acknowledgments

The research reported herein was made possible by FRIMEDBIO grant 240343 from the Norwegian Research Council to OAO. The sequencing service was provided by the Norwegian Sequencing Centre (www.sequencing.uio.no), a national technology platform hosted by the University of Oslo and supported by the *Functional Genomics* and *Infrastructure* programs of the Research Council of Norway and the Southeastern Regional Health Authorities. VD was supported by the Slovak Research and Development Agency grant APVV-17-0570.

## Author Contributions

<u>OAO</u> initiated and designed the *P. patens* mutant study and contributed to writing the manuscript. <u>VD</u> carried out the *P. patens* wet lab experiment including microscopy, observations and interpretations, contributed to the phenotypic factor annotations, performed RNA extraction, genotyped the *oex1* line, prepared Figs. 1b-c, SF9 and SF10a and contributed to writing the manuscript. <u>TB</u> performed initial computational analysis of the DEK1 RNA seq data. <u>MM</u> performed QC of the DEK1 RNAseq data and contributed to the initial DEK1 RNA seq analysis. <u>TH</u> contributed to the initial computational analysis of the DEK1 RNAseq data. <u>PFP</u> contributed to the initial DEK1 RNAseq data analysis, created the *oex1* line and contributed to the phenotypic factor annotations. <u>KFXM</u> contributed to writing the manuscript. <u>WJ</u> and <u>AEA</u> contributed to molecular characterization of the *oex1* line and preparation of RNA samples used for the RNAseq analysis. <u>DL</u> conceived, designed, implemented and carried out all presented data analyses (final DEK1 RNA and global RNA seq analyses, GRN inference and analysis, calpain target predictions, phylogenomics, ontology and other functional annotations, enrichment analyses, FDGENEA…), postulated the DEK1-NERD hypothesis, wrote software, created and curated data sets, prepared the figures and wrote the manuscript.

## STAR Methods

### RESOURCE AVAILABILITY

#### Lead contact

Further information and requests for resources and reagents should be directed to and will be fulfilled by the lead contact, Daniel Lang.

### Materials availability

#### Data and code availability

- *Physcomitrium patens* line (*oex1*) generated in this study as well as other *P. patens* lines used are deposited at Comenius University in Bratislava, Department of Plant Physiology moss collection and are listed in the key resources table.
- RNASeq data have been deposited at ENA and are publicly available as of the date of publication. Accession numbers are listed in the key resources table.
- All generated data sets have been deposited at Zenodo and are publicly available as of the date of publication. DOIs are listed in the key resources table.
- This paper analyzes existing, publicly available data. A table listing accession numbers for the datasets are listed in the key resources table.
- Raw images generated in this study, including microscopy images, gel and gel blot images are publicly available as part of the Zenodo archive listed in the key resources table.
- All 27 *P. patens* gene sets used in the figures or the text are provided as geneid lists in plain text files in the `gene_sets/` folder of the Zenodo archive listed in the key resources table.
- All original code was committed to git repositories. Parallelized Snakemake (Köster and Rahmann, 2012) workflows are provided as individual repositories. Data analyses, statistics and visualizations were implemented via R or Python Jupyter Notebooks (Kluyver et al., 2016) and for convenience are also accessible via the GitHub repository (https://github.com/dandaman/moss_DEK1_GRN_analysis). All git repositories have been pushed to GitHub, deposited at Zenodo and are publicly available as of the date of publication. DOIs are listed in the key resources table.
- Postgresql table dumps as well as additional TSV/CSV tables that are not explicitly mentioned in the below text but are used in the Jupyter notebooks, are provided in the Zenodo archive listed in the key resources table.
- Used packaged software is provided via conda environments included in the Zenodo archive listed in the key resources table. File names of the environments correspond to the Jupyter kernels of each notebook.
- Any additional information required to reanalyze the data reported in this paper is available from the lead contact upon request.

### EXPERIMENTAL MODEL AND SUBJECT DETAILS

#### Physcomitrium patens

*Physcomitrium* (Physcomitrella) *patens* Gransden wild type strain and four mutants *Δdek1* (Perroud et al., 2014); *Δloop* (Demko et al., 2014); *Δlg3* (Johansen et al., 2016) and *oex1* (this work) were used. All five strains were grown in parallel in the same cultivation room. Starting cultures of protonemata were maintained on minimal media supplemented with 920 mg L^-1^ of ammonium tartrate (BCDA) at 16-h light [70–80 mmol m^-2^ s^-1^]/8-h dark at 25°C. Cultures for phenotypic characterization and RNA extraction were grown under the same conditions on minimal BCD media with no ammonium tartrate added (Cove et al. 2009).

### METHOD DETAILS

Supplementary figures and tables represent a selection. The provided Jupyter notebooks comprise all figures and calculation outputs discussed in the below sections.

#### DEK1 protein domain structure

Positions of the 23 transmembrane helices in Fig. 1a (DEK1 MEM, dark blue) inferred by MEMSAT3 (Jones, 2007). Positioning of the remaining domains (DEK1 Linker, DEK1 Calpain) is based on previous phylogenetic analyses (Johansen et al., 2016).

#### Generation of the *DEK1* Linker-Calpain over-expressing strain *oex1*

The Linker-Calpain overexpression vector was built in two steps. First, the cDNA coding for the Linker-Calpain domains was PCR amplified with primer P1 and P2 (Table ST12) using as template the vector containing the *cPpDEK1* full cDNA (Perroud et al. 2014). The fragment was subsequently cloned into the pCR8/GW/TOPO TA vector according to the manufacturer’s instructions (Invitrogen). The Linker-Calpain-coding fragment was subcloned into the pTHUBI-Gateway vector (Perroud et al. 2011) using LR Clonase following standard protocol (Invitrogen) and checked by sequencing for the proper orientation. This vector allows expression during the whole moss life cycle and its targeting to the *108* locus does not induce phenotypic change (e.g. Coudert et al. 2019 or Radin et al. 2021). For transformation purposes the targeting fragment was PCR-amplified using the primer P3 and P4 designed at each end of the targeting sequence.

Wild-type *P. patens* PEG-mediated protoplast transformation procedure was performed (Cove et al. 2009) and the Linker-Calpain over-expressing strain labeled *oex1* was selected for further analysis. In parallel to the molecular analyses (see below), *oex1* went through a cycle of sexual reproduction (Perroud et al. 2011) and displayed normal sporophyte development. Subsequent spore germination and gametophyte development was consistent with the observed phenotype of the original *oex1* transformant (Figure 1). PCR genotyping showed 5’ targeting of the construct at the *108* neutral locus (Schaefer and Zrÿd, 1997). Additionally a Southern blot (Perroud et al., 2014) analysis indicated that *oex1* is a multicopy construct insertion at the locus (Fig. SF11). Genomic DNA for Southern Blot analysis was extracted using the NucleonTM PhytoPureTM Genomic DNA Extraction Kit (GE Healthcare). Southern Blot was performed as described in Perroud and Quatrano (2006) using 1 μg DNA per digestion. Probes were DIG-labelled using the DIG Probe PCR synthesis kit (Roche, Indianapolis, USA) according to the manufacturer’s instructions. pTHUBI-Gateway vector (Perroud et al. 2011) DNA was used as template for PCR amplification of the TS and hygromycin-resistance probes; primers for amplification of the; 5′ TS probe: 108_5fw and 108_5rev; 3′ TS probe: 108_3fw and 108_3rev; HRC probe: HRC-fwd and HRC-rev. The sequences of the primers used to generate the probes are provided in Table ST12. Western blotting using the PpDEK1 specific antibody anti-CysPc-C2L (GenScript, polyclonal produced in rabbit, epitope sequence WSRPEEVLREQGQDC) confirmed the Linker-Calpain protein accumulation. For protein extraction, tissue from 12 days old cultures was homogenized in liquid nitrogen and 300 μg of powder was resuspended in 600 μl of the extraction buffer (0.43% DTT, 6.0% sucrose, 0.3% Na_2_CO_3_, 0.5% SDS, 1.0 mM EDTA, Roche cOmplete Protease Inhibitor Cocktail).

Samples were incubated at 70 °C for 15 min, centrifuged at 2000 rpm for 10 min. Proteins were separated on 4-15% Mini-PROTEAN TGX Gels (Bio-Rad) and transferred on nitrocellulose membranes using Trans-Blot Turbo Transfer Packs (Bio-Rad). Membranes were incubated with primary antibody (anti-CysPc-C2L to detect *P. patens* DEK1 Calpain epitope) diluted 1:500 in TBST buffer supplemented with 5% skimmed milk. Goat Anti-Rabbit IgG (H + L)-HRP Conjugate (Bio-Rad) was used as a secondary antibody and signal was detected using Clarity Max Western ECL Substrate (Bio-Rad) according to the manufacturer protocol.

#### Time series analysis of *P. patens* juvenile gametophyte development

For comparison of wild type and *DEK1* mutant juvenile gametophytic development (Fig. 1b, SF9, SF10a), tissue from one week-old culture of protonemata was homogenized in sterile water and inoculated on minimal medium (BCD) overlaid with cellophane. Material for RNA extraction was harvested after 3, 5, 9, 12, and 14 days of growth, always at the same time of the day. The samples were frozen in liquid nitrogen and stored at −80°C until further processing. Three starting cultures of each strain were used to initiate parallel cultures (biological replicates) used for RNA extraction. Phenotypic characterization of the plant material has been performed using light microscopy and image analysis using ImageJ software.

#### RNA extraction, RNA quality assessment, RNA sequencing of *DEK1* mutants

Total RNA was extracted from frozen material using the RNeasy lipid tissue mini kit (Qiagen) with few modifications. Briefly, the frozen tissue was thoroughly homogenized using a tissuelyser with pre-frozen blocks. Approximately 120 μg of powdered tissue was lysed in 1 mL of QIAzol lysis reagent. Two hundred μL of chloroform was added and the mixture was centrifuged at 4°C. The aqueous phase was collected, 1.5 volumes of 100% (v/v) ethanol was added, and the mixture was vortexed. After binding of the RNA to the RNeasy mini spin column, on-column DNase I treatment was performed to remove genomic DNA. The column was washed with the RPE buffer (Qiagen), dried, and RNA eluted in 45 μL of ribonuclease-free water. The concentration of RNA was measured and RNA integrity was further assessed using an Agilent 2100 Bioanalyzer (DE54704553; Agilent Technologies) with an RNA 6000 LabChip kit. The RNA samples were stored at −80°C until sent for sequencing. Strand-specific TruSeq^TM^ RNAseq library construction of 74 libraries and sequencing using HiSeq2500 as 125 bp paired end reads were performed at *The Norwegian High Throughput Sequencing Centre* in Oslo.

#### Non-redundant gene annotation, phylogenomics framework, regulator classification, improved ontology annotation and updated gene names

For optimal gene-level RNASeq quantification results, a non-redundant transcript representation of the v3.3 cosmoss genome annotation of *P. patens* (Lang et al., 2018) was generated. To this end, GFF3 transcript features of protein-coding and non-protein-coding genes were exported using gffread (Pertea and Pertea, 2020) to FASTA and independently clustered at 100% sequence identity using CD-HIT (Fu et al., 2012). The v3.3 genome annotation contains genes encoding both mRNAs and ncRNAs. As these two transcript types might represent opposite regulatory outcomes (e.g. an antisense transcript to a protein-coding mRNA), they were analyzed independently. The original v3.3 gene ids were extended by adding the primary tag of the transcript feature (i.e. mRNA vs. ncRNA, tRNA, miRNA or rRNA). Resulting transcripts were traced to genes using the original GFF3 parent-child relationships.

Gene families were defined in an automated, phylogenomics approach incorporating protein sequences from 69 Viridiplantae genomes (KRT) using OrthoFinder (Emms and Kelly, 2015). Homologous relationships among gene family members were analyzed by species tree reconciliation of gene trees to infer orthologs, inparalogs and outparalogs. Transcription factors, transcriptional regulators and other transcription associated proteins were inferred based on gene family membership and classification of domain architectures using the TAPScan rule set (Ramírez-González et al., 2018).

Inferred orthologous relationships were used to transfer automatic and experimentally validated annotations from orthologous genes. Gene Ontology (The Gene Ontology Consortium, 2017) and Plant Ontology (Cooper et al., 2013) term annotations were obtained and pooled from Gene Ontology (http://geneontology.org/gene-associations), TAIR (https://www.arabidopsis.org/), and Gramene (ftp://ftp.gramene.org/pub/gramene/release52/data/ontology/) resources. Gene identifiers were mapped to public resources using the UniProtKB mapping table (ftp://ftp.uniprot.org/pub/databases/uniprot/current_release/knowledgebase/idmapping/). The pfam2GO mapping table available from the Gene Ontology resource (http://geneontology.org/external2go/pfam2go) was also employed to transfer GO terms based on the inferred domain architectures. The source evidence classes of the annotated, orthologous genes were translated into target evidence codes of *P. patens* genes as follows: a) automatic annotations: IEA (Inferred by Electronic Annotation) b) experimental and reviewed computational analyses (for full list of evidence codes in these categories see http://www.geneontology.org/page/guide-go-evidence-codes): e.g. EXP (Inferred from Experiment) and e.g. RCA (Reviewed Computational Analysis) and ISO (Inferred by Sequence Orthology) c) pfam2GO: ISM (Inferred from Sequence Model). Subcellular localization predictions using YLOC (Briesemeister et al., 2010), TMHMM (Krogh et al., 2001) and memsat3 (Jones, 2007) that were translated into GO subcellular localization terms. Existing cosmoss *P. patens* v1.6 GO and PO ontology annotations were integrated (Cooper et al., 2013; Zimmer et al., 2013). Altogether, extended annotation comprising 336K GO terms and 877K PO terms was used for the various ontology term enrichment analyses.

Gene names were transferred from the community-curated cosmoss legacy annotations and updated throughout the project to incorporate names from published moss and orthologous plant genes relevant to the study. Final gene names, description lines, regulator and superfamily classifications are provided as part of the Supplementary Files (`genome_annotation/ Physcomitrium_patens.names_and_regulators.tsv`).

#### RNA-seq data collection, read quality analysis and mapping

In total, 299 available RNA-seq libraries were downloaded from EMBL ENA (Harrison et al., 2021) service using the Snakemake workflow searchENA2fastq with the query library_layout=“PAIRED” AND tax_tree(3218) AND instrument_platform=“ILLUMINA” AND (instrument_model!=“Illumina Genome Analyzer” AND instrument_model!=“Illumina Genome Analyzer II” AND instrument_model!=“Illumina Genome Analyzer IIx”). Including the 74 DEK1 RNA-seq libraries produced in this study, 373 libraries were analyzed in total.

Raw data was checked for quality using FastQC (Andrews, 2010) and trimmed in order to remove adapter contamination and poor quality base calls using Trimmomatic (Bolger et al., 2014) with parameters (`ILLUMINACLIP:Illumina-PE.fasta:2:30:10:8:true LEADING:3 TRAILING:3 SLIDINGWINDOW:4:20 MINLEN:60`). After the trimming step, the data was again analyzed with FastQC but also using the `geneBody_coverage.py`, `inner_distance.py`, `RNA_fragment_size.py` and `tin.py` scripts from the RSeQC toolkit (Wang et al., 2012). As a basis for these analyses, trimmed reads were mapped against the *P. patens* V3 genome using HISAT2 (Kim et al., 2019).

Genome-wide alignments of trimmed reads derived from day 14 of gametophyte development in all four *DEK1* mutants and the wild type were created with STAR (Dobin et al., 2013), filtered to the genomic locus of *DEK1* (*P. patens* V3.3 annotation – `Chr17:11873479-11889951`) and displayed (Fig SF11f) using the Integrative Genome Viewer (Robinson et al., 2011).

#### DGE set analyses

Preprocessing, filtering and preliminary analysis of all DGE analyses conducted in this study was implemented in the Jupyter Notebook `dge_analysis/SetAnalysis.factors.ipynb.`Set analysis of DEGs including definition of the *repressor* and *activator targets* sets (Fig. 1f) was conducted using the UpSetR R package (Conway et al., 2017; R Jupyter notebook `dge_analysis/ SetAnalysis4Paper.ipynb`).

### QUANTIFICATION AND STATISTICAL ANALYSIS

#### Quantitative analysis of gametophore meristematic bud formation in *DEK1* mutants

Frequency of gametophore apical stem cell initiation (Fig. 1c) was expressed as the number of buds formed per 15 cell long filament (blue bars; left x-axis) and percentage of filaments forming buds (purple bars; right x-axis). One hundred filaments from each strain were analysed. Statistical significance (a, b, c) of the means was assessed at 95% confidence. Analysis of variance (ANOVA) and least significant difference (LSD) was performed for multiple sample comparisons.

#### RNASeq transcript mapping, gene-level quantification and expression matrix

Paired-end reads were aligned to the set of 80,244 *P. patens* unique transcripts and quantified with Kallisto (Bray et al., 2016) applying 100 bootstrap replicates using the Snakemake workflow workflow_kallisto. Bootstrapped, individual transcript abundances obtained from kallisto were directly used for downstream analysis of differential gene expression (see below). To generate the input expression matrix for GRN analysis, gene-level TPM abundances were calculated using the R package tximport and subsequently normalized using the variance-stabilizing transformation (VST) implemented in the DESeq2 R package (implemented in the Jupyter notebook `grn_analysis/getGeneMatrix. ipynb`).

#### Pairwise, differential time-series gene expression analysis of *DEK1* mutants and the wild type along the developmental time course

Based on the bootstrapped kallisto transcript abundances, we performed pairwise, differential time-series gene expression (DGE) analysis of the *DEK1* mutants and the wild type using the response error linear modeling implemented in the sleuth R package (Pimentel et al., 2017; Jupyter notebooks in folder `dge_analysis/TimeSeriesAnalysis.*_vs_*.ipynb`).

To identify differentially expressed genes (DEGs) between genotypes, we performed pairwise comparisons in the context of the developmental time course. The full linear model thus comprised a variable for the respective genotype pair (e.g. WT vs. *oex1*) and a B-spline basis matrix for a natural cubic spline with four degrees of freedom modelling the data at timepoints 3, 5, 9, 12, and 14 days. The contrasted reduced model only comprised the latter time component. We utilized both the likelihood ratio test (LRT) as well as the Wald’s test of sleuth to test for differential expression. DEGs were inferred using the LRT’s false discovery rate (FDR; *qval* Table ST1) at 10% and 1% confidence. Directionality of differential expression (up-vs. down-regulation; Fig. 1D,f) was defined based on the b-value obtained from Wald’s test.

To identify DEGs during the wild type developmental time course, we selected only the WT samples and performed likelihood ratio and Wald’s testing comparing the B-spline time-series matrix as described above for the full model and the null model as the reduced model (Jupyter notebook `dge_analysis/TimeSeriesAnalysis.WT.ipynb`). FDR cutoff values were chosen accordingly.

To identify DEGs between the early (3-5 days) and the late (9-14 days) phase of wild type development (Fig. 2a), we selected only the WT samples and performed likelihood ratio and Wald’s testing comparing the two phases in the full model versus the null model (Jupyter notebook `dge_analysis/TimeSeriesAnalysis.WT.early_vs_late.ipynb`). FDR cutoff values were chosen accordingly.

To define patterns or profiles of deregulation referred to in the main text as *repressor targets* (deregulation profile 1) and *activator targets* (deregulation profile 2), DEGs were filtered using R Jupyter notebooks `dge_analysis/profile_phases/identify_profiles.ipynb` and `dge_analysis/profile_phases/getRelaxed.ipynb`. For profile 1, we selected genes significantly up-regulated in WT vs. *oex1*, down-regulated in WT vs. *Δdek1* and down-regulated in *oex1* vs. *Δdek1*. For profile 2, we selected genes significantly down-regulated in WT vs. *oex1*, up-regulated in WT vs. *Δdek1* and up-regulated in *oex1* vs. *Δdek1*. For an additional description of the different phases along the time series, each profile was clustered using k-means into three clusters. These clusters were manually interpreted and translated into *phase* descriptions. For profile 1, cluster 1 comprises 943 *late*, cluster 2,958 *early* and cluster 3,738 *uniform*ly regulated genes. For profile 2, cluster 2 comprises 1,019 *late*, cluster 1,667 *early* and cluster 3,759 *uniform*ly regulated genes.

#### Ontology term enrichment in deregulated genes

Ontology term enrichment analyses for the distinct sets of DEGs obtained from the pairwise comparisons of wild type and mutant genotypes (Fig. SF1 and Table ST2), were carried out using the Snakemake workflow `ontology_enrichment_workflow` that builds on the Ontologizer software to test multiple sets in parallel for enrichment of terms in any OBO formatted ontology. Percentages of deregulated ontology terms for each mutant genotype were calculated and drawn in the Jupyter notebook `ontology_enrichment_workflow/PercentDeregulated.ipynb` (Fig. SF1).

#### Prediction and characterization of gene regulatory interactions and subnetwork inference

Regulatory interactions were predicted in the genome-wide VST-transformed expression matrix based on 1,736 regulators using the Random Forest predictor of GENIE3 (Huynh-Thu et al., 2010; R Jupyter notebook `grn_analysis/GENIE3.ipynb`). A set of 992 transcription factors (TFs), 413 transcriptional regulators (TRs), 79 putative transcription-associated proteins (PTs), 275 microRNAs and DEK1 were specified as candidate regulators.

Overall directionality of regulatory interactions was determined by the Pearson correlation coefficient of regulator and target gene expression levels along the developmental time course in the wild type and *DEK1* mutant samples as well as globally using all columns of the matrix (Python Jupyter notebook `grn_analysis/GetCorrelation.ipynb`).

Community detection was carried out using the Parallel Louvain Method implemented in the NetworKit Python package based on the top10 regulatory interactions of each target gene (Staudt et al., 2016; Python Jupyter notebook `grn_analysis/GetCommunities.ipynb`).

To characterize the connectivity of nodes and rank the nodes, several centrality measures were calculated, among them local reaching centrality, degree centrality, betweenness centrality and eigenvector centrality implemented using the NetworKit Python package (Python Jupyter notebook `grn_analysis/GetCommunities.ipynb`). Local reaching centrality of a given node represents the fraction of the subnetwork that is reachable from it by following the outgoing edges. Degree centrality measures the number of edges or connections of a node. Betweenness centrality measures the amount of “traffic” a node handles in relation to the entire network. Eigenvector Centrality is defined as the centrality of a node that is proportional to the sum of the centralities of the nodes it is connected to. These values leading by the local reaching centrality were used to sort and rank the nodes to establish a regulator hierarchy.

Barplot summarizing the subnetwork structure (Fig. SF2a) of the *P. patens* GRN was drawn in the Jupyter notebook `grn_analysis/PlotSubnetworkDeregulationPatterns.ipynb`.

#### Identification of the five DEK1-controlled subnetworks

Identification of predominantly DEK1-controlled subnetworks encoding the 2D-to-3D transition (Figs. 2a and SF2b) was carried out by tracing the over- and under-representation of the relevant DGE sets (`gene_sets/*.set,` names as in heatmap column and row descriptions in Fig. 2a) via a Network Enrichment Analysis Test (NEAT; Signorelli et al., 2016) implemented in the R Jupyter notebook `grn_analysis/PlotSubnetworkDeregulationPatterns.ipynb`. P-values were adjusted for multiple testing using the Benjamini-Hochberg method and filtered at 99% confidence.

The overall network structure of the putative indirect DEK1 targets (Fig. 2b) was analyzed and drawn in Cytoscape (Shannon et al., 2003) applying the AutoAnnotate (Kucera et al., 2016) app on the *TFs with deregulated targets*, *activator* and *repressors target* sets with subsequent manual alignment and color coding based on the subnetwork affiliation (Cytoscape file `grn_analysis/Only.intersect_NEAT_profiles_and_gene_atlas_spore_proton ema_gametophore.cys`).

#### Differential gene expression (DGE) analysis of developmental stages in the *P. patens* Gene Atlas

Based on the bootstrapped kallisto transcript abundances of the *P. patens* Gene Atlas data set (Perroud et al. 2018; `dge_gene_atlas/subset.full_metadata.txt`), we carried out DGE analysis for each of the five covered developmental stages (spores, protonema, gametophores, green sporophytes, brown sporophytes) using the sleuth R package as described above for the developmental time course. In this case, the full model compared samples for each developmental stage against all other samples. Analyses were carried out in the Jupyter notebooks `dge_gene_atlas/DGE.gene_atlas.*.ipynb`.

#### Functional characterization of the subnetworks

We utilized the developmental stage samples included in the Physcomitrella Gene Atlas (Perroud et al. 2018) as well as Plant Ontology (PO) and Gene Ontology (GO) annotations for functional characterization of the subnetworks. Both approaches independently considered the network’s structure to assess overrepresentation of functional concepts among the genes in the network. Combined with the manually curated set of experimentally/genetically characterized moss genes, the two analyses provided the basis for the assignment of subnetworks to tissue and cell types (Figs. 2c-d) and to subcellular compartments (Fig 2e).

Network enrichment analysis for the Gene Atlas developmental stage DGE sets defined at FDR < 0.1 (Fig. SF3a) was carried out using the NEAT R package (Signorelli et al., 2016) in the Jupyter notebook `grn_analysis/NEAT.DGE.ipynb` as described above for the DEK1 DGE sets.

The ontology analysis comprised a multi-step procedure relying on a machine learning approach to identify the most specific and characteristic terms for the genes encoded in each subnetwork. The final set of most characteristic PO and GO terms for each subnetwork (Figs. SF3e-i) comprises the ontology terms that were most informative to classify the top20 master regulators from each subnetwork according to their targets’ functional composition.

Primary ontology term enrichment analysis for the subnetworks was carried out using the `ontology_enrichment_workflow` as described above for the DGE sets. Total number of enriched terms at FDR < 0.1 for each partition of GO and PO in this primary analysis (Fig. SF3b) was analyzed and plotted using the R Jupyter notebook `grn_analysis/StudyEnrichments. ipynb`.

Specificity of the primary analysis was analysed via set analysis of the enriched GO biological process concepts (Fig. SF3c) using the UpsetR package (R Jupyter notebook `grn_analysis/StudyEnrichments.ipynb`).

The sources of ontology term annotations are manifold and differ in quality, resolution and intention.

E.g. genes can be experimentally connected to multiple processes while their direct functional involvement is limited to only some of them. Primary enrichment analysis does not consider the relationships between genes. As functionally related genes tend to be co-regulated, GRNs provide an additional layer to mine functional relationships. Even more so in our case, where we are interested in identifying the predominant biological processes and anatomical structures etc. encoded by each subnetwork. Thus, the subsequent steps were directed to integrate the information from the directed graph structures of the predicted GRN of *P. patens*.

Directed network enrichment analysis tests were carried out for each enriched ontology concept among the subnetworks using the NEAT R package (Signorelli et al., 2016) filtering terms at FDR < 0.01 in Jupyter notebook `grn_analysis/NEAT_enriched_terms.ipynb`.

In a next step, we constructed a regulator matrix where the 2,084 columns contain for each NEAT enriched ontology term the annotated downstream gene frequencies for 1,667 regulators (R Jupyter notebook `grn_analysis/GetRegulatorMatrix.ipynb`).

This matrix was used for a machine learning approach to identify distinctive ontology concepts for the top 20 regulators of each subnetwork using Random Forest classification implemented in the randomForest R package (Liaw and Wiener, 2002; Jupyter R notebook `grn_analysis/enriched_terms_selection_by_RandomForest_variableImporta nce.ipynb`). For preprocessing, the regulator matrix was further filtered to discard non-plant GO concepts as well as terms from either PO or GO that did represent >= 10% of the respective ontology’s annotated gene space in at least one of the subnetworks. Individual regulators’ ontology term gene frequencies of the remaining 379 columns were scaled using the overall number of genes annotated with each term and the terms information content. The rows of the matrix were filtered selecting only the top 20 master regulators for each subnetwork using the centrality rank criterion (220 regulators in total). The resulting, filtered matrix was used to train a Random Forest classifier with 10,0000 trees recording variable importance i.e. each ontology term’s importance to discriminate a subnetworks regulator from those of other subnetworks. Multidimensional scaling (MDS) plot of the classifiers’ proximity matrix was carried out to analyze the conceptual similarity of the subnetworks (Fig. SF3d). The top 5 most specific terms to describe targets in subnetworks II, V, VIII, IX and X were plotted as word cloud representations (Fig. SF3e-i). We selected and ranked terms for each subnetwork demanding variable importance > 0 using the decrease in node impurity based on the Gini index implemented by the R/randomForest package.

To identify and rank the five subnetworks contributions to the seven major subcellular compartments depicted in Fig. 2e, we semi-automatically screened, sorted and ranked the distinctive terms by their gene frequencies in the subnetworks using the bash shell (`ontology_enrichment/syntax. get_DEK1_Fig2_numbers`).

#### Identification of the major inter-connections between the five DEK1-controlled subnetworks

To identify the major regulatory interactions between subnetworks (Fig. 2f), we analyzed the cross-sectional distribution of inter-subnetwork connections considering the predicted directionality based on the Pearson correlation coefficient of expression profiles between a regulator and a predicted target from another subnetwork.

In the Jupyter notebook `grn_analysis/PlotSubnetworkDeregulationPatterns.ipynb` utilized stacked barcharts in polar coordinate plots, a mosaic plot depicting the distribution of Pearson residuals indicating significant over- or under-representation of inter-connecting edges obtained from a significant Χ^2^ test (p-value < 2.22e-16; Fig. SF4) and cross-tabulation via Χ^2^, Fisher and McNemar tests implemented in the gmodels R package (Warnes et al., 2018) to assess the distribution of inter-subnetwork edges. The graph of inter-subnetwork connections shown in Fig. 2f depicts the major, significantly enriched inter-subnetwork connections with Pearson residuals > 4 (Χ^2^ test).

#### Subgraph analysis – regulatory hierarchies, regulons and regulatory contexts

To analyze and visualize regulatory hierarchies, regulons and compare regulatory contexts (Figs. SF5a,b, SF10i,k), we utilized the Cytoscape and the k-shortest paths algorithm with additive edge weights implemented in the PathLinker app (Gil et al., 2017). We used the edge weights computed by GENIE3 (Huynh-Thu et al., 2010) and usually explored several k’s to optimize resolution.

#### Prediction of calpain cleavage sites and classification of potential for NERD targeting

We predicted calpain cleavage sites for all predicted protein isoforms of the *P. patens* V3.3 genome annotation using GPS-CCD (Liu et al., 2011) with high confidence threshold setting and subsequently parsed and classified the result using regular expressions capturing the N-end rule (Fig. 2a; `calpain_cleavage_prediction/n-terminal_site_classes.csv`) using Perl-one-liners (bash syntax file `calpain_cleavage_prediction/syntax`). Individual number of predicted cleavage sites per protein was tabulated overall and for each of the five types of NERD signatures (Fig. 3a; <u>NERD confirmed *in planta*</u>: tertiary_deaminated, tertiary_oxidized_or_acetylated, secondary_ATE and secondary_peptidase; <u>other not yet confirmed *in planta*</u>: primary_acetylated).

Subsequently we classified the respective proteins based on overall abundance of putative DEK1 cleavages and prevalence of NERD signatures in the resulting, novel N-termini using a combination of PCA and model-based clustering implemented in the R packages FactoMineR and mclust (Lê et al., 2008; Scrucca et al., 2016 Figs. SF6a-f; Jupyter R notebook `calpain_cleavage_prediction/ DEK1_cleavage_sites.ipynb`).

Overall site frequency and individual NERD site type frequencies were scaled by the total protein length. Log-transformed, scaled overall site frequencies were clustered with k-means clustering into 5 site abundance level categories (SLC; Fig. SF6a). Subsequently, the entire six column matrix was utilized for PCA (Fig. SF6b). The first ten principal components were used for model-based clustering using default parameters (Fig. SF6c-d). The resulting clusters were interpreted in the context of the first five principal components eigenvectors (∼99% of total variation; Fig. SF6d) and the distribution among the five SLCs (Fig. SF6e-f). In particular, we assessed the proportion of cleavages resulting in NERD-like signatures that have been experimentally confirmed *in planta* (Fig. SF6e-f).

Doing so, we could classify category I proteins falling in SLC = 0 i.e. with no predicted cleavages (Fig. SF6f <u>unchanged</u>; Jupyter notebook data frame attribute `fate_strict==”unchanged”`), category II proteins bearing higher proportion of NERD-like signatures (Fig. SF6f <u>NERD</u>; mclust clusters 1 or 2; uncertainty for cluster 3 < 0.05; `fate_strict==”NERD”`) and category III proteins with other or ambiguous signatures (<u>other</u>; `fate_strict==”other”`). In the subsequent analyses, proteins classified as category II were considered as potential DEK1 calpain targets (if they showed consistent deregulation in the *DEK1* mutants; see below) while categories I and III were not.

#### Tracing deregulation and predicted calpain cleavage patterns in the *P. patens* GRN

Directed network enrichment analysis of the calpain cleavage classifications (Fig. 3a) was performed with the NEAT R package (Signorelli et al., 2016) using the R Jupyter notebook `calpain_cleavage_prediction/NEAT_cleavage.ipynb`.

To study gene-wise impact of *DEK1* mutation, we calculated cumulative deregulation levels of genes with significantly altered expression levels in the mutants as the sum of the three individual χ2 test statistics of the three likelihood ratio tests (LRTs) comparing wild type vs. *Δdek1*, *oex1* vs. wild type and *Δdek1* vs. *oex1* employing it in the sense of an absolute, cumulative effect size.

Together with the calpain cleavage classifications, these cumulative deregulation levels were used to trace and describe their patterns in the five DEK1-controlled subnetworks from multiple angles using several data science approaches including generalized linear modeling, Random Forest classification, PCA, χ2 tests, correlation analysis (Figs. 3b, SF6g-h; R Jupyter notebook `calpain_cleavage_prediction/CCinRegulators.TargetPerspective.only_tar get_subnetworks.ipynb`). As a basis, we analyzed the upstream regulatory context of all genes in the five DEK1-controlled subnetworks by tracing incoming regulatory edges for each gene up to the third order (TF3 → TF2 → TF1 → gene) building on the make_ego_graph function of the igraph R package (Csárdi and Nepusz, 2006). To focus on the DEK1-controlled cell fates, we repeated these analyses for the three subnetworks encoding the 2D-to-3D transition (V → II → X; Figs. 3c-d; R Jupyter notebook `calpain_cleavage_prediction/CcinRegulators.Target Perspective.only_II_V_X.ipynb`).

#### Filtering final set of direct and indirect DEK1 targets

In order to define the final set of direct and indirect DEK1 targets, we analyzed the initial set of 215,189 regulatory interactions from all subnetworks involving TFs as regulators (R Jupyter notebook `calpain_cleavage_prediction/GetCandidateTargets.ipynb`). For both sides of the interaction (TF → target) information on calpain cleavage classification, significance of overall deregulation in *DEK1* mutants and the global *DEK1* mutant deregulation pattern was added. Subsequently the final set of DEK1-controlled regulatory interactions (Figs. 3e and SF7) was defined and analyzed using the R Jupyter notebook `calpain_cleavage_prediction/ filter_analyze_DEK1Targets.ipynb` selecting TF regulators with a classified NERD-like cleavage pattern and significantly deregulated target genes. This resulted in 10,120 network edges (Table ST6). Mosaic plot comparing the different types of DEK1-controlled regulatory interactions across the 11 subnetworks (Fig. SF7c) was created using the R Jupyter notebook `calpain_cleavage_prediction/filter_analyze_DEK1Targets. ipynb`. Alluvial plots depicted in Figs 3e and SF7a were created using R Jupyter notebook `calpain_cleavage_prediction/alluvial.ipynb`.

#### Functional characterization of DEK1 targets

Primary ontology term enrichment analysis for the DEK1 targets was carried out using the `ontology_enrichment_workflow` as described above for the DGE sets. We carried out two types of comparisons looking globally at all targets as well as at individual subnetwork pairings and deregulation patterns (e.g. `activator_II_vs_X` or `repressor_X_V`). The enriched terms at FDR < 0.1 for each partition of GO and PO was analyzed and plotted using the R Jupyter notebook `calpain_cleavage_prediction/ontologies/AnalyseEnrichment.ipynb`.

As already demonstrated for the overall DGE sets, DEK1 mutation results in a deregulation of broad gene functions. Thus, our goal was to identify overall trends without losing specificity. We chose a two-pronged approach, combining automated semantic analysis with manual identification of representative key concepts.

The basis for the automated analysis of enriched ontology terms, is semantic similarity analysis (Gan et al., 2013) and information content- or ontology-based ranking of similar concepts. We used the ontologyX suite of R packages (Greene et al., 2017) in the above-mentioned R Jupyter notebook to compare and cluster entire gene sets (Figs. SF8a-c) as well as compare, group and rank individual enriched terms to select the most informative for plotting their semantic similarity-derived distance matrix (1-S) via multidimensional scaling (Figs. SF8d,e).

The result of this automated analysis was then manually inspected, interpreted and curated in light of the external knowledge of *DEK1* mutant phenotypes in *P. patens* and other plants, calpains as well as the NERD pathway to select the most informative and non-redundant concepts (Figs. 3f,g; R Jupyter notebook `calpain_cleavage_prediction/ontologies/PlotSelectedTerms. ipynb`).

#### Factorial Differential Gene Expression Network Enrichment Analysis (FDGENEA)

As a basis for FDGENEA, phenotypic observations of the *DEK1* mutant lines (Figs. SF9 and SF10a) were translated into 17 binary, factorial variables (TSV file `fdgenea/ phenotypic_factors.tsv`; Figs. 4a, SF10b) and used for DGE analysis using the R sleuth package (R Jupyter notebooks `fdgenea/Analysis.*.ipynb`) as described above for the developmental time course. In this case, the full model compared the samples representing the respective `normal` vs. the respective `alt` state of the trait (e.g. `normal` vs. `high` for `number_buds_per_filament`). Positive association of a gene with a trait corresponds to a Wald’s test b value > 0 and negative association is determined by b < 0. The LRT test statistic is interpreted as an effect size for the strength of the association. The full data for this analysis is provided in the files `fdgenea/dge/*/dge.full.tsv.gz`.

Subsequently, for each trait network enrichment analysis of significantly associated genes (LRT FDR < 0.01) was conducted using the NEAT R package (R Jupyter notebook `fdgenea/NEAT.iypnb`). P-values were adjusted for multiple testing using the Benjamini-Hochberg method and filtered at 99% confidence. We carried out two types of NEAT analyses:

1. testing enrichment of the two directional sets of significantly associated genes independently (e.g. trait `number_buds_per_filament` comparison of `normal` vs. `high` is `up`, i.e. genes with b > 0). Full results of this analysis are provided in TSV file `fdgenea/ NEAT_subnetwork_enrichment.phenotypic_factors.tsv`
2. testing enrichment of the entire set of genes that is significantly associated with a given trait (e.g. trait number_buds_per_filament comparison of normal vs. high, i.e. genes with b < 0 or b > 0). Full results of this analysis are provided in TSV file `fdgenea/ NEAT_subnetwork_enrichment.phenotypic_factors.simplified.tsv`

The results of these two analyses were combined for Fig SF10a, using the second analysis as a basis for the heatmap and the results of the first to derive the cell annotations (`+/-`), to indicate significantly enriched sets of either positively (`up` i.e. +) or/and negatively (`down` i.e `-`) associated genes for any of the subnetworks.

As a final step, we implemented a procedure that analyzed the network structure of the genes associated with each trait, isolated subgraphs with enriched trait association and identified common upstream transcription factors. For each gene the algorithm evaluates the cumulative effect size (i.e. the association of the downstream genes with the trait) and records the numbers of all downstream regulators and only the downstream TFs in both cases distinguishing between direct (1^st^ order downstream genes and indirect (higher order) downstream genes. The cumulative effect size was recorded both as a sum of the LRT test statistic (columns ending on `_cb`) and as the sum of the absolute value of the LRT test statistic (columns ending on `_cab`). The identified connected components were reported as TSV files with all the collected statistics as well as individual GraphML files that can be opened e.g. in Cytoscape. The algorithm was applied to two data sets, one comprising all subnetworks and one comprising only the NEAT enriched (FDR < 0.01) subnetworks for each trait. The algorithm was implemented in R using the igraph package (Multi-threaded R code `fdgenea/FDGENEAE_all.R` and `fdgenea/FDGENEA_only_enriched.R`). An example of the procedure with additional plots and analyses is provided for the *overbudding* trait (`normal` vs. `high` for `number_buds_per_filament`; Fig. 4) in the R Jupyter notebook `fdgenea/ FDGENEA.overbudding_only.ipynb`.

The GraphML of the identified connected components for both data sets for the *overbudding* phenotype (Fig. 4a) as well as subgraphs/regulons of identified key players were analysed and plotted using Cytoscape (Figs. 4a,c and SF10c,d,i-m).

Intersections among the FDGENEA sets for all traits were analyzed in the R Jupyter notebook `fdgenea/Intersections.ipynb` (Figs. SF10e,f).

Intersections among the FDGENEA sets for all traits and the two types of predicted DEK1 targets (*activator* and *repressor targets*) were analyzed in the R Jupyter notebook `fdgenea/DEK1_targets.Intersections.ipynb` (Figs. SF10g,h).

#### Further characterization of *overbudding* genes using cell-type specific transcriptomes

Significant DEGs from the cell-type specific transcriptome data for protonemal tip cells and bud cells (Frank and Scanlon 2015) were mapped to the current genome annotation, intersected with the predicted direct and indirect DEK1 targets as well as the genes associated with the *overbudding* phenotype and plotted as an alluvial plot (Fig. 4b) using the R Jupyter notebook `fdgenea/Cell-type_transcriptomes.Intersections.ipynb`. The partial overlap between DEK1-controlled AP2 and MYB TFs in subnetworks II (Fig. SF10l,m) was analyzed using Cytoscape and intersections were drawn using the VIB Venn web interface (http://bioinformatics.psb.ugent.be/webtools/Venn/).

### Supplemental information titles and legends

#### Supplementary Figures

**Supplementary Figure SF1. Percentage of ontology terms in deregulated genes indicates that DEK1 mutation affects most cellular and organismic functions.** Percentage of unique ontology terms from the Gene Ontology *biological process*, *molecular function*, *cellular component* partitions and the Plant Ontology *plant anatomical entity* and *plant structure development stage* partitions that are assigned to deregulated genes in the mutant lines.

**Supplementary Figure SF2. Tracing DEK1 impact on the gene regulatory networks (GRN) of *P. patens*.**

**SF2a. Summary of the predicted *P. patens* GRN.** Barplot depicts the percentage of total edges (regulatory interactions) and nodes (genes) comprising the 11 subnetworks that form the *P. patens* GRN. Absolute numbers are given as bar stack annotations. Inter-subnetwork edges are displayed in white.

**SF2b. Network enrichment analysis of DEK1-deregulated genes.** Barplot displaying the major patterns of deregulation in *DEK1* mutants (middle and right bar) as well as upstream transcription factors (left bar) predicted to directly target the affected genes across the 11 subnetworks. Numeric annotations in bar stacks are provided for subnetworks with significant (FDR < 0.01; network enrichment analysis) enrichment of genes with a mutant deregulation pattern that is consistent with being a target of a DEK1-regulated activator or repressor (see Fig. 1e and 1f) and depicted in the network graph of Fig. 2b. Individual, significant data points are indicated by an asterisk.

**SF2c. Inter- and intra-connections between enriched subnetworks.** Pie chart depicting the overall percentage of edges within and between the enriched subnetworks (Fig 2b and SF2b) split according to edge directionality inferred by pearson correlation coefficient of normalized expression levels (r > 0 [+]: outer ring; r < 0 [-]: inner ring). Individual subplots display the subnetwork origin of the target while the color coding depicts the subnetwork origin of the respective regulator.

**SF2d. Non-redundant intersections between the different gene sets.** UpsetR plot displaying non-overlapping intersections between different gene sets discussed in the main text and figures (Fig 2 and SF2b). The gene sets are available as individual lists in the supplementary file archive (`gene_sets.zip`).

**Supplementary Figure SF3. Functional characterization of the subnetworks.** We utilized the developmental stage samples included in the *Physcomitrella* Gene Atlas (Perroud et al. 2018) as well as Plant Ontology and Gene Ontology annotations for functional characterization of the subnetworks. Both approaches independently considered the network’s structure to assess overrepresentation of functional concepts among the genes in the network. The ontology analysis comprised a multi-step procedure relying on a machine learning approach to identify the most specific and characteristic terms for the genes encoded in each subnetwork. The final set of most characteristic PO and GO terms for each enriched subnetwork (Panels SF3e-i) comprises the ontology terms that were most informative to classify the top20 master regulators from each subnetwork according to their targets’ functional composition.

**SF3a. Network enrichment analysis of differentially expressed genes in developmental stages from the *Physcomitrella* Expression Atlas.** We inferred up- and down-regulated genes for each developmental stage represented in the Gene Atlas data by performing differential gene expression analysis using a combination of the likelihood ratio test (LRT) and Wald test implemented in R/sleuth comparing the samples derived from the respective stage with all other samples. Results were filtered using LRT-qvalue < 0.1 and classified into up and down-regulation based on b-values of the Wald test. Individual Jupyter notebooks for each of these analyses are provided as part of the supplementary file archive. Individual gene sets were tested for undirected enrichment in the subnetworks using the R/NEAT software package. Results were plotted as ratios of observed/expected values in a heatmap using the R/pheat package (CRAN) annotating significant (FDR < 0.01) enrichment and depletion.

**SF3b. Subnetwork ontology enrichment analysis**. We performed ontology enrichment analysis for the subnetworks based on an updated Plant Ontology (PO) and Gene Ontology (GO) annotations available as part of the supplementary file archive using the command line version of Ontologizer (http://ontologizer.de/) with Benjamini-Hochberg (FDR) correction at 90% confidence. The stacked barchart depicts the absolute numbers of filtered enriched terms.

**SF3c. Non-redundant overlap among GO biological process (BP) concepts among the subnetworks.** Unsurprisingly for gene regulatory subnetworks potentially encoding distinct cell fates, enriched terms overlapped among some of the subnetworks. The bar chart displays an UpsetR analysis of the 20 largest non-redundant sets of enriched GO BP terms.

**SF3d. Functional similarity of the subnetworks’ top20 master regulators’ targets.** In order to identify the most specific enriched terms considering the network structure and hierarchy, we employed the Random Forest machine learning approach classifying subnetwork membership of the master regulators using their targets’ association among the globally enriched GO and PO terms. Globally enriched terms were selected analyzing enrichment in the subnetworks regulatory links using R/NEAT with FDR < 0.01. Multidimensional scaling plot of proximity matrix from R/randomForest of the top20 master regulators for each subnetwork.

**SF3e-i. Top5 most specific concepts to describe targets in subnetworks II, V, VIII, IX and X.** Word cloud representations of the most important terms for each subnetwork to classify subnetwork membership of the top20 master regulators using the Random Forest predictor. We selected and ranked terms for each subnetwork demanding variable importance > 0 using the decrease in node impurity based on the Gini index implemented by the R/randomForest package. See Supplementary Table ST3 for complete results.

**Supplementary Figure SF4. Mosaic plot of inter-subnetwork connections.** Mosaic plot or *Marimekko diagram* showing the cross-sectional distribution of inter-subnetwork connections considering the predicted directionality based on the Pearson correlation coefficient of expression profiles between a regulator and a predicted target from another subnetwork. Color-scale of the boxes depicts the distribution of Pearson residuals indicating significant over-(green) or under-(orange) representation of inter-connecting edges obtained from a significant *Χ^2^* test (p-value < 2.22e^-16^).

**Supplementary Figure SF5. K-shortest simple path network graphs.**

**SF5a. APB regulatory hierarchy.** Network plot depicting the regulatory hierarchy of the AINTEGUMENTA, PLETHORA and BABY BOOM (APB) subfamily of AP2/ANT transcription factors in *P. patens*. The network subgraph was extracted using the PathLinker Cytoscape app (k = 20) and visualized in Cytoscape. Nodes are color-coded by subnetwork identity. Node shapes differentiate between NERD-targeted calpain cleavages (Arg-N-end rule = triangle) and no or other cleavages (round rectangle). Edge arrows indicate predicted directionality based on the Pearson correlation coefficient of expression profiles (R > 0 black arrowhead; R < 0 red square). Edge width is based on the path rank score obtained from the PathLinker algorithm (1. rank = thickest).

**SF5b. DEK1 upstream regulon.** Network plot depicting the upstream regulatory context of DEK1. The network subgraph was extracted using the PathLinker Cytoscape app (k = 50) and visualized in Cytoscape. Nodes are color-coded by subnetwork identity. Node shapes differentiate between NERD-targeted calpain cleavages (Arg-N-end rule = triangle) and no or other cleavages (round rectangle). Edge arrows indicate predicted directionality based on the Pearson correlation coefficient of expression profiles (R > 0 black arrowhead; R < 0 red square). Edge width is based on the path rank score obtained from the PathLinker algorithm (1. rank = thickest).

**Supplementary Figure SF6. Prediction of direct and indirect DEK1 targets.**

**SF6a. Scatter plot of protein length vs. number of predicted calpain cleavage sites.** Overall site frequencies were scaled by the total protein length. Log-transformed, scaled overall site frequencies were clustered with k-means clustering into 5 site abundance level categories (SLC; point colors), representing: S0 = no; S1 = very few – few; S2 = few – medium; S3 = medium – many; S4 = many - very many predicted sites.

**SF6b. PCA of calpain cleavage site predictions.** Overall site frequency and individual NERD site type frequencies (see Fig. 3a and Methods) were scaled by the total protein length and analyzed by PCA. Eigenvector plot of the first two principal components. While overall site frequency and primary acetylated sites contribute only to the first dimension, the NERD-like signatures can also be separated by the second dimension.

**SF6c. Model-based clustering of calpain cleavage site predictions.** Line plot depicting the Bayesian Information Criterion vs. the number of resulting clusters (components) using different clustering models. The VVE model resulted in the optimal BIC.

**SF6d. Optimal model-based clustering (VVE model) of calpain cleavage site predictions.** Cluster scatter plot of the first two principal components of a PCA of calpain cleavage site classifications for *P. patens* proteins.

**SF6e. Fractions of NERD-like cleavages among the VVE clusters color-coded by the overall abundance level of predicted sites.** Box-whisker plots depicting the distribution of the relative proportion of NERD-like cleavages. Proteins with no predicted cleavages were represented as 1. Color-coding according to 5 site abundance level categories (SLC; point colors), representing: S0 = no; S1 = very few – few; S2 = few – medium; S3 = medium – many; S4 = many - very many predicted sites. X-axis corresponds to the three identified clusters obtained by model-based clustering via VVE.

**SF6f. Fractions of NERD-like cleavages among the VVE clusters.** Box-whisker plots depicting the distribution of the relative proportion of NERD-like cleavages. Proteins with no predicted cleavages were represented as 0. Color-coding based on VV3 clustering.

**SF6g. Absolute deregulation of subnetworks in *DEK1* mutants.** Box-whisker plots depicting the distribution of the cumulative level of deregulation of genes with significantly altered expression levels in the mutants encoded by the five DEK1-controlled subnetworks. The levels of deregulation were represented by the sum of the χ2 test statistic of the likelihood ratio tests (LRTs) comparing wild type vs. *Δdek1*, *oex1* vs. wild type and *Δdek1* vs. *oex1* employing it in the sense of an absolute, cumulative effect size. Fill-color depicts subnetwork affiliation.

**SF6h. Regulatory cascades demonstrate consistent deregulation patterns.**

Mosaic plot or *Marimekko diagram* depicting the three-way cross-tabulation of three three factorial variables: left axis – <u>order</u> of the regulon i.e. 1st order: TF→target; 2nd: TF→TF→target and 3rd: TF→TF→TF→target. Upper and left axis represent binary factors representing the binary status of their upstream regulon with respect to predicted levels of DEK1 control. Upper axis – <u>upstream_NERD_cleavage</u> – i.e. predicted direct DEK1 cleavage targets. Right axis – <u>indirect DEK1 targets</u> i.e. deregulation of upstream TFs. Binary status defines whether the regulon comprises TFs predicted direct (x) or indirect (y) DEK1 targets (> 0% of the TFs) or not (= 0% TFs). Boxes are colored based on Pearson residuals from a significant χ2 test of the cross-table comparing the proportions of the three classes.

**Supplementary Figure SF7. Filtered set of predicted direct and indirect DEK1 targets.**

**SF7a. Alluvial plot depicting the distribution of the filtered, predicted direct and indirect DEK1 targets among the 11 subnetworks.** Color-coding of bands reflects directionality of deregulation patterns in the mutant lines (see Fig. 1f for details). The green bands represent unaffected upstream TFs predicted to control the significantly deregulated target genes.

**SF7b. Global network of the filtered, predicted direct and indirect DEK1 targets.** Network plot of the 4,125 direct and indirect DEK1 targets. Nodes are color-coded by subnetwork affiliation. Node size is relative to the nodes’ local reaching centrality (master regulator TFs have largest). Predicted direct DEK1 cleavage targets are depicted as triangles. Indirect DEK1 targets as circles. Edges are color-coded by deregulation pattern i.e. type of DEK1-controlled regulatory interaction: *repressor targets* (red) and *activator targets* (blue).

**SF7c. Distribution of intra- and inter-subnetwork regulatory interactions controlled by DEK1.** Mosaic plot or Marimekko diagram depicting the three-way cross-tabulation of three three factorial variables: <u>Left y-axis</u>: mutant deregulation pattern i.e. type of regulatory interaction *repressor targets* and *activator targets*. <u>Top x-axis</u>: regulator subnetwork affiliation. <u>Right x-axis</u>: target subnetwork affiliation. Box size relative to the number of edges. Boxes are colored based on Pearson residuals from a significant χ2 test of the cross-table (green: enrichment; orange: under-representation).

**Supplementary Figure SF8. Functional characterization of direct and indirect DEK1 targets by ontology analysis.**

**SF8a. Semantic similarity of DEK1-controlled regulatory interactions.** Multidimensional scaling plot of a distance matrix derived by comparing enriched GO biological process terms between the sets of DEK1-controlled regulatory interactions within and between subnetworks using semantic similarity. Sets are depicted as symbols that represent one of six clusters of similar concepts and color-coded by deregulation pattern (DEK1-controlled *repressor* [red] or *activator* [blue] *target*). Point size is relative to the total number of genes in the set.

**SF8b. Model-based clustering of DEK1-controlled regulatory interactions.** Line plot depicting the Bayesian Information Criterion vs. the number of resulting clusters (components) using different clustering models. The EII model with n = 6 resulted in the optimal BIC.

**SF8c. Optimal model-based clustering (EII model) of DEK1-controlled regulatory interactions.** Cluster scatter plot of a multidimensional scaling plot of the semantic similarity/distance matrix (see Panel SF8a of the optimal EII model clustering analysis. Points are color-coded by cluster affiliation. Clusters with more than one gene set are depicted with ellipses depicting the variation around the cluster center (larger symbol of same color).

**SF8d. and SF8e. Automatically inferred representative key concepts of enriched (d) PO anatomical entity and (e) GO biological process for DEK1-controlled regulatory interactions.** Multidimensional scaling plot of a distance matrix derived by comparing enriched ontology terms from the respective ontology partition for the gene sets of DEK1-controlled regulatory interactions. All enriched terms are represented by a point that is scaled by the number of genes with that term in the set. For groups of similar concepts (aggregated by level 3 ontology terms), terms were ranked in ascending order by information content and log FDR value of term enrichment.

**Supplementary Figure SF9. Major phenotypes of the *DEK1* deletion and overexpression mutants**. Schematic overview of *P. patens* gametophyte development in wild type (WT), *DEK1* deletion mutant (*Δdek1*) and DEK1 Linker-Calpain overexpressing line (*oex1*). Numbers indicate days after culture initiation. Primary filament is oriented horizontally with side-branches progressively forming secondary filaments positioned vertically. *Δdek1* mutant shows reduced elongation of secondary filaments and accelerated gametophore bud initiation when compared to WT. Developmentally arrested *Δdek1* buds are often formed in clusters. In contrast to *Δdek1*, *oex1* shows increased elongation of secondary filaments and delayed gametophore bud initiation.

**Supplementary Figure SF10. Deciphering complex phenotypes by Factorial Differential Gene Expression Network Enrichment Analysis (FDGENEA).**

**SF10a. Translation of the complex DEK1 phenotype to phenotypic factors.** Graphical summary of the quantified *DEK1* mutant phenotypes and translation into binary phenotypic factors (first table column in Panel SF10b. Cell numbers are used throughout the other panels to refer to a specific phenotypic trait.

**SF10b. Contrasting phenotypic factors in Differential Gene Expression Network Enrichment Analysis.** Numbers refer to cell numbers containing drawings of the respective phenotype in Panel SF10a. Symbols in the heat map cells represent directionality of differential gene expression for each component trait for each of the subnetworks: + depicts enrichment of up-regulated genes. - depicts enrichment of down-regulated genes. Heatmap fill-color of each cell represents the z-score-scaled, overall enrichment of DEGs (irrespective of their directionality) for each phenotypic factor. Subnetworks X, II and V are predominantly enriched for regulons affecting 2D-to-3D phenotypes.

**SF10c. Enriched subnetworks and upstream regulators associated with high number of buds per filament – subnetwork colors.** Nodes are color-coded by subnetworks. Node sizes relative to the cumulative, absolute fold-change of deregulation of the respective gene and any possible target gene in the mutants. Node shapes: TFs depicted as triangles, TRs as diamonds, miRNAs as inverted triangles and targets as circles. Edge colors: positive correlation of connected nodes in DEK1 RNASeq data (black) and negative correlation coefficient (orange).

**SF10d. Enriched subnetworks and upstream regulators associated with high number of buds per filament – log2foldchanges.** Same graph and attributes as Panel SF10c, but color-coded by directionality of DGE analysis (intensity gradient relative to log-fold change). The three groups of nodes depicted in Panels SF10c and SF10d correspond to genes that are (R) positively associated with a high number of buds (i.e. are up-regulated in *Δdek1* and *Δloop*; right group), (L) display a negative association (i.e. up-regulated in WT, *oex1* and *Δlg3*; left group and down-regulated in *Δdek1* and *Δloop*) or (T) upstream regulators without significant change in gene expression with respect to this phenotype (top group). This clustering is also evident in the type of connections between these groups – especially the two major assemblages harbor many negative regulatory interactions (orange color of links between them). Overall, while subnetwork V dominates the left group, the right group of nodes is more diverse and consists of subnetworks X, XI and IX. Subnetwork II is prominent in both groups. This pattern gives important insight into the implementation of this developmental phase transition – a predominantly negative interaction between the regulatory toolkit driving gametophore development (X) and the gene complement of the primary filament (V; chloronema). This transition between cell states is brokered by the subnetwork II, which is also implementing the other types of secondary filaments (caulonema and rhizoids). Subnetworks IX and XI seem to act in conjunction.

**SF10e. FDGENEA gene sets.** These sets comprise genes that are significantly (LRT q-value < 0.1) associated with any of the phenotypic factors and fall within any of the enriched subnetworks (see Panels SF10a and SF10b).

**SF10f. Top50 intersections between the FDGENEA gene sets.** UpSetR plot depicting the 50 largest non-overlapping FDGENEA gene sets (Panels SF10b and SF10e).

**SF10g. The indirect DEK1 target types display specific enrichment patterns among the FDGENEA sets.** Subset of the FDGENEA gene sets that is predicted to be indirectly targeted by DEK1, i.e. controlled by either an activator or a repressor TF that is directly cleaved by the DEK1 calpain. Enrichment of the respective DEK1 target type in each of the FDGENEA gene sets was tested using the `enrichment_test` method from the RVenn package. P-values for both target types were used to cluster the sets by hierarchical clustering using the `ward.D2` method in R.

**SF10h. All intersections of gene sets associated with traits involving the number of buds per filament i.e. bud initiation.** UpSetR plot depicting non-overlapping FDGENEA gene sets (Panels SF10b and SF10e).

**SF10i. Network of DEK1-controlled, *overbudding* up-regulated genes.** Network plot depicting FDGENEA gene set with significant, positive association to the *number of buds per filament high* trait, predicted to be indirectly or directly controlled by DEK1. Total number of nodes: 901 genes. Node colors based on subnetwork affiliations. Nodes were labeled based on available manual annotations, which are usually indicative of experimental evidence or curation efforts for this study. Edge colors based on mutant expression pattern and directionality of the predicted, DEK1-controlled, regulatory interaction (i.e. blue: *activator targets* and red: *repressor targets*; see Fig 1 and main text for details). Triangular node shape indicates predicted, DEK1-controlled TFs (cleavage targets) and circles their targets. Node sizes relative to overall local reaching network centrality.

**SF10j. Network of DEK1-controlled, *overbudding* up-regulated genes and their top5 DEK1-controlled regulators.** Network plot depicting FDGENEA gene set with significant, positive association to the *number of buds per filament high* trait, predicted to be indirectly controlled by DEK1 and their top5, direct, DEK1-controlled regulators. Total number of nodes: 1,222 genes. The latter were ranked by their overall GENIE3 weight for each target and did not need to be significantly associated with the trait themselves. Node colors based on subnetwork affiliations. Nodes were labeled based on available manual annotations, which are usually indicative of experimental evidence or curation efforts for this study. Edge colors based on mutant expression pattern and directionality of the predicted, DEK1-controlled, regulatory interaction (i.e. blue: *activator targets* and red: *repressor targets*; see Fig 1 and main text for details). Triangular node shape indicates predicted, DEK1-controlled TFs (cleavage targets) and circles their targets. Node sizes relative to overall local reaching network centrality.

**SF10k. Full regulatory context of the CLV1b gene is enriched for genes involved in flowering plant meristem initiation and maintenance that are positively associated with bud initiation.** Network plot depicting the regulatory context of CLV1b, i.e. genes that are predicted to be controlled by the same regulators as CLV1b and TFs. Selected were only genes that are not more than two orders away, i.e. maximally direct targets of regulators of regulators of CLV1b. Triangular node shapes indicate predicted direct NERD-like calpain cleavage. Node colour scaled according to log2foldchange from the FDGENEA analysis of the overbudding trait (up = positive association = red; down = negative association = blue; grey = LRT q-value >= 0.1).

**SF10l. Partial overlap between DEK1-controlled AP2 and MYB TFs in subnetworks II.** Venn diagram depicting intersections between the target lists of the ten subnetwork II AP2 (APB-3, STEMIN3 as well as orthologs of PUCHI, DRN, DRNL, STEMIN3) and five subnetwork X MYB (GAMYB and orthologs of MYB80 and MIXTA) TFs. Targets were selected from *overbudding* up-regulated, predicted DEK1-controlled regulatory interactions (Fig. SF10j) choosing 1st and 2nd degree outgoing connections of said TFs (Table ST11; columns B & E).

**SF10m. Partial overlap between DEK1-controlled AP2 and MYB TFs in subnetworks II.** Venn diagram depicting intersections between the target lists of the ten subnetwork II AP2 (APB-3, STEMIN3 as well as orthologs of PUCHI, DRN, DRNL, STEMIN3) and five subnetwork X MYB (GAMYB and orthologs of MYB80 and MIXTA) TFs. Targets were selected from the extended *overbudding* up-regulated, predicted DEK1-controlled regulatory interactions that included also the up to top5 DEK-controlled regulators independent of their deregulation pattern with respect to *overbudding* (Fig. SF10j) choosing 1st and 2nd degree outgoing connections of said TFs (Table ST11; columns C&F).

**Supplementary Figure SF11. Molecular analysis of the linker-calpain overexpressing (*oex1*) strain.**

**SF11a. The *108* locus (T-5’, T-3’) in the wild type (WT).** Schematic representation of 5’ and 3’ targeting sites (green filled boxes) for the insertion of the PpDEK1 Linker-Calpain overexpression construct at the neutral *108* locus (Schaefer and Zrÿd, 1997) in the *P. patens* WT genotype. Used restriction sites are marked and annotated as ticks.

**SF11b. Schematic representation of the *108* locus in the *oex1* genotype.** Designed targeting of the *Pp*DEK1 <u>Linker-Calpain</u> construct (blue, thick arrows) under a maize <u>ubi</u>quitin <u>promoter</u> (light blue thick arrow) and <u>Ter</u>minator (light blue thick box) coupled with a hygromycin-resistance cassette (<u>35S</u> promoter – *<u>HptII</u>* hygromycin resistance gene; red box). Positions of the restriction sites (black ticks), <u>DIG-probes</u> (green and red framed boxes) and primers (black arrows) used in downstream analyses (Figs. SF11c,d,e) are marked and annotated accordingly.

**SF11c. Southern blot analysis *of oex1* transformant.** The Southern blot analysis performed with two different restriction enzymes, <u>Bgl II</u> and <u>Xcm I</u>, indicates that *oex1* is a stable transformant. The line *oex1* contains multiple concatenated copies of the *Pp*DEK1 Linker-Calpain construct inserted at a single genomic position, the locus *108*. This insertion pattern represents the majority of the mutants obtained with this transformation procedure in *P. patens* (Kamisugi et al. 2006) and is not known to induce any phenotype per se. Lanes 1, 3, 5 and 7 contain wild type genomic DNA. Lanes 2, 4, 6 and 8 contain *oex1* genomic DNA. Lanes 1, 2, 5 and 6: <u>Bgl II</u> restriction digestions; lanes 3, 4, 7, 8,10: <u>Xcm I</u> restriction digestion. Lane L: Lambda phage-HindIII DIG labeled marker.

**SF11d. Detection of the *Pp*DEK1 Linker-Calpain protein in *oex1*.** Western blot detecting *Pp*DEK1 CysPc epitope with polyclonal anti-*Pp*DEK1 CysPc-C2L antibody. L: protein ladder, 1, 2: WT, 3, 4: *Δdek1*, 5, 6: *oex1*. Protein samples in lanes 1, 3, 5 were heated before PAGE, samples 2, 4, 6 were not heated before PAGE. Chemiluminescence was detected followed by imaging of the prestained dual color protein marker (Bio-Rad) on the same membrane. The marker image (lane L; blue frame) was merged with the chemiluminescence scan. The *oex1* line displays a clear signal between the 90 and 150 kDa marker size (lanes 5, 6), that is absent in WT (lanes 1,2) or *Δdek1* (lane 3, 4) and corresponds to the predicted 120 kDa Linker-Calpain protein. The bottom part of the membrane scan contains a uniform identical background banding signal present in all tested genotypes generated by the polyclonal anti-*Pp*DEK1 CysPc-C2L antibody. Original TIFF whole versions of the blot and the gel are provided in the supplementary data archive (`wetlab/gels_and_blots/Western/C2*.tif`).

**SF11e. PCR genotyping confirms genomic insertion in the *108* locus’ 5’ region.** Primer pairs are indicated below the gel image. L: GeneRuler 1 kb DNA ladder, lanes 1, 3, 5, 7 WT DNA template, 2, 4, 6, 8 *oex1* DNA template. The specific amplification signal from the 5’ flank of the construct in *oex1* (lane 4) but not in the WT (lane 3) and not from the 3’ flank (lanes 6 and 8) points to the insertion of the *Pp*DEK1 Linker-Calpain construct at the 5’ site of the 108 locus. The presence of a full *108* signal in *oex1 (*lane 2) indicates the presence of an intact *108* fragment probably associated with the aforementioned complex concatenated insertion.

**SF11f. Genomic mapping of RNASeq data confirms *DEK1* mutant genotypes and expected transcriptional outcomes.** Integrative Genome Viewer (IGV) tracks displaying the coverage and alignments from a mapping of RNASeq data of the four *DEK1* mutants and the wild type (WT) at day 14 of gametophytic development with respect to the V3.3 gene structure of *DEK1* (last track). Each of the RNASeq tracks is composed of two subtracks: 1) an upper subtrack displaying the coverage as a density plot. Peak minimum and maximum number of reads depicted in the upper left corner of the subtrack (e.g. *oex1* [0-18015]). 2) lower subtrack displays exemplary individual read alignments (color-coded by SAM alignment flags, non-grey colors usually indicating aberrant/low quality alignments). The last track shows the predicted gene structure of *DEK1* in genome annotation V3.3 color-coded by the domain architecture of the DEK1 protein depicted in SF11g. Boxes depict exonic and connecting thin lines the intronic regions of the gene. Coverage histograms and read alignments are consistent with the expected genotype of the respective *DEK1* mutant, with reads lacking entirely or partially in the full (*Δdek1*) or partial (*Δlg3* and *Δloop*) deletion lines as well as an over-accumulation of reads only in the regions encoding the Linker-Calpain domains in the *oex1* strain.

**SF11g. DEK1 protein domain structure.** Overview of the DEK1 protein domain architecture. Individual domains and transmembrane regions are color-coded and annotated. Colored boxes represent conserved and/or functional domains and transmembrane regions, while the grey backbone depicts less conserved (spacer) regions.

#### Supplementary Tables

Individual sheets in the file `SupplementaryTables.xlsx`.

**Supplementary Table ST1.** Summary table of the differential gene expression analysis of *DEK1* mutants.

**Supplementary Table ST2.** Summary table of overall deregulated biological processes, molecular functions, cellular components, anatomical entities and developmental stages in *DEK1* mutants as represented by the respective Gene Ontology (GO) and Plant Ontology (PO) partitions.

**Supplementary Table ST3.** Enriched ontology terms for the predicted *P. patens* gene regulatory subnetworks.

**Supplementary Table ST4.** Genes encoded by the five DEK1-controlled subnetworks.

**Supplementary Table ST5.** Summary table of the predicted regulatory interactions within and between the 11 subnetworks.

**Supplementary Table ST6.** Final filtered set of DEK1-controlled regulatory interactions.

**Supplementary Table ST7.** Enriched ontology terms for the filtered set of DEK1-controlled regulatory interactions (all targets and per subnetwork-pair/deregulation pattern type).

**Supplementary Table ST8.** Factorial Differential Gene Expression Network Enrichment Analysis (FDGENEA) of 17 phenotypic factors.

**Supplementary Table ST9.** Summary table of the DEK1-controlled regulatory interactions for *overbudding* associated genes.

**Supplementary Table ST10.** Full annotation of the genes associated with the *overbudding* phenotype (FDGENEA) including the cell type specific transcriptome data (Frank and Scanlon 2015; Figs 2a,b).

**Supplementary Table ST11.** Gene sets of the AP2 and MYB controlled circuits depicted in Figs. SF10l,m.

**Supplementary Table ST12.** Primers used in this study.

