## Supplementary Figures for "Calpain DEK1 acts as a developmental switch gatekeeping cell fate transitions"

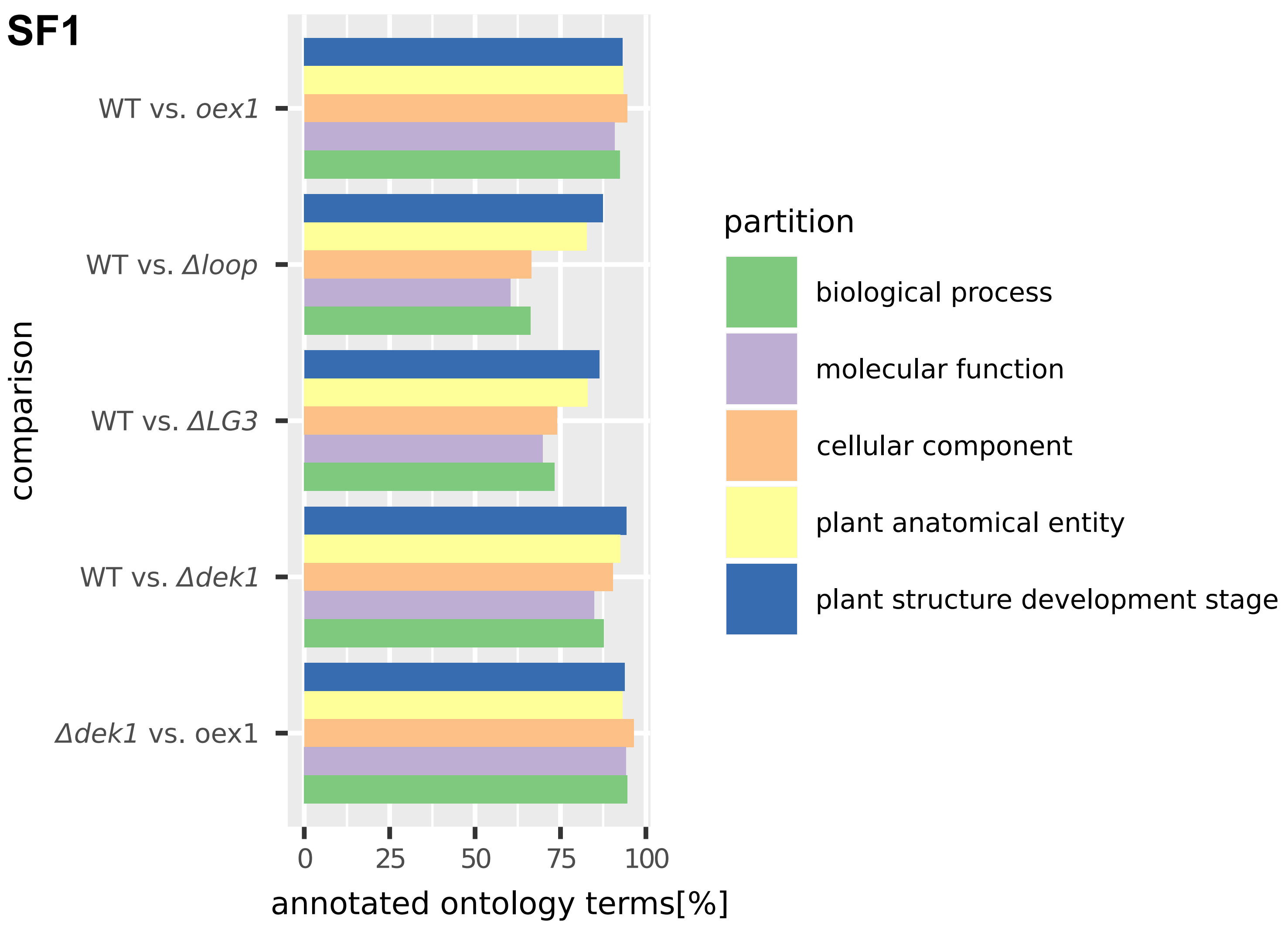
SF1
WT vs. oex1
WT vs. Δloop
WT vs. ΔLG3
WT vs. Δdek1
Δdek1 vs. oex1
comparison
0
25
50
75
100
annotated ontology terms[%]
partition
biological process
molecular function
cellular component
plant anatomical entity
plant structure development stage

**SF2**

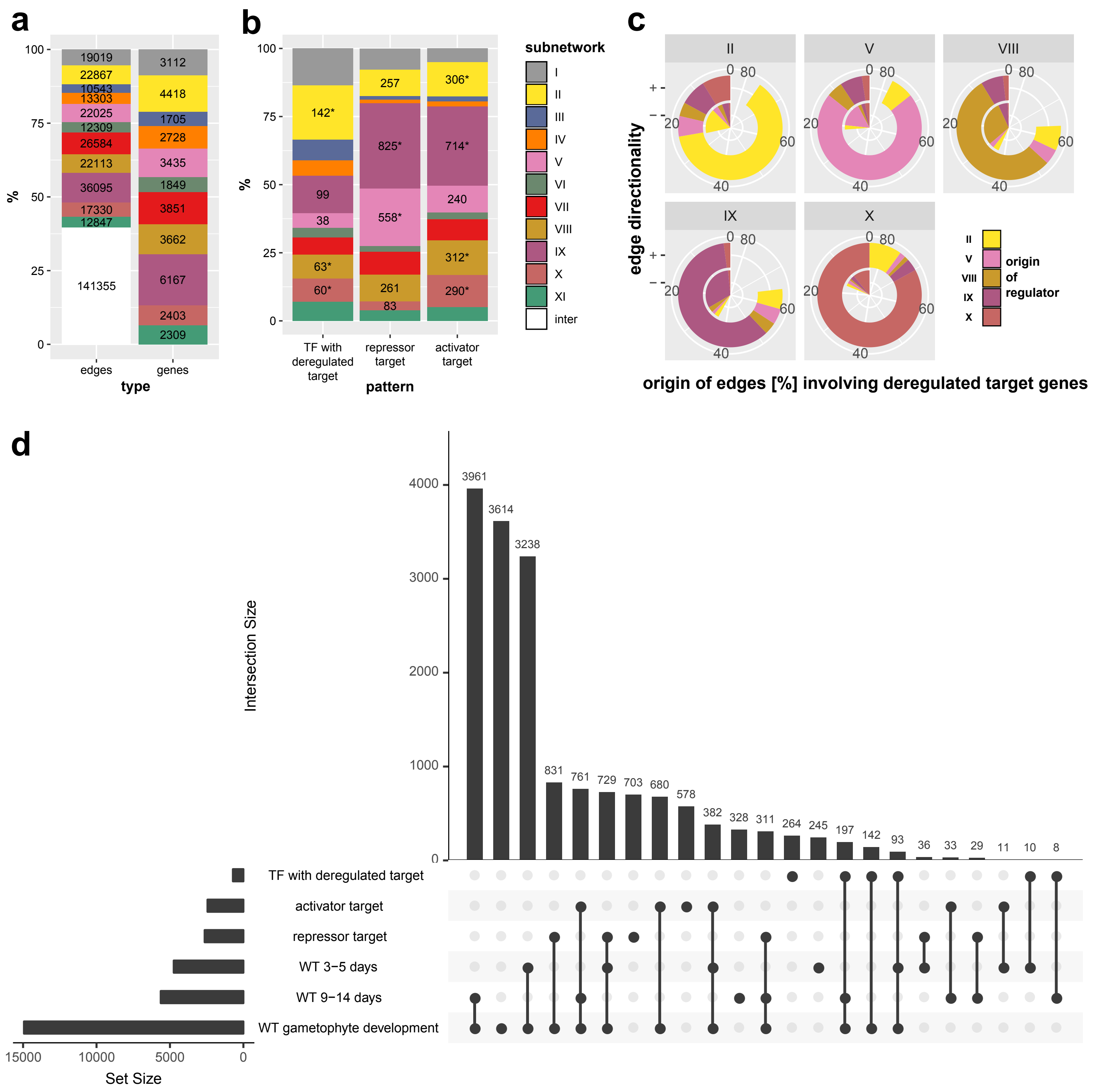
a
100
75
50
25
0
%
19019
22867
10543
13303
22025
12309
26584
22113
36095
17330
12847
141355
3112
4418
1705
2728
3435
1849
3851
3662
6167
2403
2309
edges
genes
type
b
%
142*
99
38
63*
60*
257
825*
558*
261
83
306*
714*
240
312*
290*
TF with deregulated target
repressor target
activator target
pattern
subnetwork
I
II
III
IV
V
VI
VII
VIII
IX
X
XI
inter
c
II
V
VIII
IX
X
0
20
40
60
80
edge directionality
+ -
- -
II
V
VIII
IX
X
origin of regulator
origin of edges [%] involving deregulated target genes
d
Intersection Size
4000
3000
2000
1000
0
3961
3614
3238
831
761
729
703
680
578
382
328
311
264
245
197
142
93
36
33
29
11
10
8
TF with deregulated target
activator target
repressor target
WT 3−5 days
WT 9−14 days
WT gametophyte development
15000
10000
5000
0
Set Size

SF3a

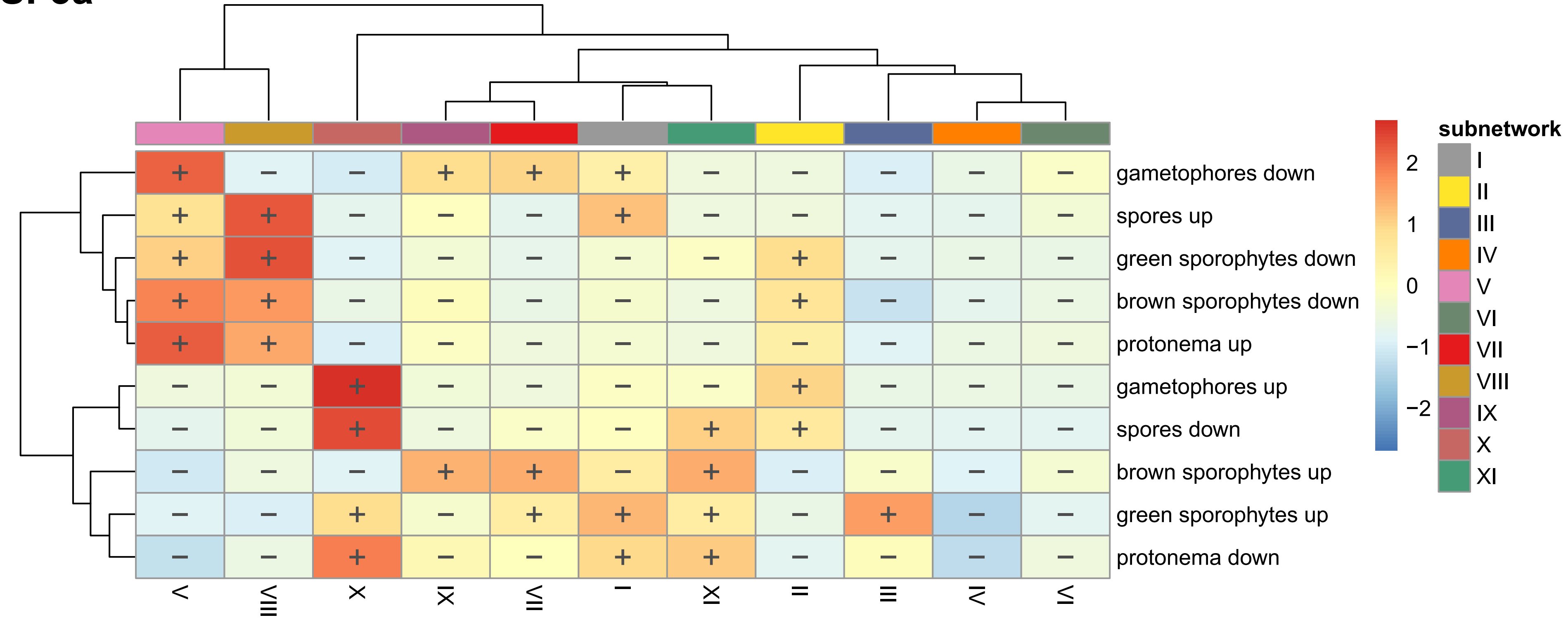
gametophores down
spores up
green sporophytes down
brown sporophytes down
protonema up
gametophores up
spores down
brown sporophytes up
green sporophytes up
protonema down
V
VIII
X
IX
VII
I
XI
II
III
IV
VI
2
1
0
−1
−2
subnetwork
I
II
III
IV
V
VI
VII
VIII
IX
X
XI

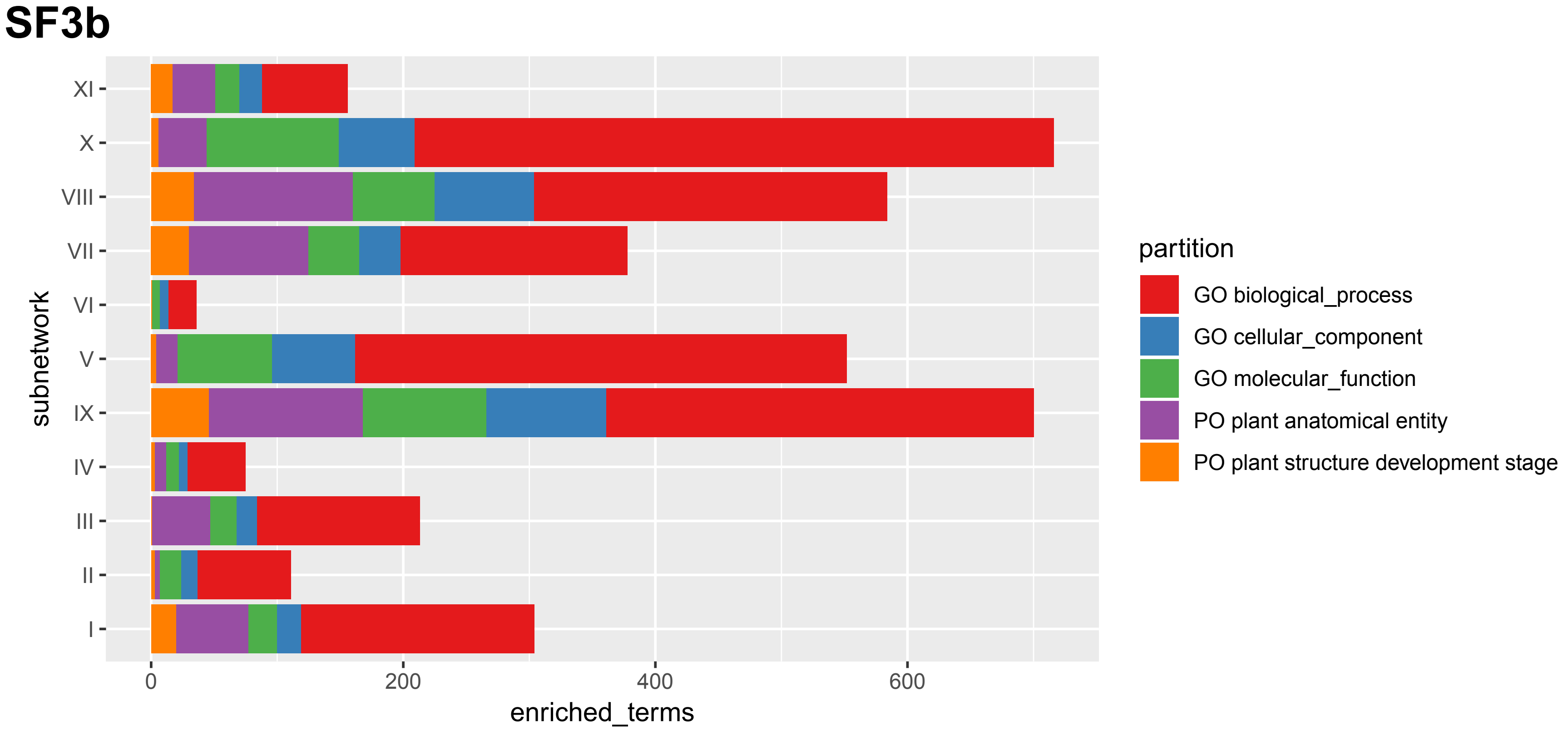
SF3b
subnetwork
XI
X
VIII
VII
VI
V
IX
IV
III
II
I
0
200
400
600
enriched_terms
partition
GO biological_process
GO cellular_component
GO molecular_function
PO plant anatomical entity
PO plant structure development stage

SF3c

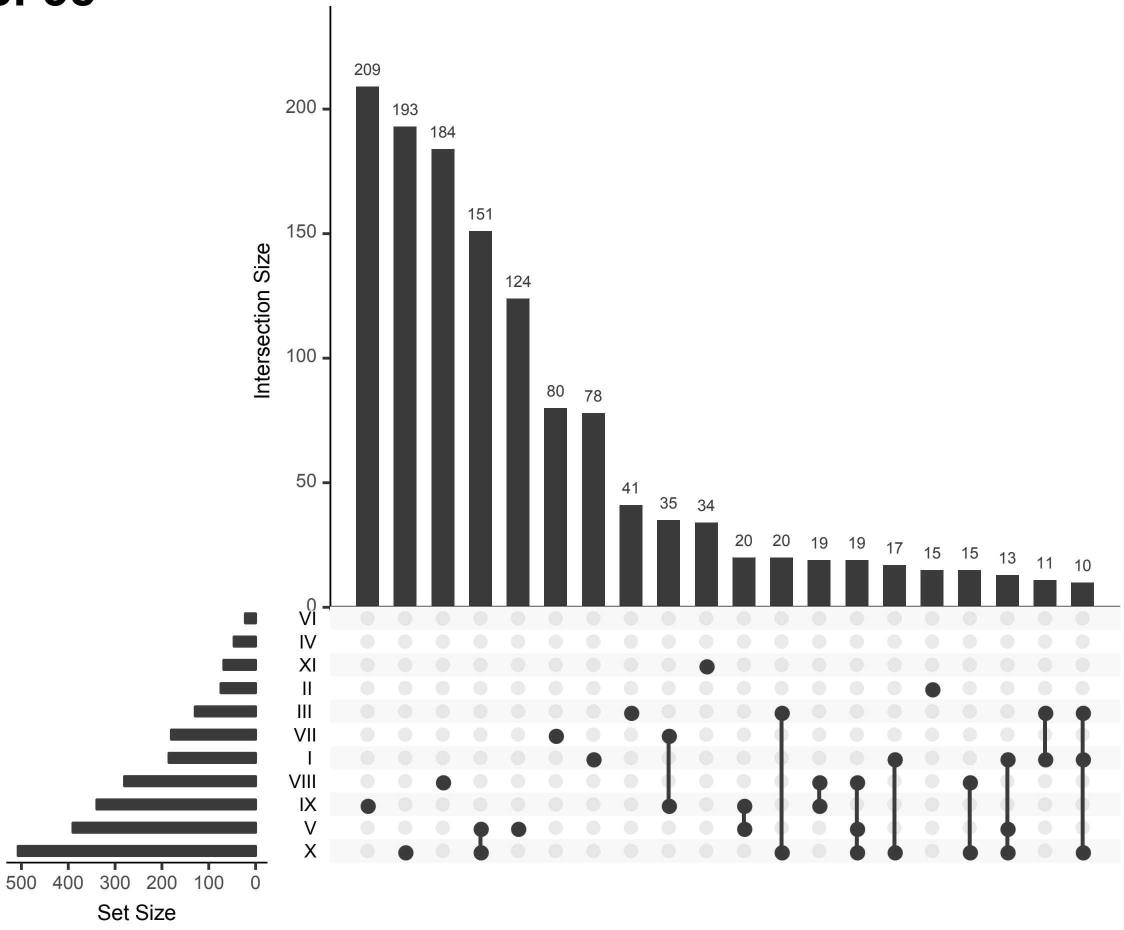

209
193
184
151
124
80
78
41
35
34
20
20
19
19
17
15
15
13
11
10
Intersection Size
200
150
100
50
0
VI
IV
XI
II
III
VII
I
VIII
IX
V
X
500
400
300
200
100
0
Set Size

SF3d

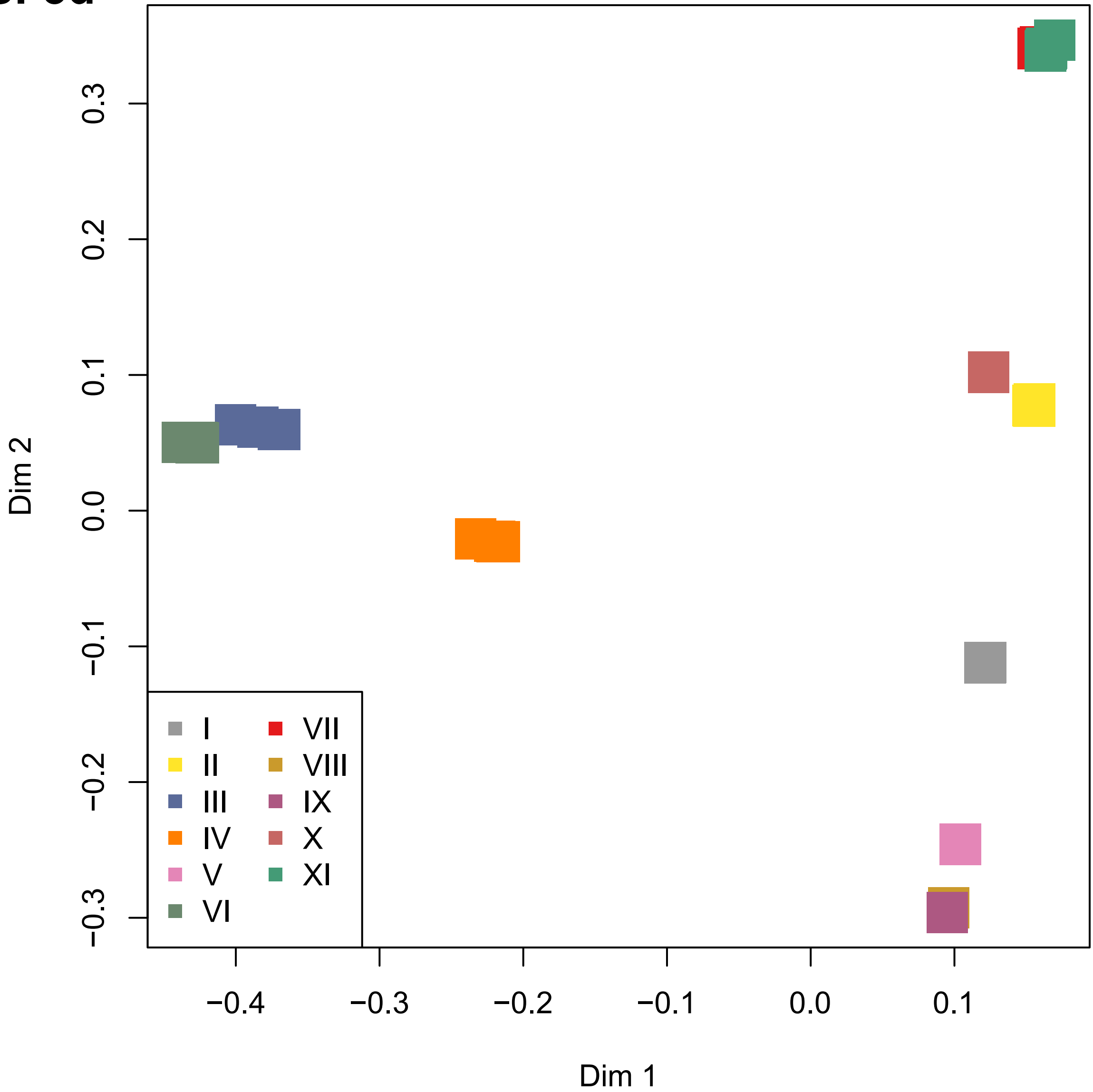

0.3
0.2
0.1
0.0
−0.1
−0.2
−0.3
Dim 2
−0.4
−0.3
−0.2
−0.1
0.0
0.1
Dim 1
I
II
III
IV
V
VI
VII
VIII
IX
X
XI

**SF3e**


gametophyte vegetative stage
response to carbohydrate
cell periphery
organelle lumen
nucleus
cytosol
response to lipid
response to endogenous stimulus
protonema sub−apical initial cell
photoperiodism
external encapsulating structure
long−day photoperiodism
protonema whole plant development stage

**SF3f**

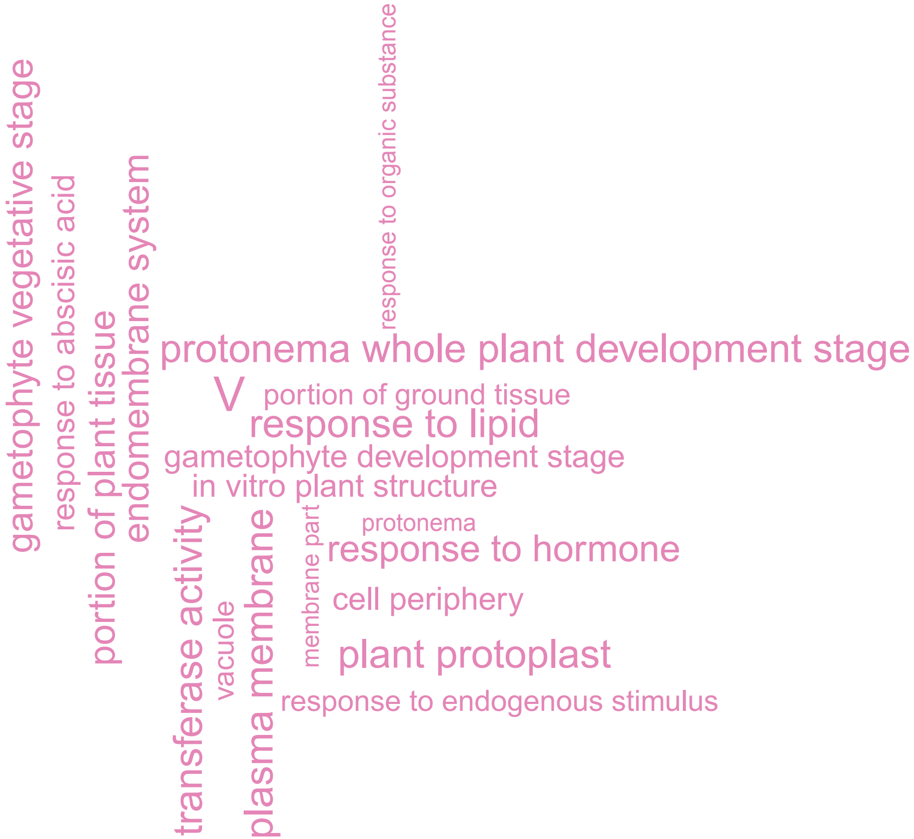
response to organic substance
gametophyte vegetative stage
response to abscisic acid
portion of plant tissue
endomembrane system
protonema whole plant development stage
V
portion of ground tissue
response to lipid
gametophyte development stage
in vitro plant structure
transferase activity
vacuole
plasma membrane
membrane part
protonema
response to hormone
cell periphery
plant protoplast
response to endogenous stimulus

**SF3g**

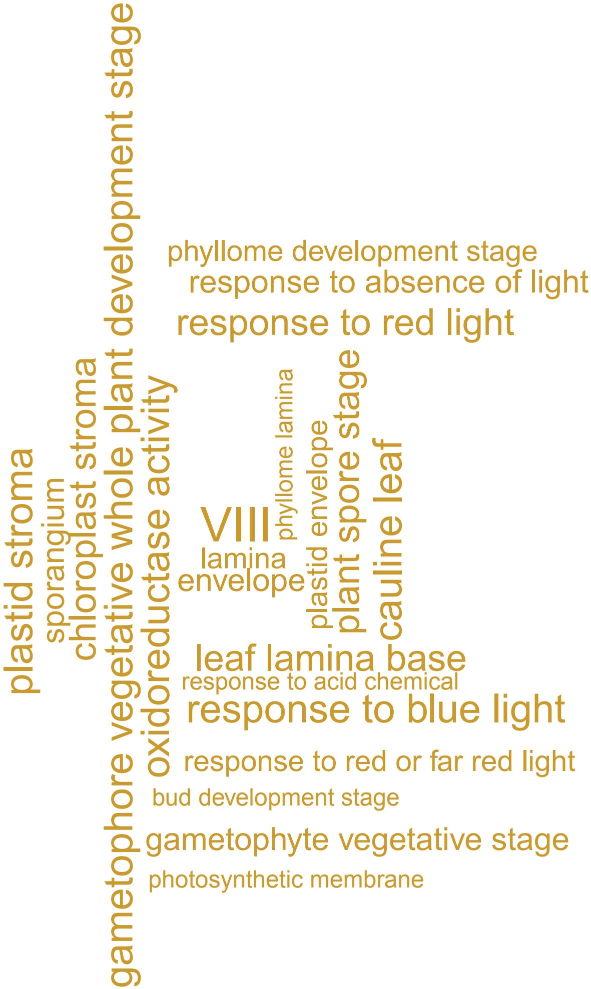
gametophore vegetative whole plant development stage
plastid stroma
sporangium
chloroplast stroma
oxidoreductase activity
phyllome development stage
response to absence of light
response to red light
phyllome lamina
plastid envelope
plant spore stage
cauline leaf
VIII
lamina
envelope
leaf lamina base
response to acid chemical
response to blue light
response to red or far red light
bud development stage
gametophyte vegetative stage
photosynthetic membrane

**SF3h**

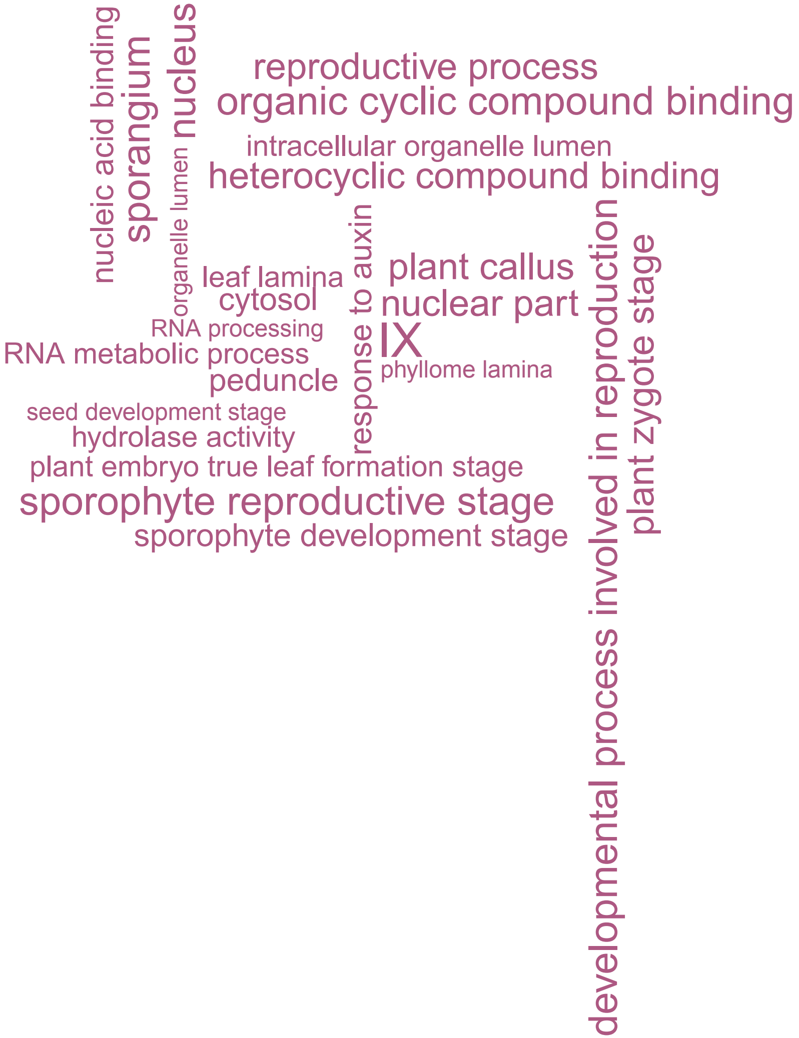
nucleic acid binding
sporangium
nucleus
organelle lumen
reproductive process
organic cyclic compound binding
intracellular organelle lumen
heterocyclic compound binding
leaf lamina
cytosol
response to auxin
plant callus
nuclear part
RNA processing
IX
RNA metabolic process
phyllome lamina
peduncle
seed development stage
hydrolase activity
plant embryo true leaf formation stage
sporophyte reproductive stage
sporophyte development stage
developmental process involved in reproduction
plant zygote stage

**SF3i**

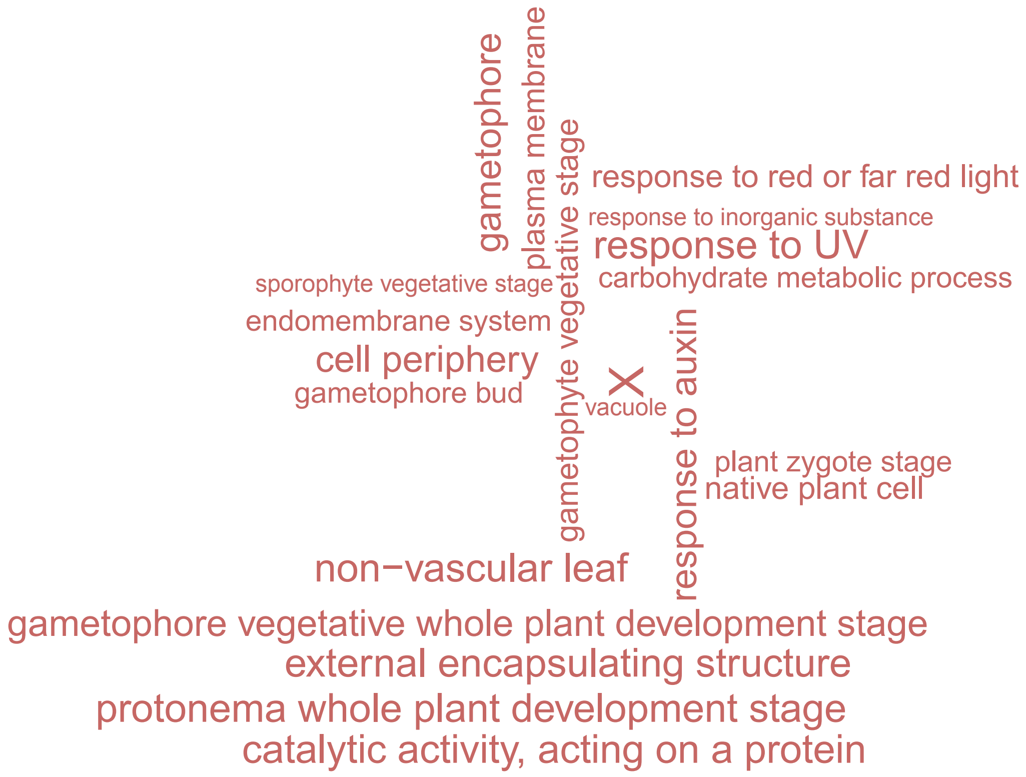
gametophore
plasma membrane
gametophyte vegetative stage
response to red or far red light
response to inorganic substance
response to UV
carbohydrate metabolic process
sporophyte vegetative stage
endomembrane system
cell periphery
gametophore bud
X
vacuole
response to auxin
plant zygote stage
native plant cell
non−vascular leaf
gametophore vegetative whole plant development stage
external encapsulating structure
protonema whole plant development stage
catalytic activity, acting on a protein

**SF4**

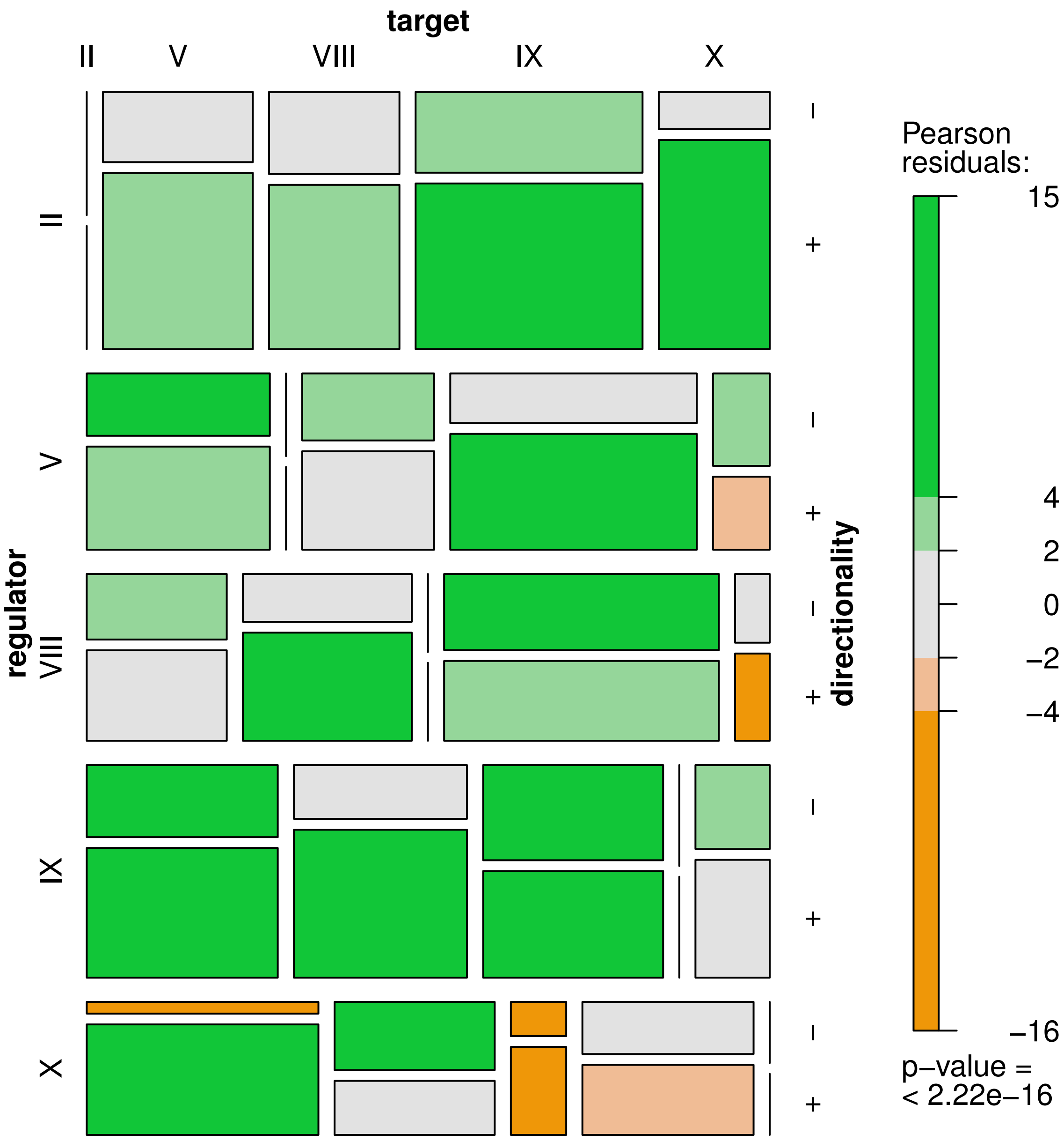
target
II
V
VIII
IX
X
regulator
II
V
VIII
IX
X
directionality
-
+
Pearson residuals:
15
4
2
0
−2
−4
−16
p−value =
< 2.22e−16

**SF5a**

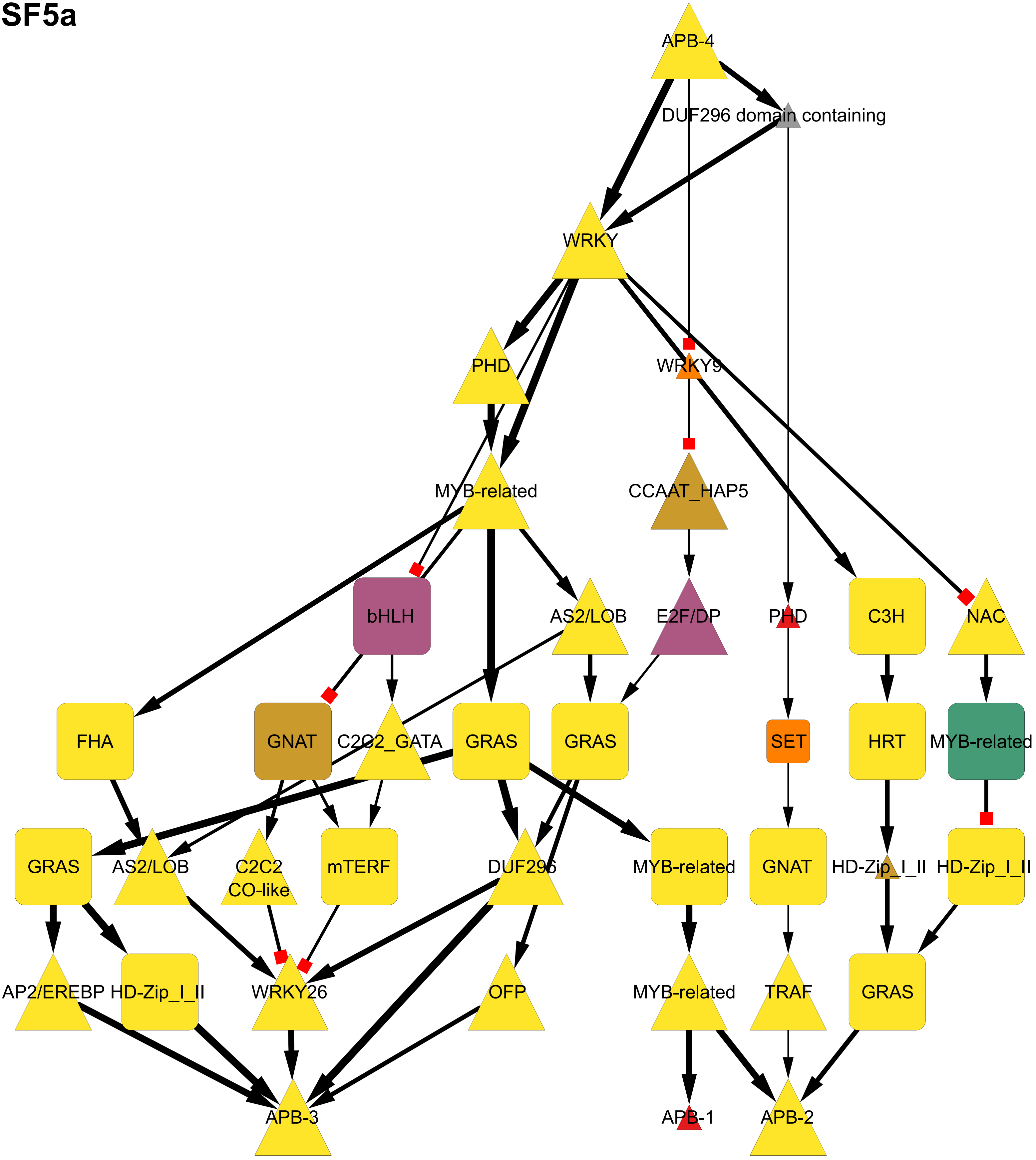
APB-4
DUF296 domain containing
WRKY
PHD
WRKY9
MYB-related
CCAAT_HAP5
bHLH
AS2/LOB
E2F/DP
PHD
C3H
NAC
FHA
GNAT
C2C2_GATA
GRAS
GRAS
SET
HRT
MYB-related
GRAS
AS2/LOB
C2C2
CO-like
mTERF
DUF296
MYB-related
GNAT
HD-Zip_I_II
HD-Zip_I_II
AP2/EREBP
HD-Zip_I_II
WRKY26
OFP
MYB-related
TRAF
GRAS
APB-3
APB-1
APB-2

**SF5a**

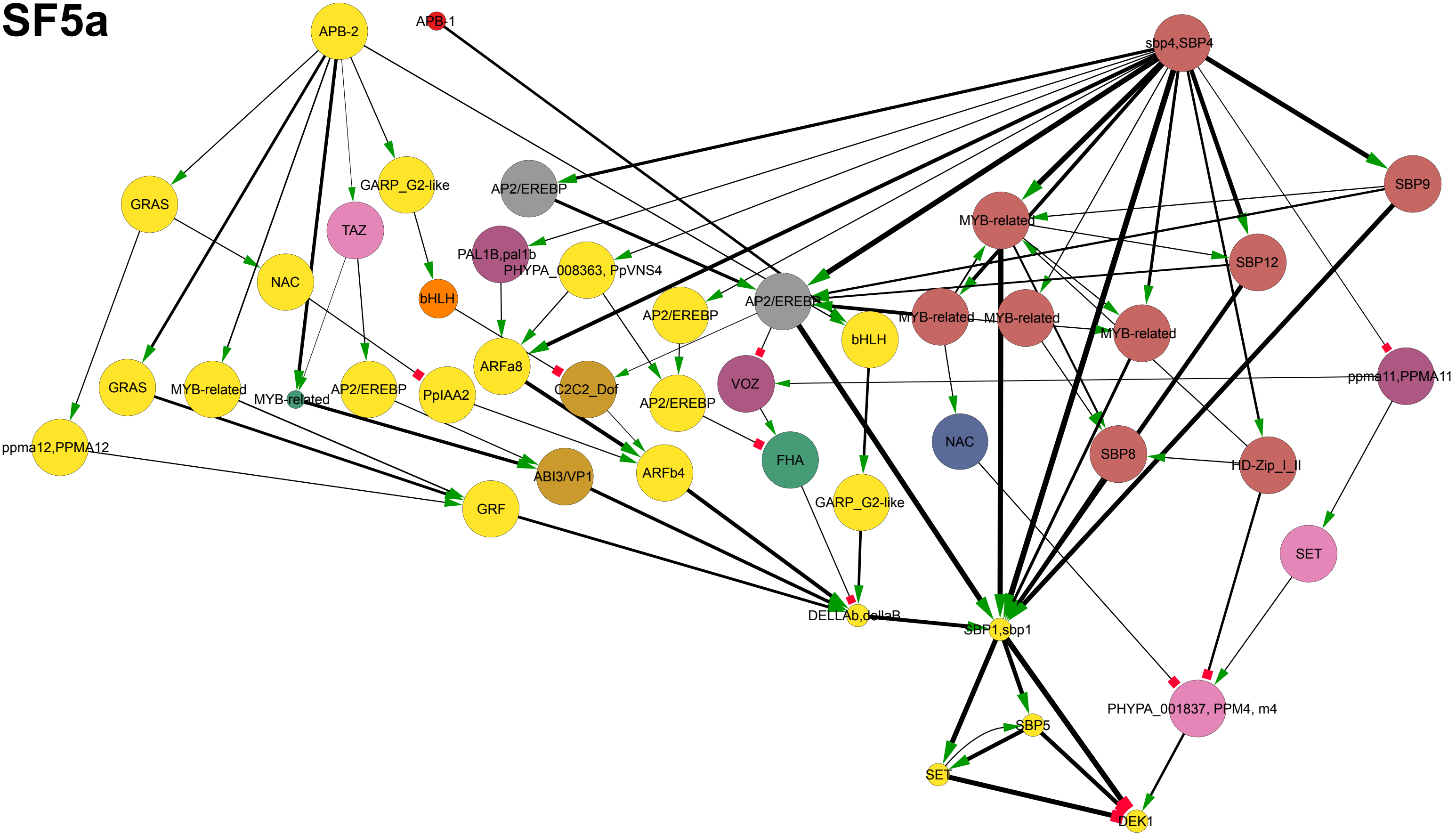
APB-2
APB-1
sbp4,SBP4
GRAS
GARP_G2-like
AP2/EREBP
TAZ
SBP9
MYB-related
PAL1B,pal1b
PHYPA_008363, PpVNS4
SBP12
NAC
bHLH
AP2/EREBP
AP2/EREBP
MYB-related
MYB-related
MYB-related
bHLH
ARFa8
GRAS
MYB-related
MYB-related
AP2/EREBP
PpIAA2
C2C2_Dof
AP2/EREBP
VOZ
ppma11,PPMA11
NAC
ppma12,PPMA12
ABI3/VP1
ARFb4
FHA
SBP8
HD-Zip_I_II
GRF
GARP_G2-like
SET
DELLAb,dellaB
SBP1,sbp1
PHYPA_001837, PPM4, m4
SBP5
SET
DEK1

**SF6a**

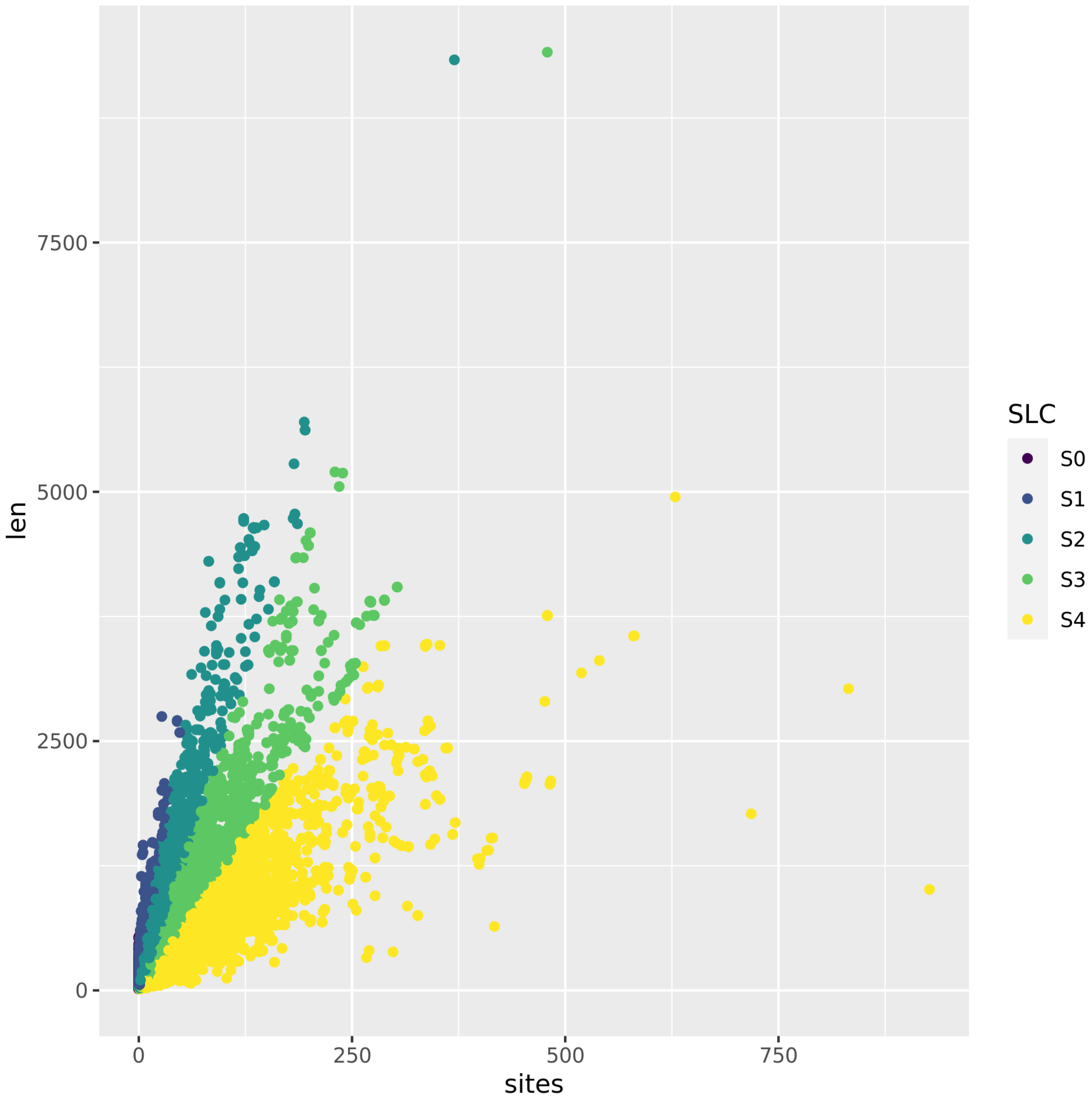

**SF6b**

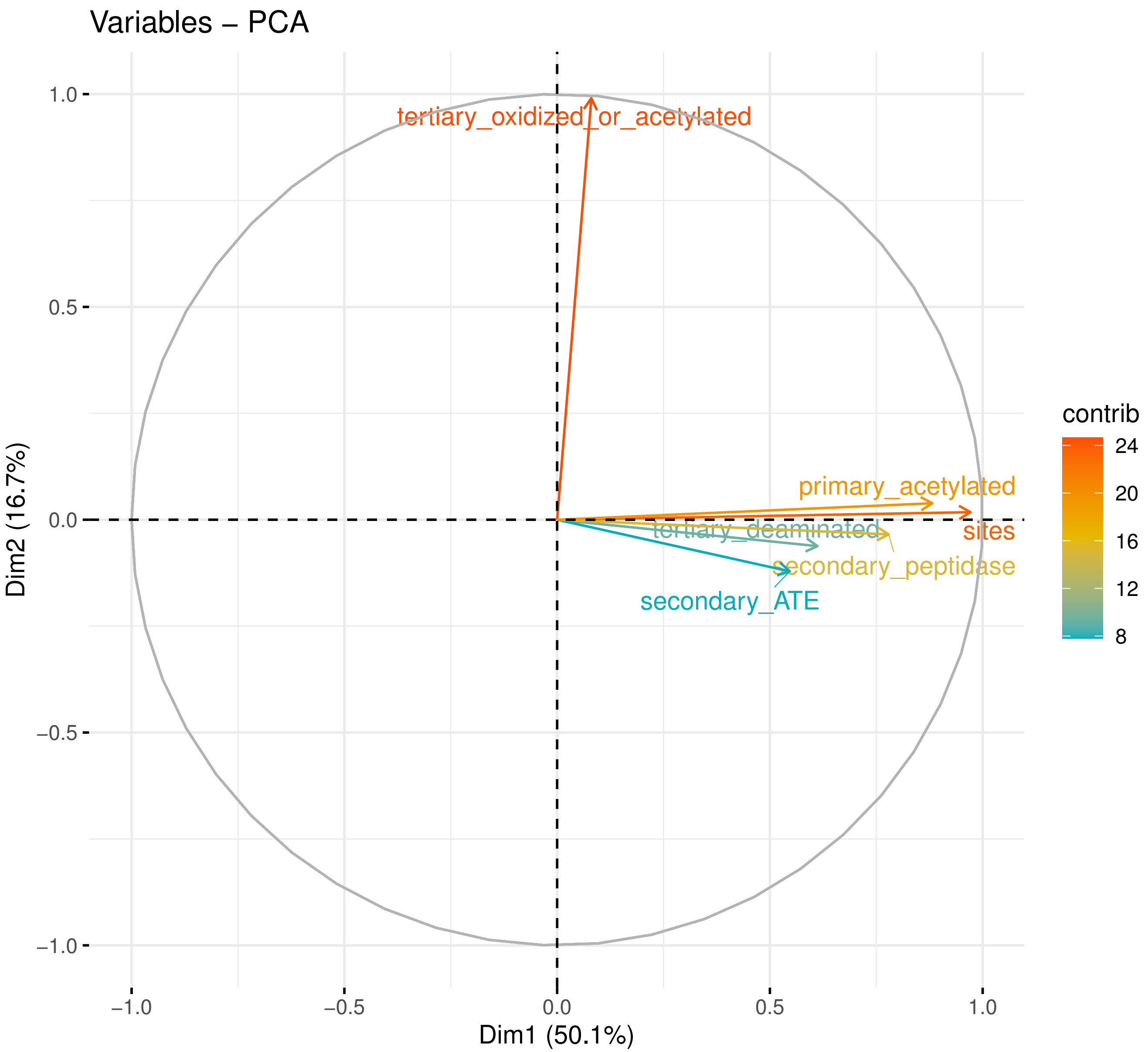
Variables − PCA
tertiary_oxidized_or_acetylated
primary_acetylated
tertiary_deaminated
sites
secondary_peptidase
secondary_ATE
contrib
24
20
16
12
8
1.0
0.5
0.0
−0.5
−1.0
Dim2 (16.7%)
−1.0
−0.5
0.0
0.5
1.0
Dim1 (50.1%)

**SF6c**

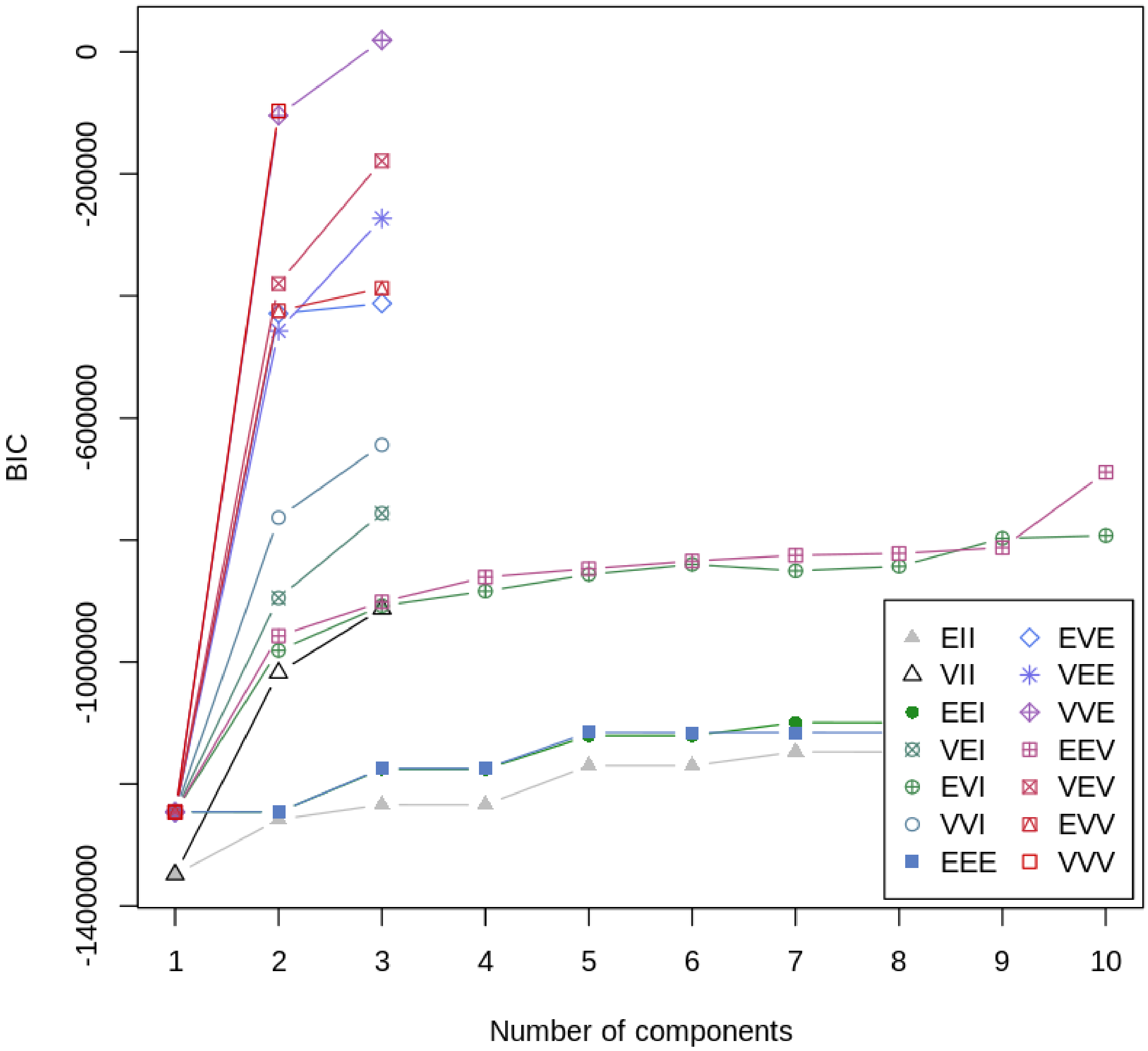

BIC
0
-200000
-600000
-1000000
-1400000
1
2
3
4
5
6
7
8
9
10
Number of components
EII
VII
EEI
VEI
EVI
VVI
EEE
EVE
VEE
VVE
EEV
VEV
EVV
VVV

## SF6d

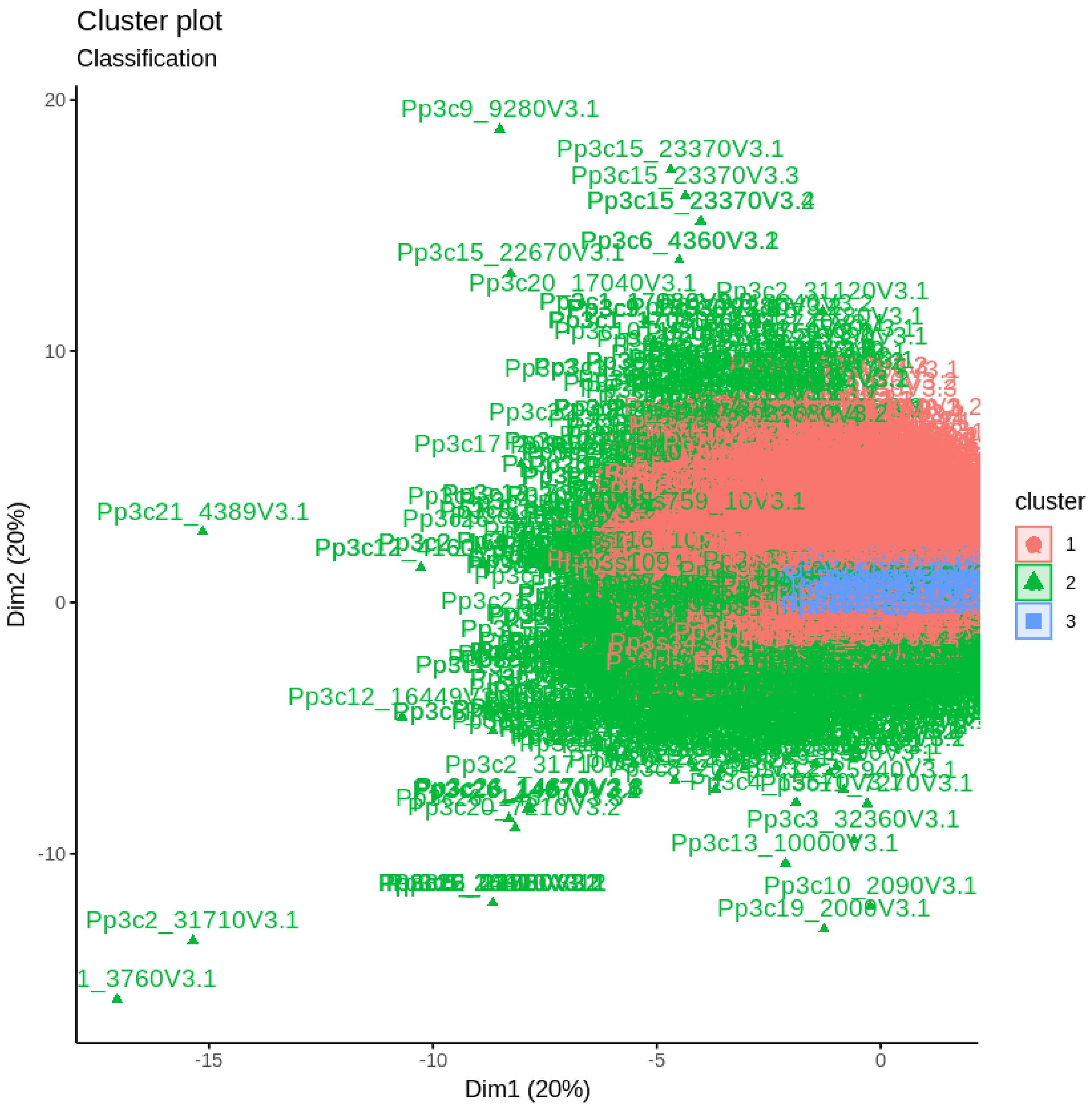
Cluster plot
Classification
Pp3c9_9280V3.1
Pp3c15_23370V3.1
Pp3c15_23370V3.3
Pp3c15_22670V3.1
Pp3c20_17040V3.1
Pp3c21_4389V3.1
Pp3c12_16449V
Pp3c3_32360V3.1
Pp3c13_10000V3.1
Pp3c10_2090V3.1
Pp3c19_2000V3.1
Pp3c2_31710V3.1
1_3760V3.1
Dim2 (20%)
Dim1 (20%)
20
10
0
-10
-15
-10
-5
0
cluster
1
2
3

# SF6e

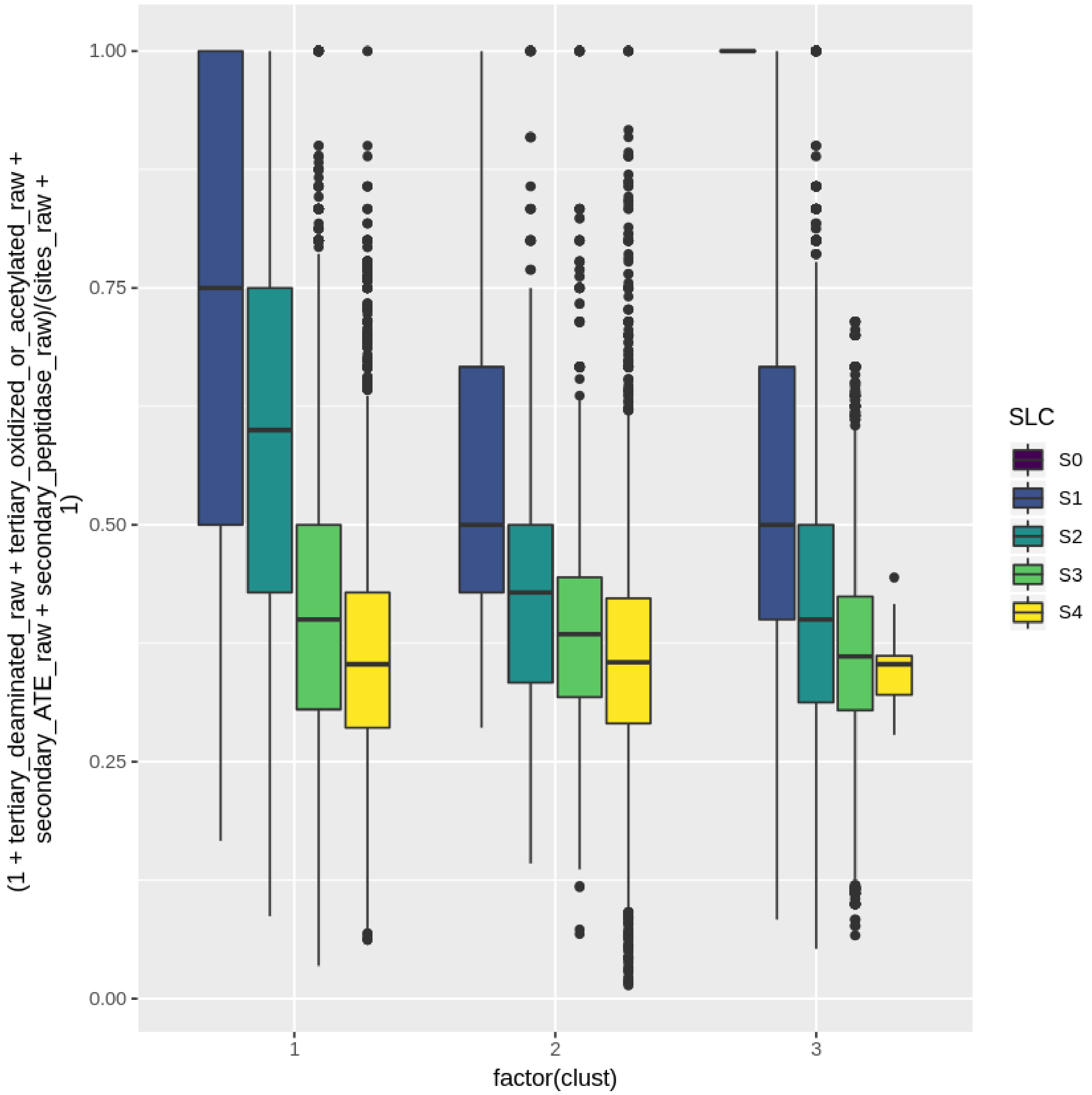

(1 + tertiary_deaminated_raw + tertiary_oxidized_or_acetylated_raw + secondary_ATE_raw + secondary_peptidase_raw)/(sites_raw + 1)
1.00
0.75
0.50
0.25
0.00
1
2
3
factor(clust)
SLC
S0
S1
S2
S3
S4

**SF6f**

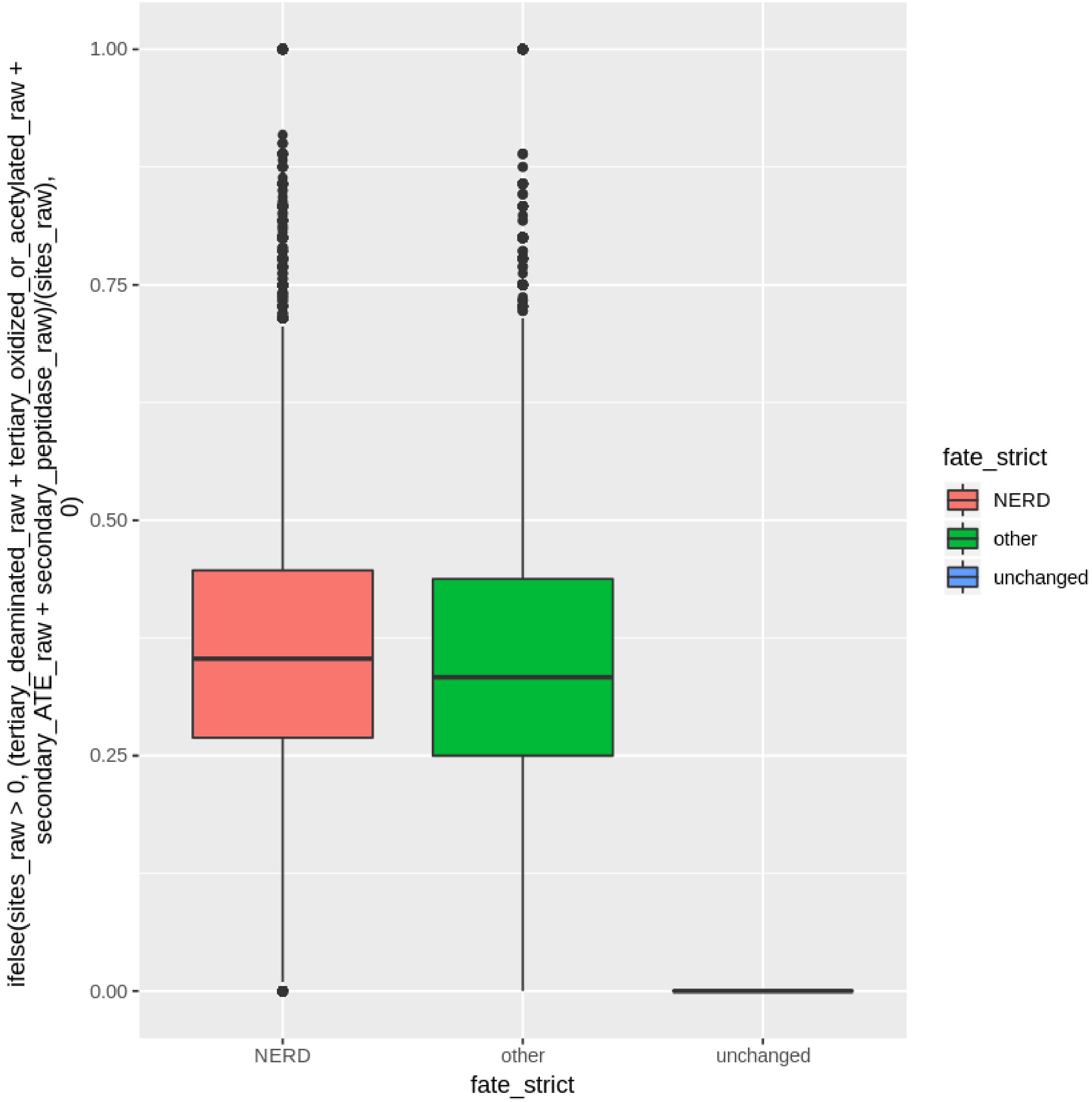
ifelse(sites_raw > 0, (tertiary_deaminated_raw + tertiary_oxidized_or_acetylated_raw + secondary_ATE_raw + secondary_peptidase_raw)/(sites_raw), 0)
1.00
0.75
0.50
0.25
0.00
NERD
other
unchanged
fate_strict
fate_strict
NERD
other
unchanged

# SF6g

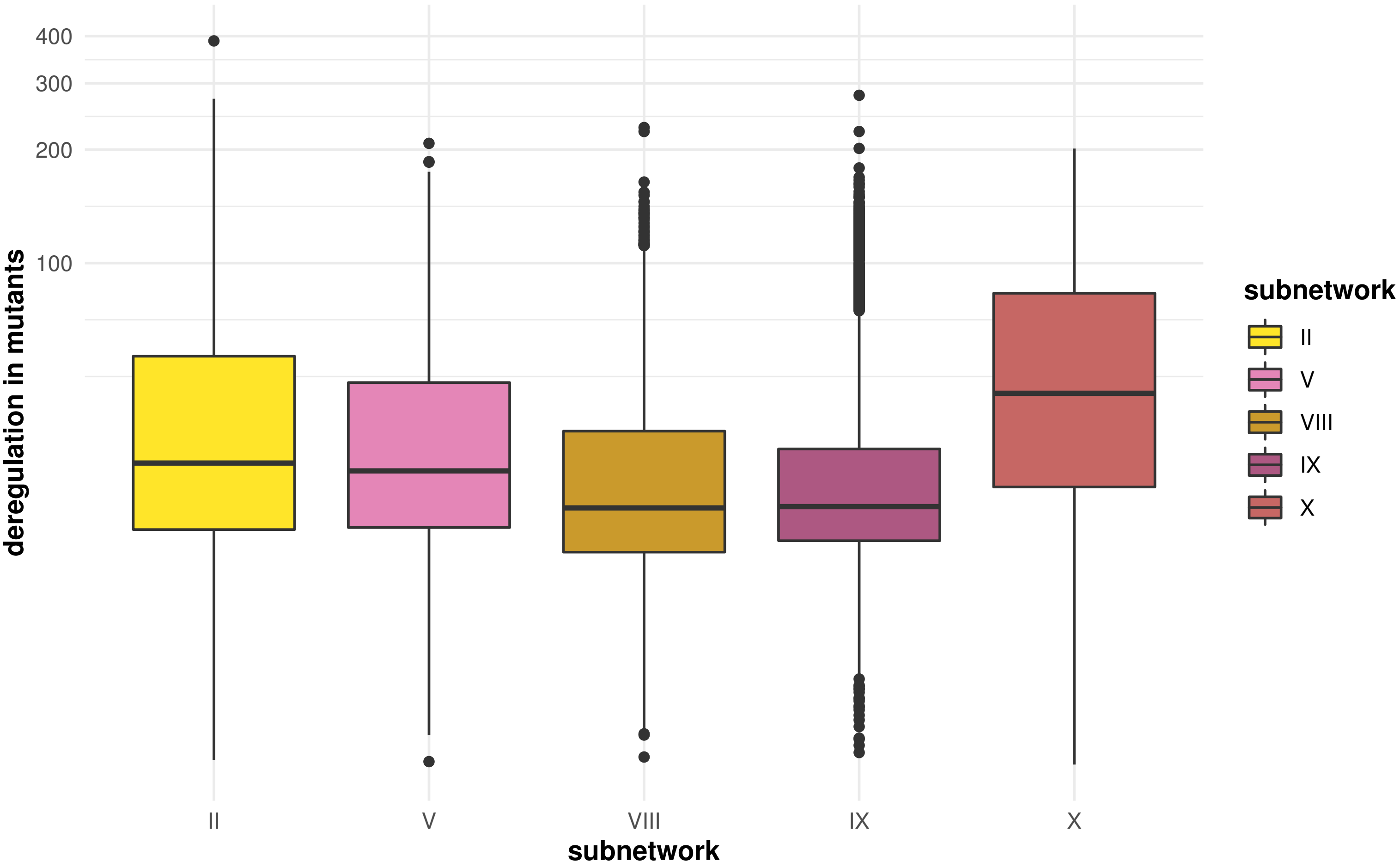
400
300
200
100
deregulation in mutants
II
V
VIII
IX
X
subnetwork
subnetwork
II
V
VIII
IX
X

## SF6h

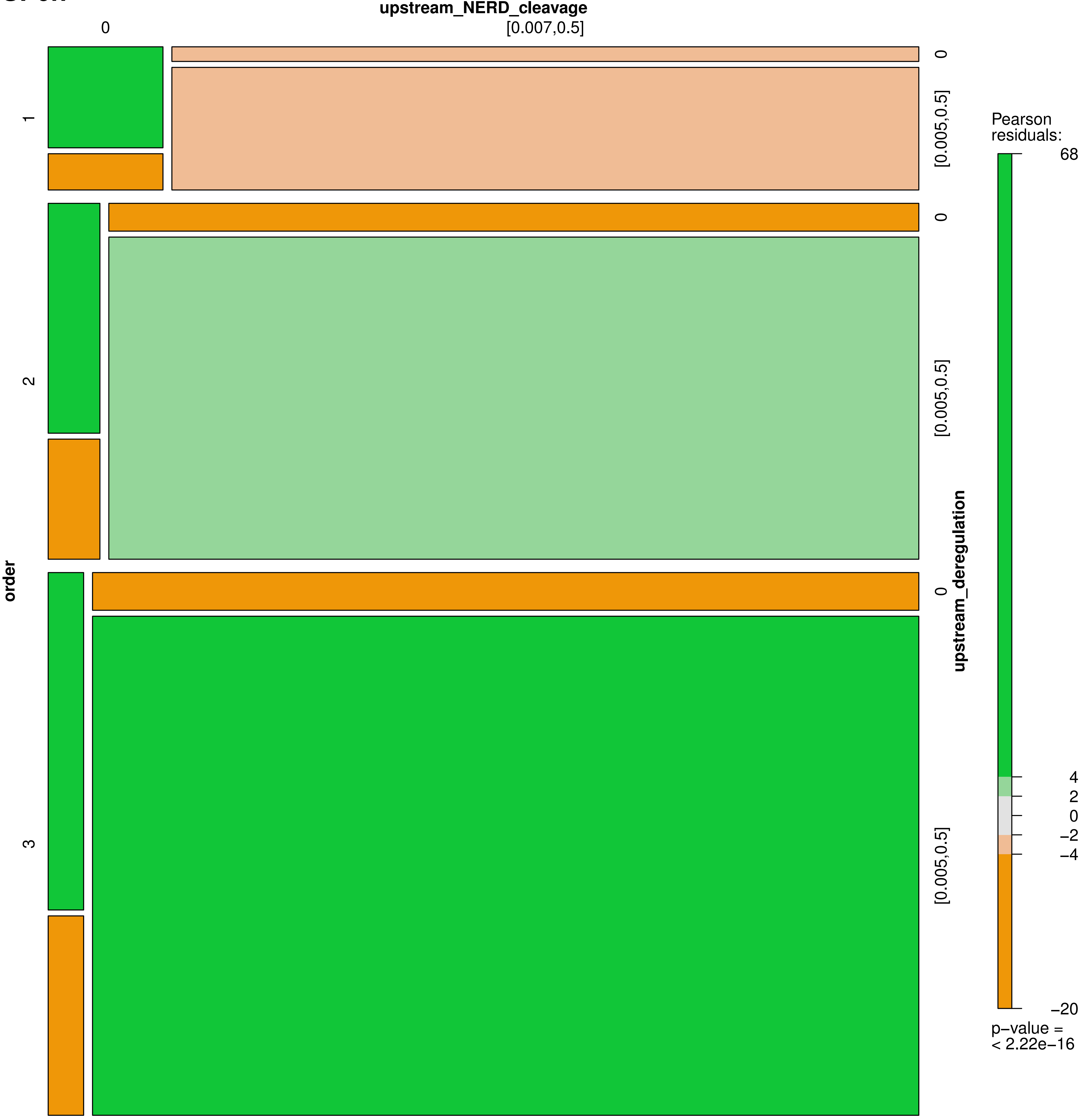

upstream_NERD_cleavage
0
[0.007,0.5]
order
1
2
3
upstream_deregulation
0
[0.005,0.5]
0
[0.005,0.5]
0
[0.005,0.5]
Pearson residuals:
68
4
2
0
−2
−4
−20
p−value =
< 2.22e−16

**SF7a**

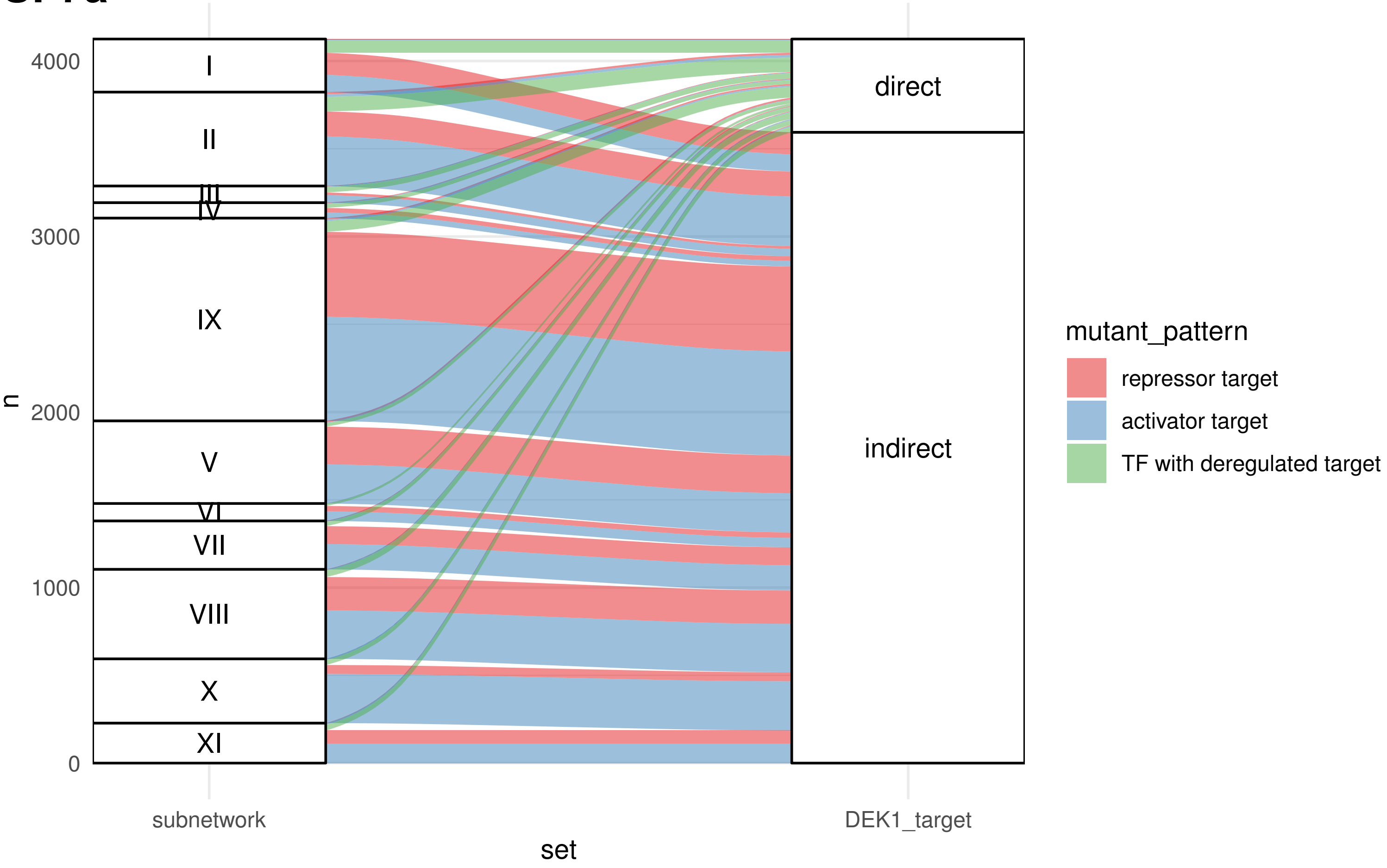

4000
3000
2000
1000
0
n
I
II
III
IV
IX
V
VI
VII
VIII
X
XI
direct
indirect
subnetwork
DEK1_target
set
mutant_pattern
repressor target
activator target
TF with deregulated target

SF7b

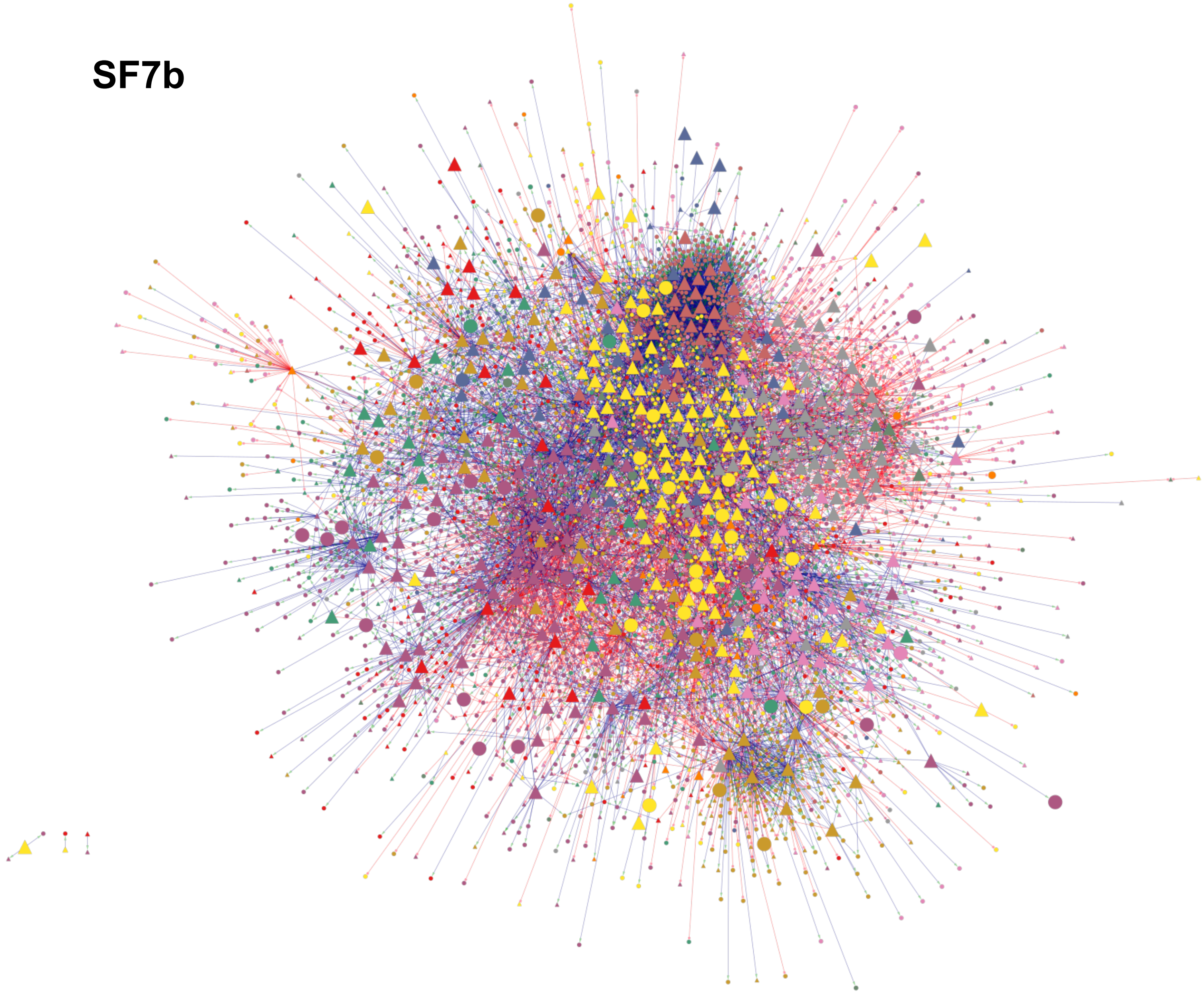

SF7c

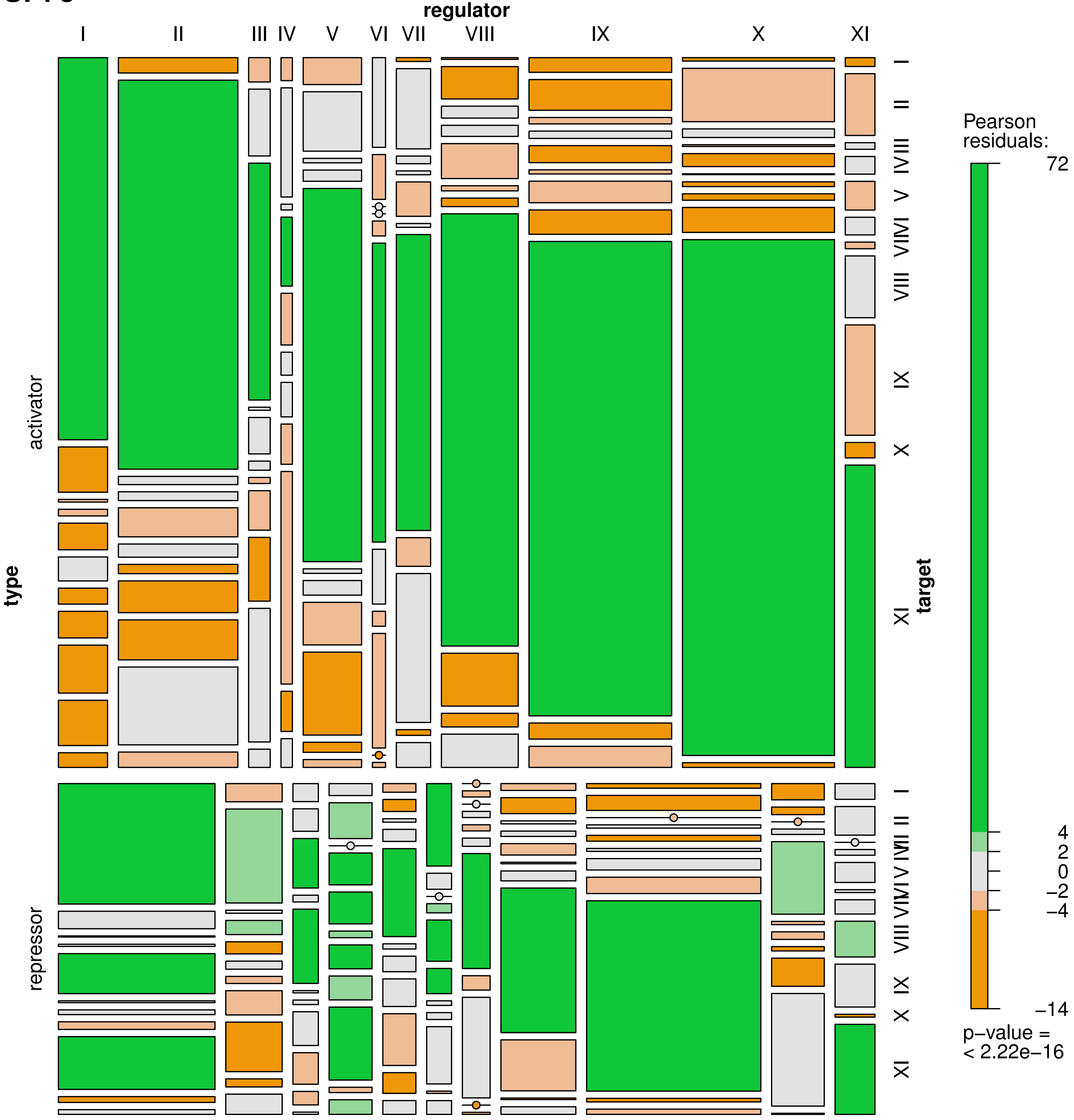

regulator
I
II
III
IV
V
VI
VII
VIII
IX
X
XI
type
activator
repressor
target
Pearson residuals:
72
4
2
0
−2
−4
−14
p−value = < 2.22e−16

**SF8a**

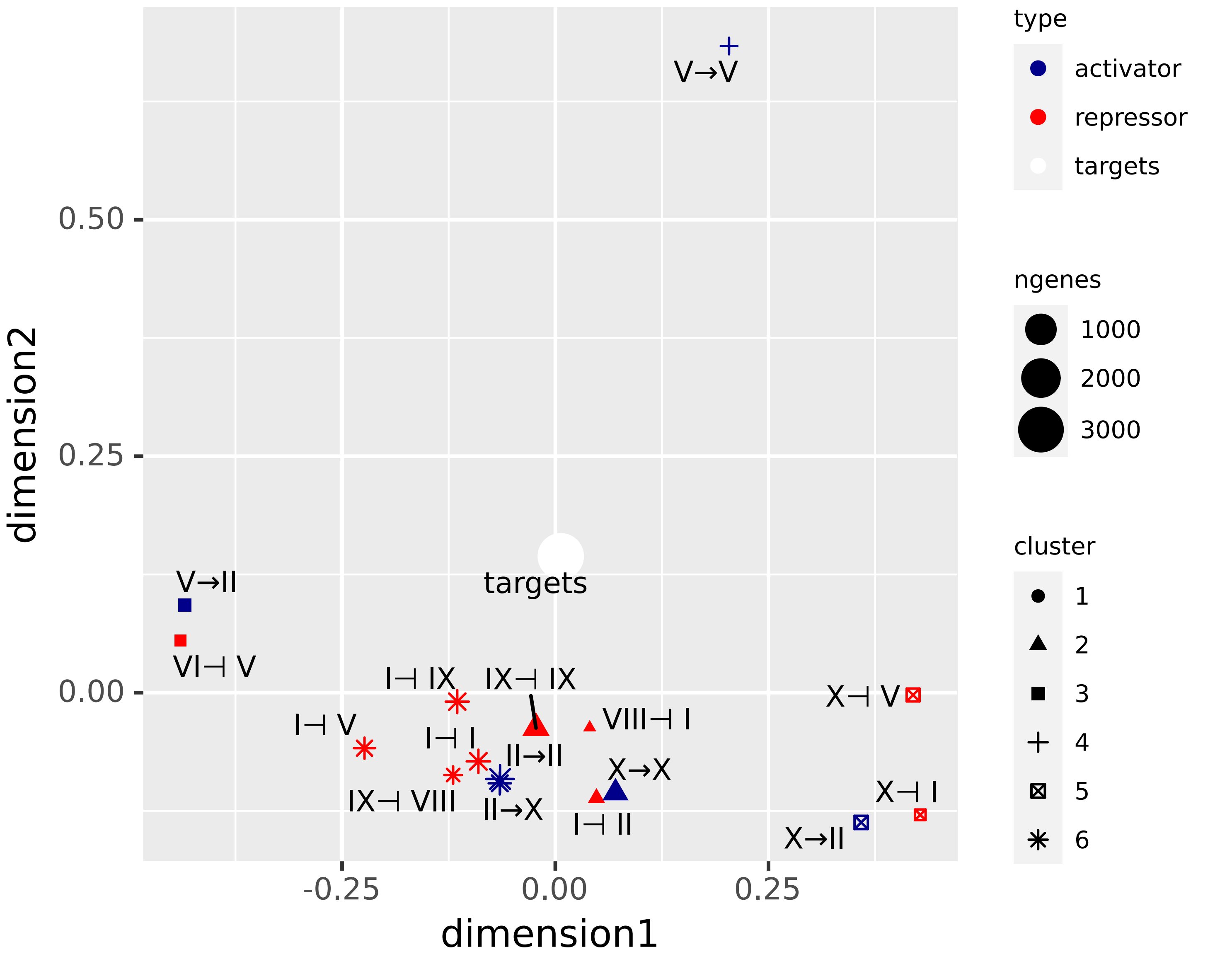
V→V
V→II
targets
VI⊣ V
I⊣ IX
IX⊣ IX
I⊣ V
I⊣ I
II→II
VIII⊣ I
X→X
IX⊣ VIII
II→X
I⊣ II
X⊣ V
X⊣ I
X→II
0.50
0.25
0.00
-0.25
0.00
0.25
dimension2
dimension1
type
activator
repressor
targets
ngenes
1000
2000
3000
cluster
1
2
3
4
5
6

**SF8b**

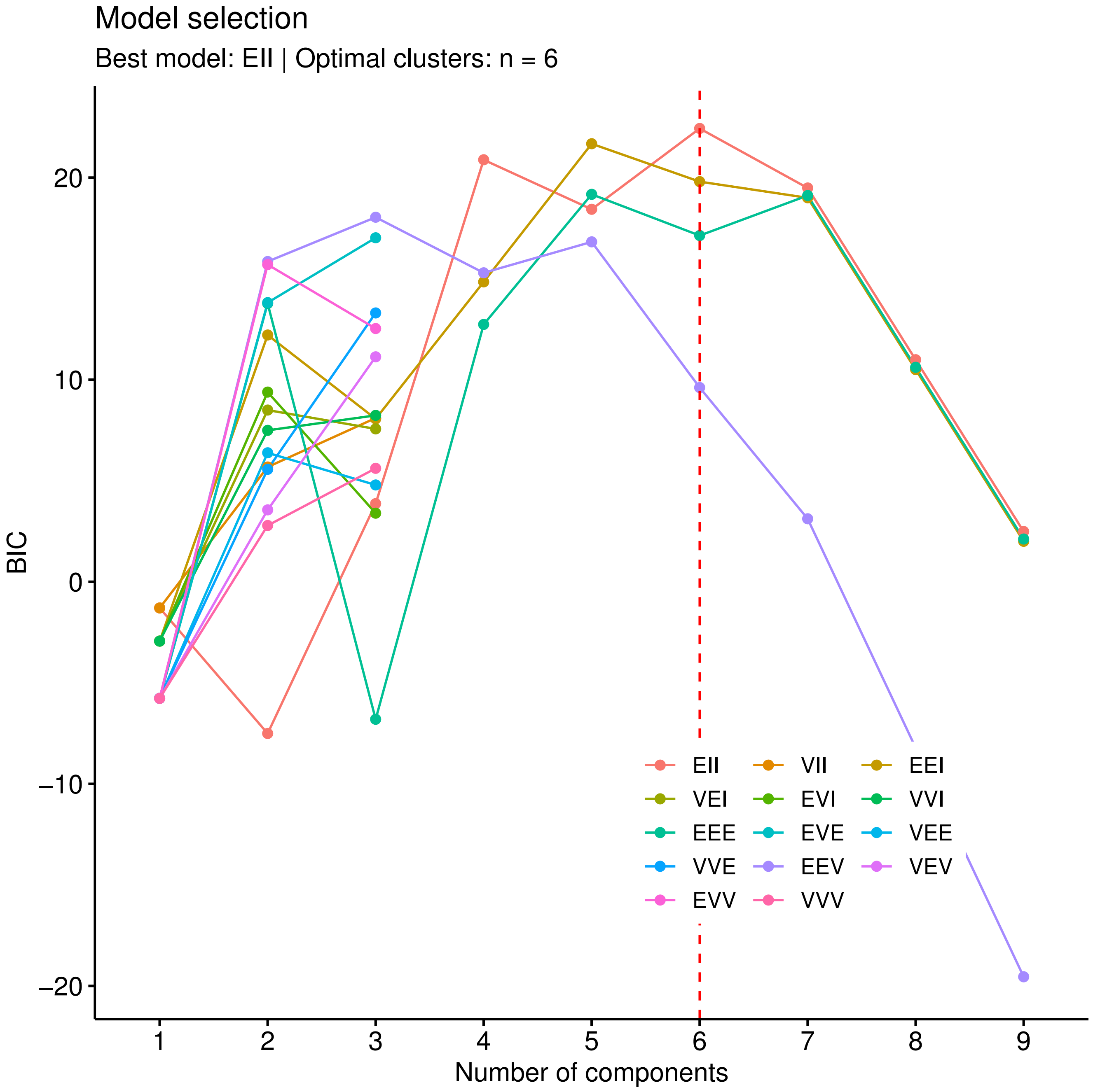
Model selection
Best model: EII | Optimal clusters: n = 6
BIC
20
10
0
−10
−20
1
2
3
4
5
6
7
8
9
Number of components
EII
VII
EEI
VEI
EVI
VVI
EEE
EVE
VEE
VVE
EEV
VEV
EVV
VVV

**SF8c**

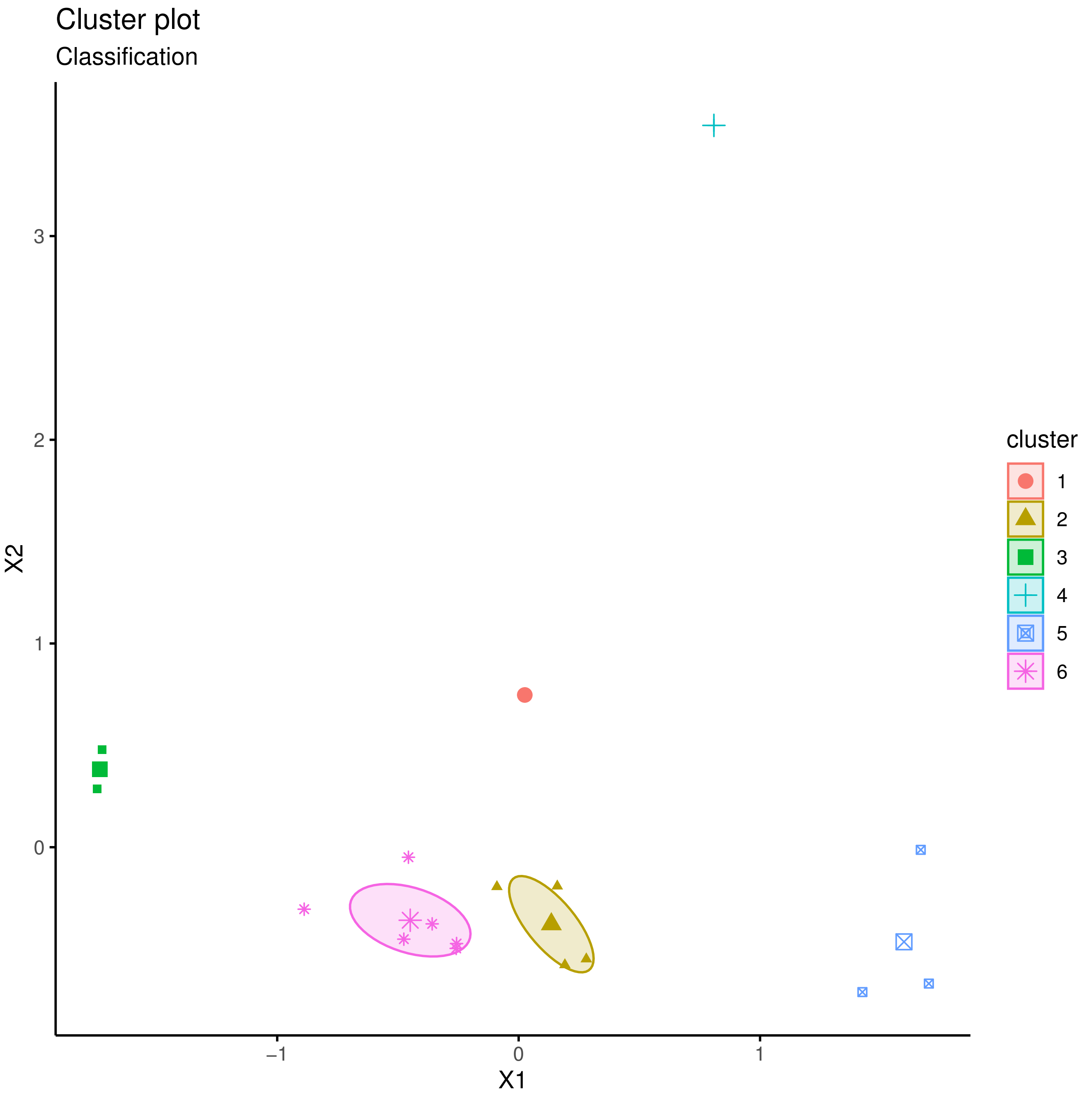

**SF8d**

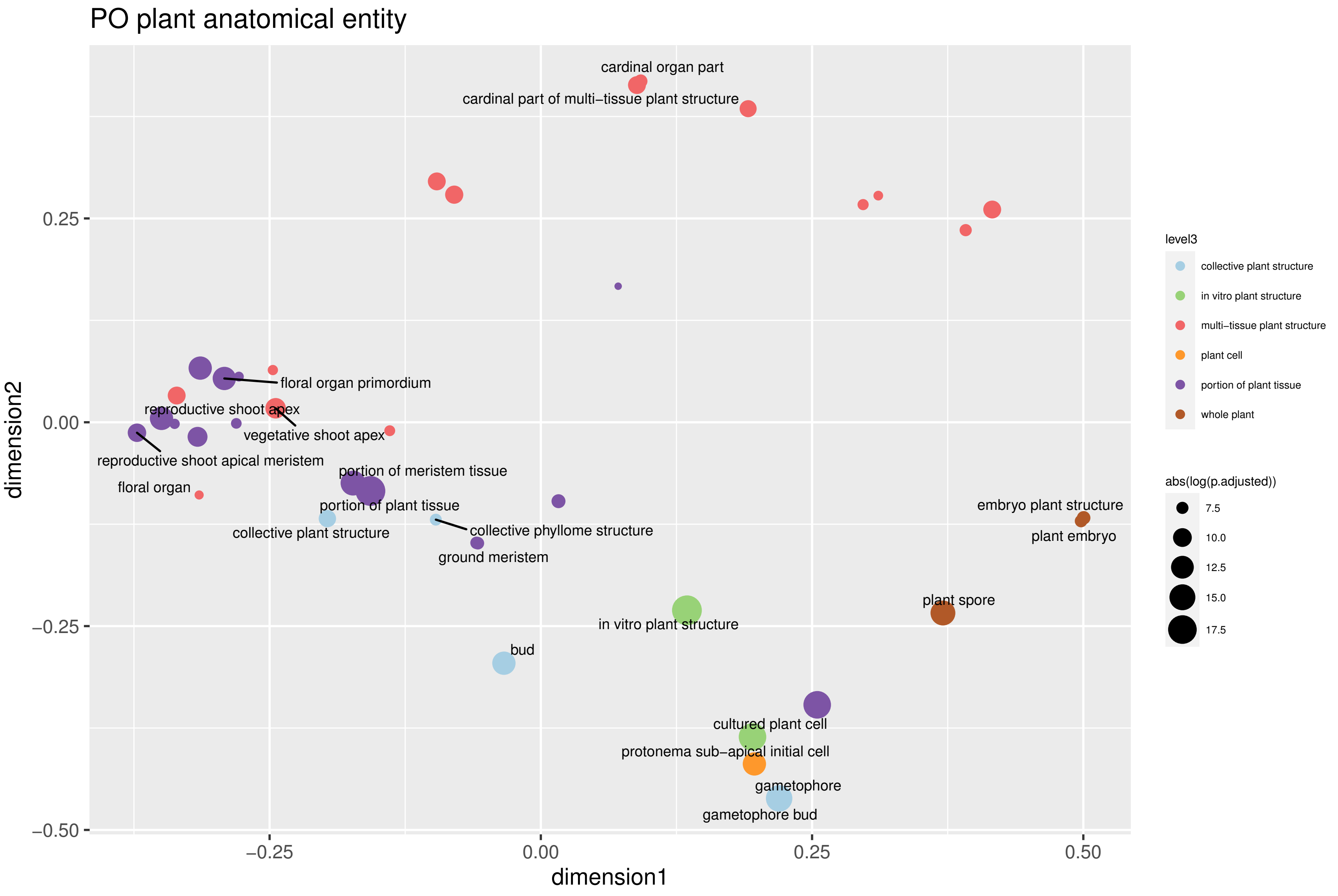

PO plant anatomical entity
cardinal organ part
cardinal part of multi−tissue plant structure
floral organ primordium
reproductive shoot apex
vegetative shoot apex
reproductive shoot apical meristem
floral organ
portion of meristem tissue
portion of plant tissue
collective plant structure
collective phyllome structure
ground meristem
embryo plant structure
plant embryo
plant spore
in vitro plant structure
bud
cultured plant cell
protonema sub−apical initial cell
gametophore
gametophore bud
0.25
0.00
−0.25
−0.50
−0.25
0.00
0.25
0.50
dimension1
dimension2
level3
collective plant structure
in vitro plant structure
multi−tissue plant structure
plant cell
portion of plant tissue
whole plant
abs(log(p.adjusted))
7.5
10.0
12.5
15.0
17.5

**SF8e**

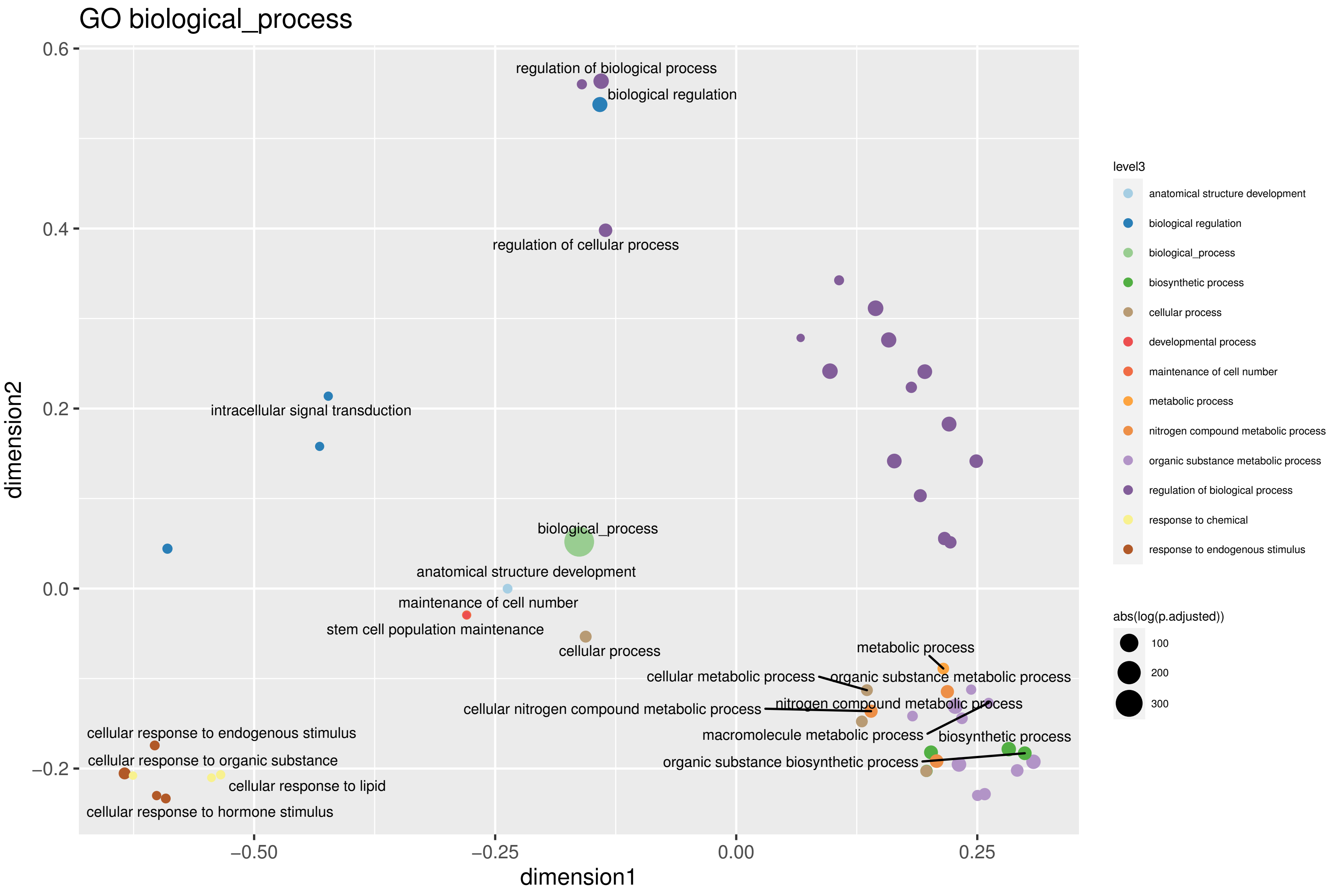
GO biological_process
regulation of biological process
biological regulation
regulation of cellular process
intracellular signal transduction
biological_process
anatomical structure development
maintenance of cell number
stem cell population maintenance
cellular process
metabolic process
cellular metabolic process
organic substance metabolic process
cellular nitrogen compound metabolic process
nitrogen compound metabolic process
macromolecule metabolic process
biosynthetic process
organic substance biosynthetic process
cellular response to endogenous stimulus
cellular response to organic substance
cellular response to lipid
cellular response to hormone stimulus
0.6
0.4
0.2
0.0
−0.2
−0.50
−0.25
0.00
0.25
dimension1
dimension2
level3
anatomical structure development
biological regulation
biological_process
biosynthetic process
cellular process
developmental process
maintenance of cell number
metabolic process
nitrogen compound metabolic process
organic substance metabolic process
regulation of biological process
response to chemical
response to endogenous stimulus
abs(log(p.adjusted))
100
200
300

# SF9

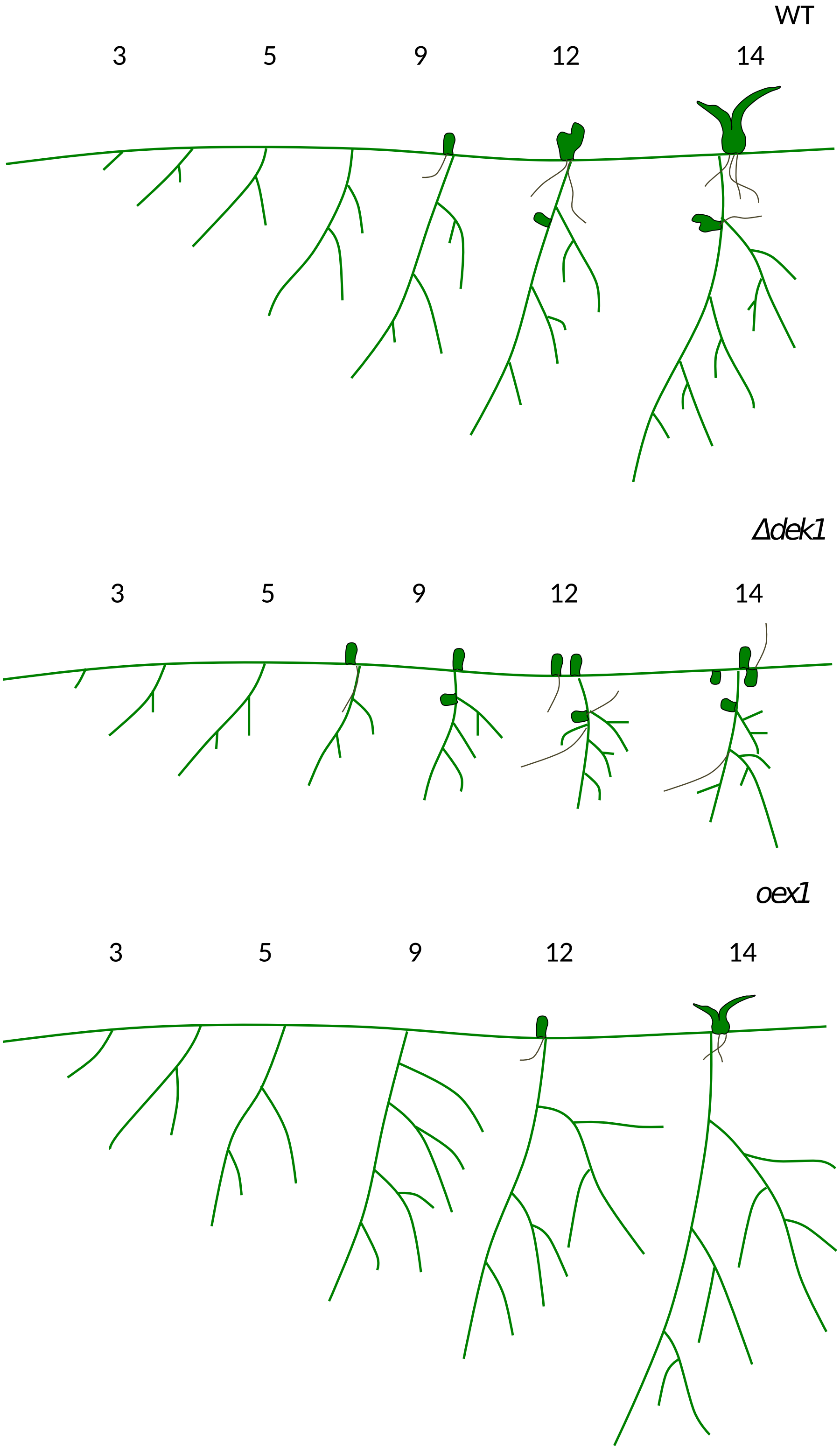


| | primary filaments | side branches | bud initiation | gametophore development |
|---|---|---|---|---|
| *Δdek1* | normal | reduced elongation | increased | arrested |
| *oex1* | elongated | increased elongation | reduced | normal |

# SF10

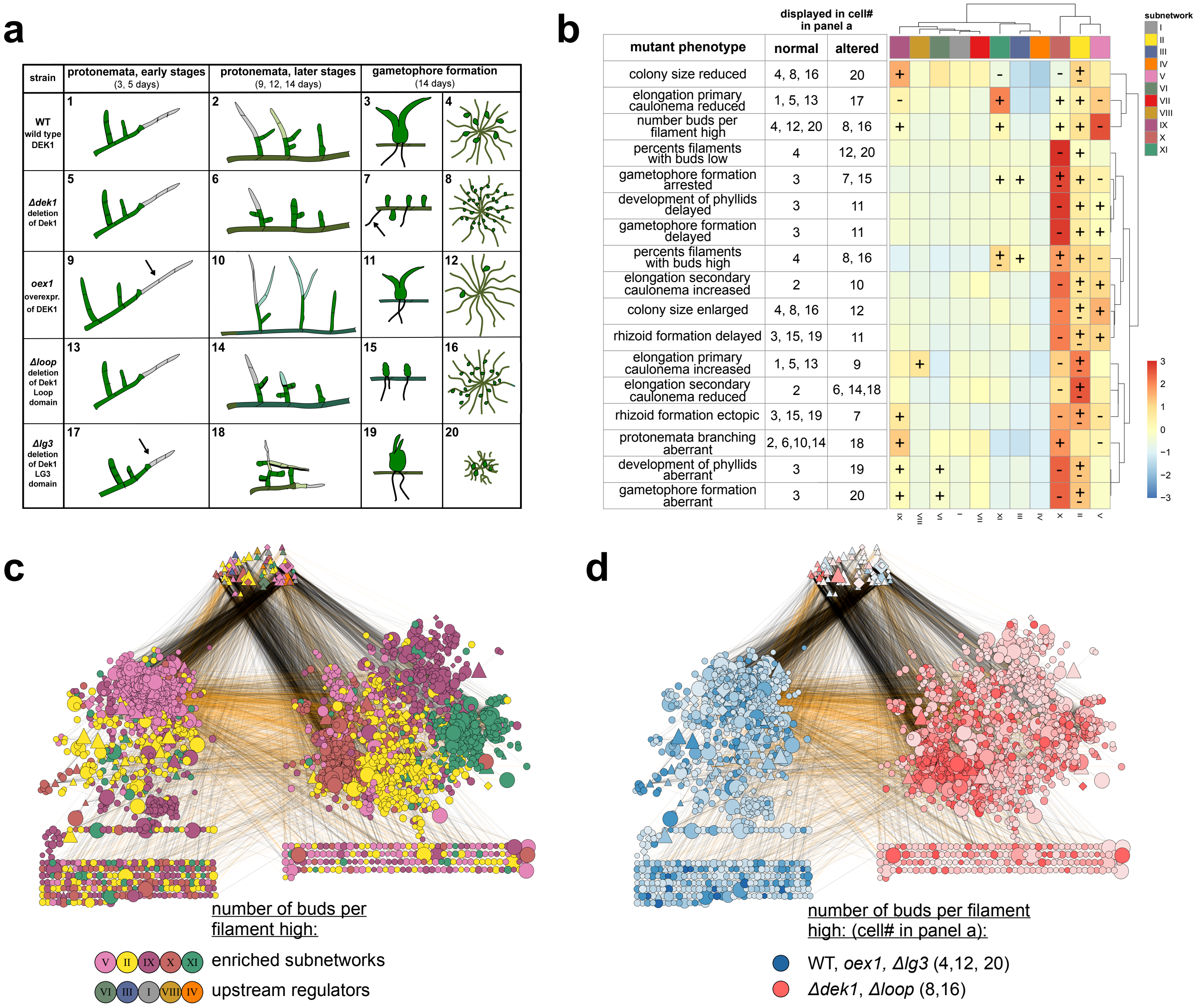
a
b
c
d
strain
protonemata, early stages
(3, 5 days)
protonemata, later stages
(9, 12, 14 days)
gametophore formation
(14 days)
WT
wild type
DEK1
Δdek1
deletion
of Dek1
oex1
overexpr.
of DEK1
Δloop
deletion
of Dek1
Loop
domain
Δlg3
deletion
of Dek1
LG3
domain
displayed in cell#
in panel a
mutant phenotype
normal
altered
colony size reduced
4, 8, 16
20
elongation primary caulonema reduced
1, 5, 13
17
number buds per filament high
4, 12, 20
8, 16
percents filaments with buds low
4
12, 20
gametophore formation arrested
3
7, 15
development of phyllids delayed
3
11
gametophore formation delayed
3
11
percents filaments with buds high
4
8, 16
elongation secondary caulonema increased
2
10
colony size enlarged
4, 8, 16
12
rhizoid formation delayed
3, 15, 19
11
elongation primary caulonema increased
1, 5, 13
9
elongation secondary caulonema reduced
2
6, 14,18
rhizoid formation ectopic
3, 15, 19
7
protonemata branching aberrant
2, 6,10,14
18
development of phyllids aberrant
3
19
gametophore formation aberrant
3
20
subnetwork
I
II
III
IV
V
VI
VII
VIII
IX
X
XI
number of buds per filament high:
enriched subnetworks
upstream regulators
number of buds per filament high: (cell# in panel a):
WT, oex1, Δlg3 (4,12, 20)
Δdek1, Δloop (8,16)

SF10e

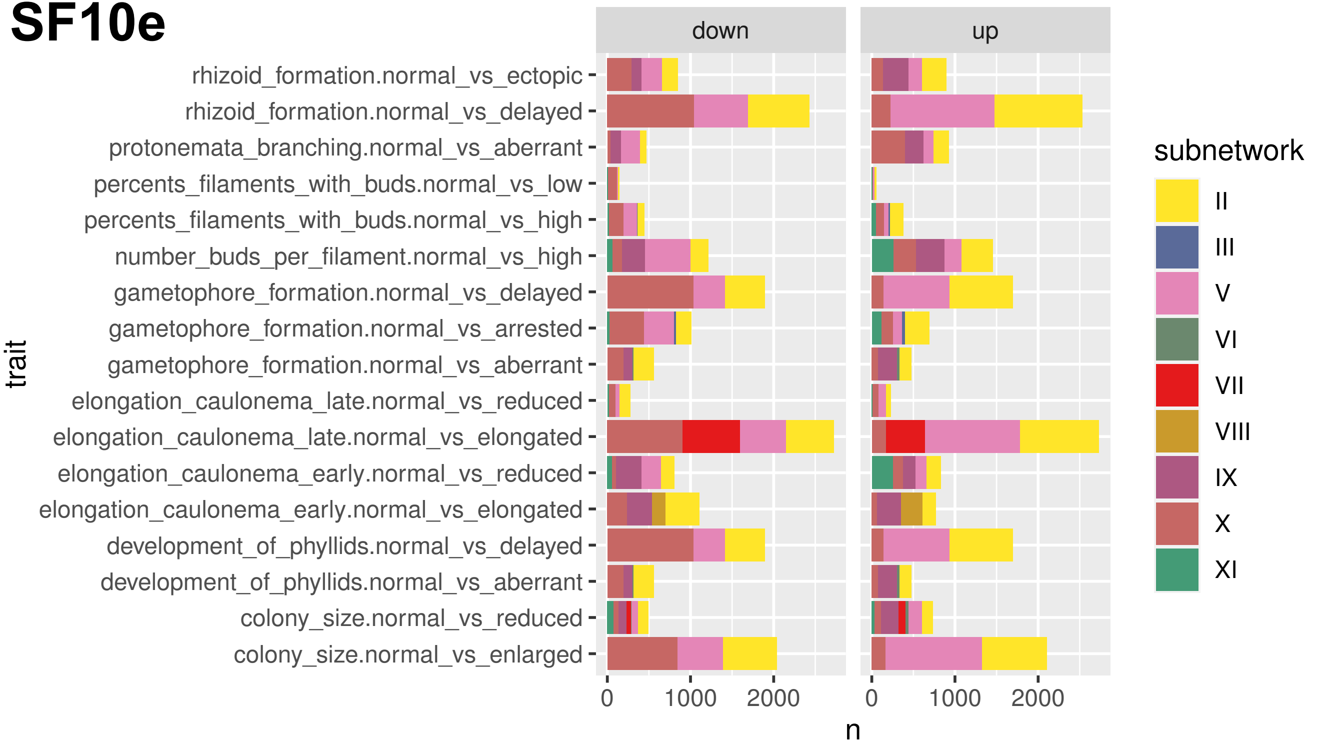
down
up
rhizoid_formation.normal_vs_ectopic
rhizoid_formation.normal_vs_delayed
protonemata_branching.normal_vs_aberrant
percents_filaments_with_buds.normal_vs_low
percents_filaments_with_buds.normal_vs_high
number_buds_per_filament.normal_vs_high
gametophore_formation.normal_vs_delayed
gametophore_formation.normal_vs_arrested
gametophore_formation.normal_vs_aberrant
elongation_caulonema_late.normal_vs_reduced
elongation_caulonema_late.normal_vs_elongated
elongation_caulonema_early.normal_vs_reduced
elongation_caulonema_early.normal_vs_elongated
development_of_phyllids.normal_vs_delayed
development_of_phyllids.normal_vs_aberrant
colony_size.normal_vs_reduced
colony_size.normal_vs_enlarged
trait
0
1000
2000
0
1000
2000
n
subnetwork
II
III
V
VI
VII
VIII
IX
X
XI

SF10f

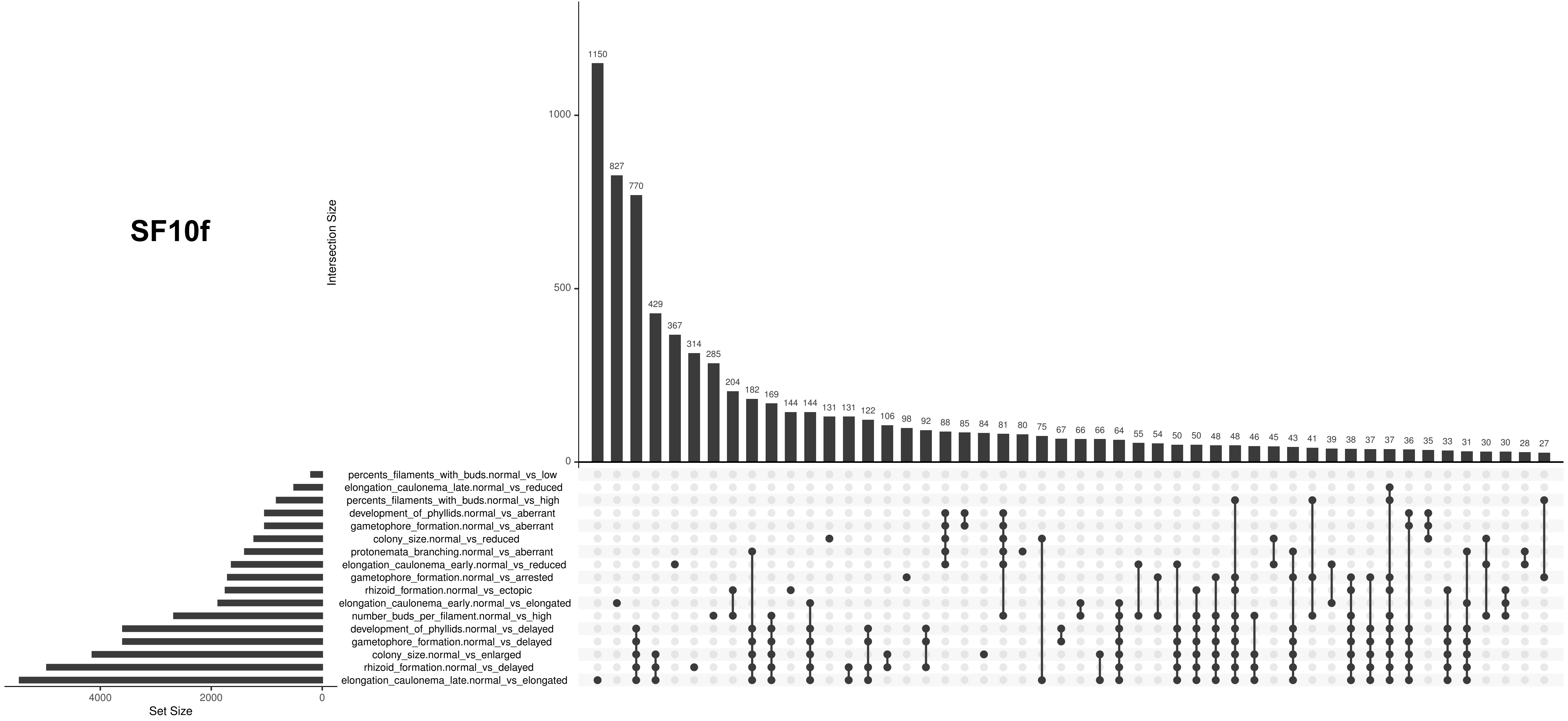

Intersection Size
1150
827
770
429
367
314
285
204
182
169
144
144
131
131
122
106
98
92
88
85
84
81
80
75
67
66
66
64
55
54
50
50
48
48
46
45
43
41
39
38
37
37
36
35
33
31
30
30
28
27
1000
500
0
percents_filaments_with_buds.normal_vs_low
elongation_caulonema_late.normal_vs_reduced
percents_filaments_with_buds.normal_vs_high
development_of_phyllids.normal_vs_aberrant
gametophore_formation.normal_vs_aberrant
colony_size.normal_vs_reduced
protonemata_branching.normal_vs_aberrant
elongation_caulonema_early.normal_vs_reduced
gametophore_formation.normal_vs_arrested
rhizoid_formation.normal_vs_ectopic
elongation_caulonema_early.normal_vs_elongated
number_buds_per_filament.normal_vs_high
development_of_phyllids.normal_vs_delayed
gametophore_formation.normal_vs_delayed
colony_size.normal_vs_enlarged
rhizoid_formation.normal_vs_delayed
elongation_caulonema_late.normal_vs_elongated
4000
2000
0
Set Size

# SF10g

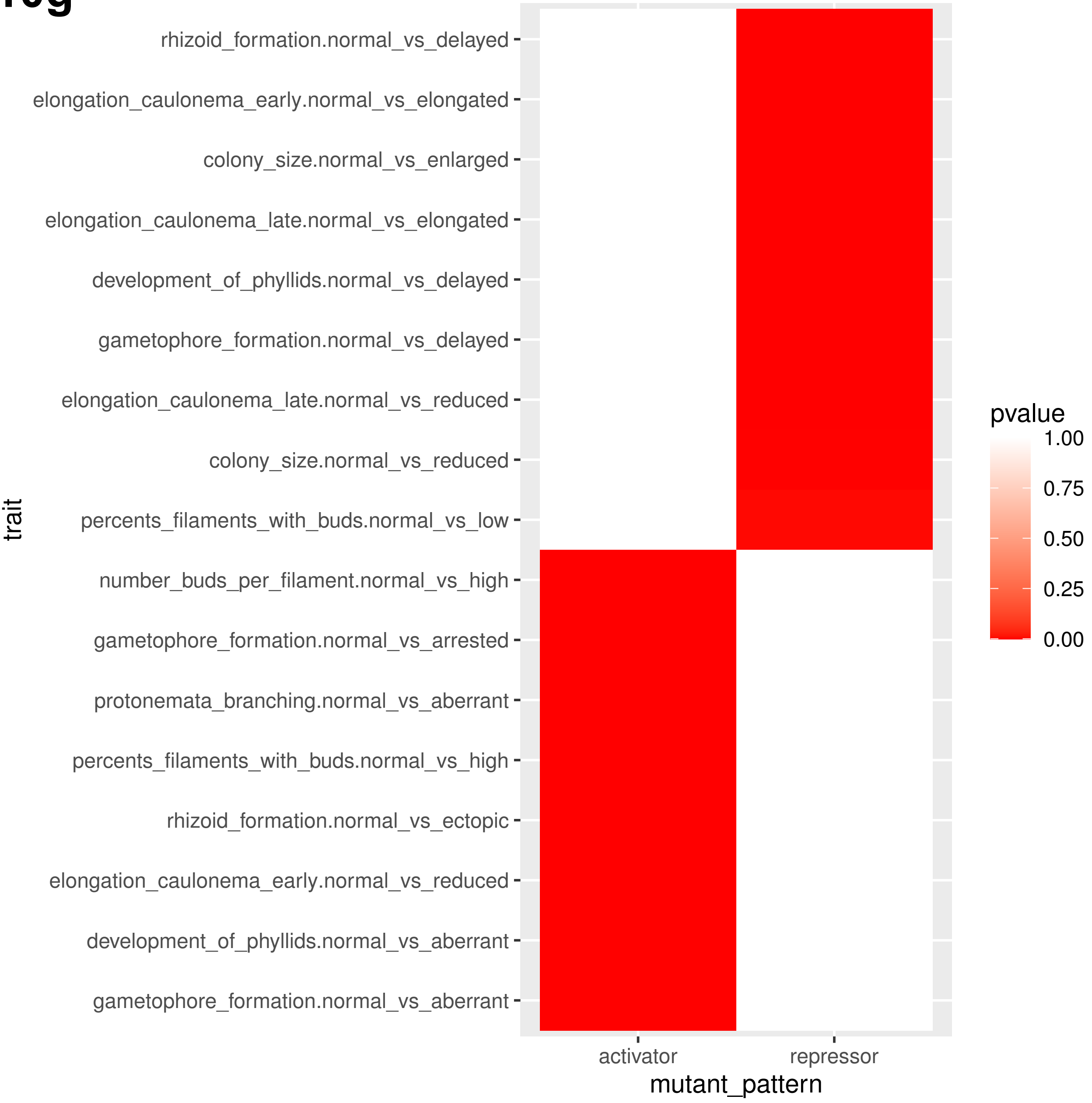

rhizoid_formation.normal_vs_delayed
elongation_caulonema_early.normal_vs_elongated
colony_size.normal_vs_enlarged
elongation_caulonema_late.normal_vs_elongated
development_of_phyllids.normal_vs_delayed
gametophore_formation.normal_vs_delayed
elongation_caulonema_late.normal_vs_reduced
colony_size.normal_vs_reduced
percents_filaments_with_buds.normal_vs_low
number_buds_per_filament.normal_vs_high
gametophore_formation.normal_vs_arrested
protonemata_branching.normal_vs_aberrant
percents_filaments_with_buds.normal_vs_high
rhizoid_formation.normal_vs_ectopic
elongation_caulonema_early.normal_vs_reduced
development_of_phyllids.normal_vs_aberrant
gametophore_formation.normal_vs_aberrant
trait
activator
repressor
mutant_pattern
pvalue
1.00
0.75
0.50
0.25
0.00

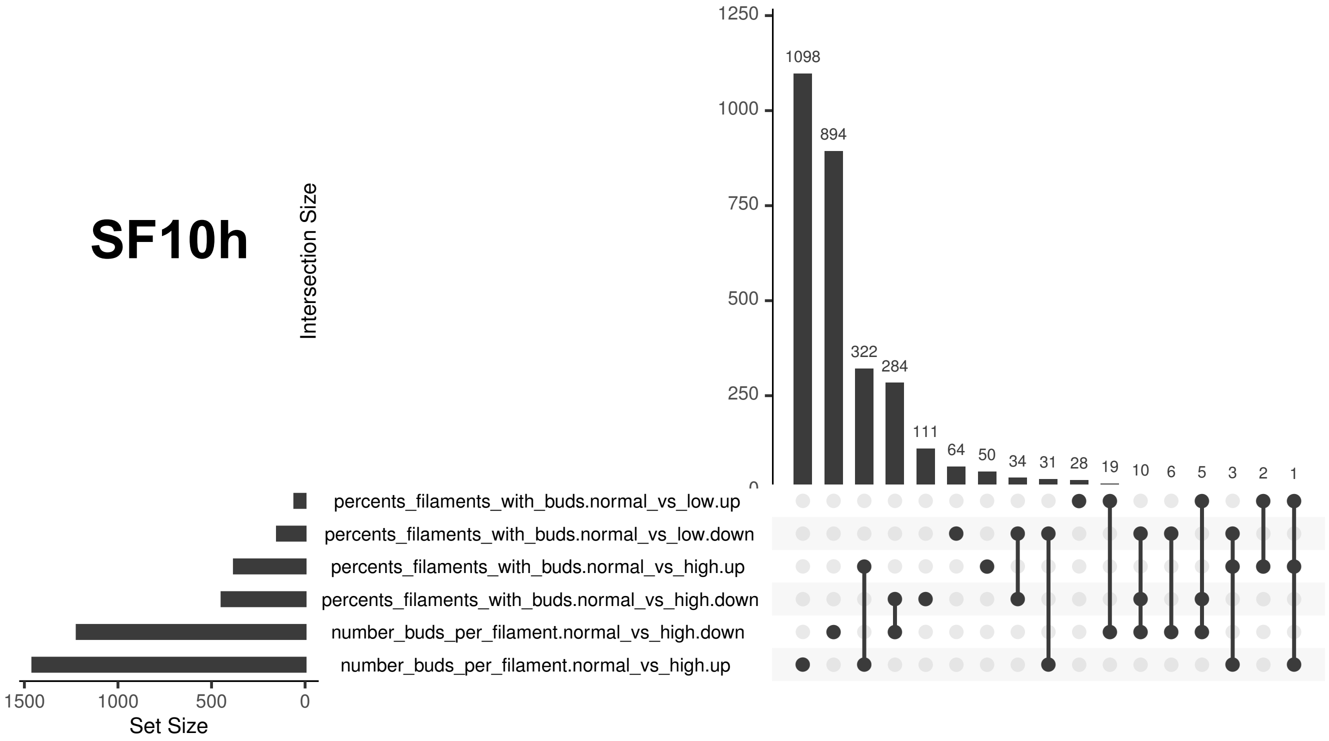
SF10h
Intersection Size
1250
1000
750
500
250
0
1098
894
322
284
111
64
50
34
31
28
19
10
6
5
3
2
1
percents_filaments_with_buds.normal_vs_low.up
percents_filaments_with_buds.normal_vs_low.down
percents_filaments_with_buds.normal_vs_high.up
percents_filaments_with_buds.normal_vs_high.down
number_buds_per_filament.normal_vs_high.down
number_buds_per_filament.normal_vs_high.up
1500
1000
500
0
Set Size

# SF10i

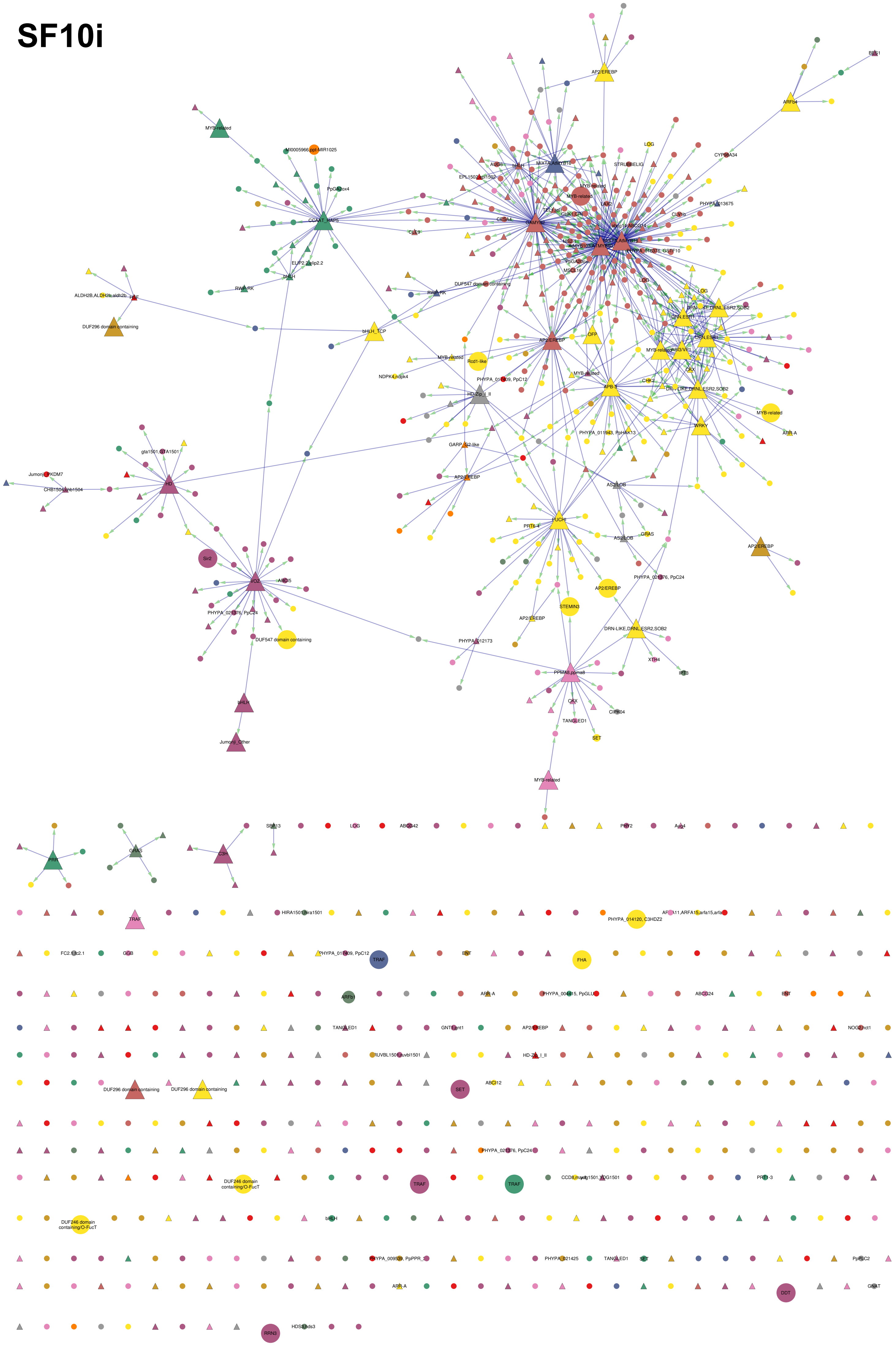

AP2/EREBP
ARFb4
BES1
MYB-related
MI0005966,ppt-MIR1025
PpGA2ox4
CCAAT_HAP5
ELIP2.2,elip2.2
bHLH
RWP-RK
ALDH2B,ALDH2b,aldh2b
HSF
DUF296 domain containing
bHLH_TCP
CYP98A34
LOG
STRUBBELIG
MIXTA,AtMYB16
EPL1502,epl1502
CESA4
CLE9
GAMYB2
CHK1,CRI
CLV1b
PHYPA_013675
PHYPA_016378, GSTF10
DUF547 domain containing
OFP
Rcd1-like
NDPK4,ndpk4
PHYPA_017409, PpC12
HD-Zip_I_II
APB-3
CHK2
WRKY
PHYPA_011943, PpHAK13
GARP_G2-like
ARR-A
gta1501,GTA1501
Jumonji_PKDM7
CHB1504,chb1504
HD
Sir2
VOZ
ABCI5
PHYPA_021376, PpC24
DUF547 domain containing
PUCHI
PRT6-4
AS2/LOB
GRAS
STEMIN3
PHYPA_012173
DRN-LIKE,DRNL,ESR2,SOB2
XTH4
IPT3
PPMA8,ppma8
CKX
CIPK04
TANGLED1
SET
Jumonji_Other
PRR
C3H
SBP13
ABCG42
PHY2
Aux4
TRAF
HIRA1501,hira1501
PHYPA_014120, C3HDZ2
FC2.1,fc2.1
GGB
ENT
FHA
ARFb1
ABCG24
GNT1,gnt1
NOG2,hct1
RUVBL1501,ruvbl1501
SET
ABCI12
DUF246 domain containing/O-FucT
PHYPA_009539, PpPPR_3
PHYPA_021425
PpPLC2
DDT
GNAT
RRN3
HDS3,hds3

**SF10j**

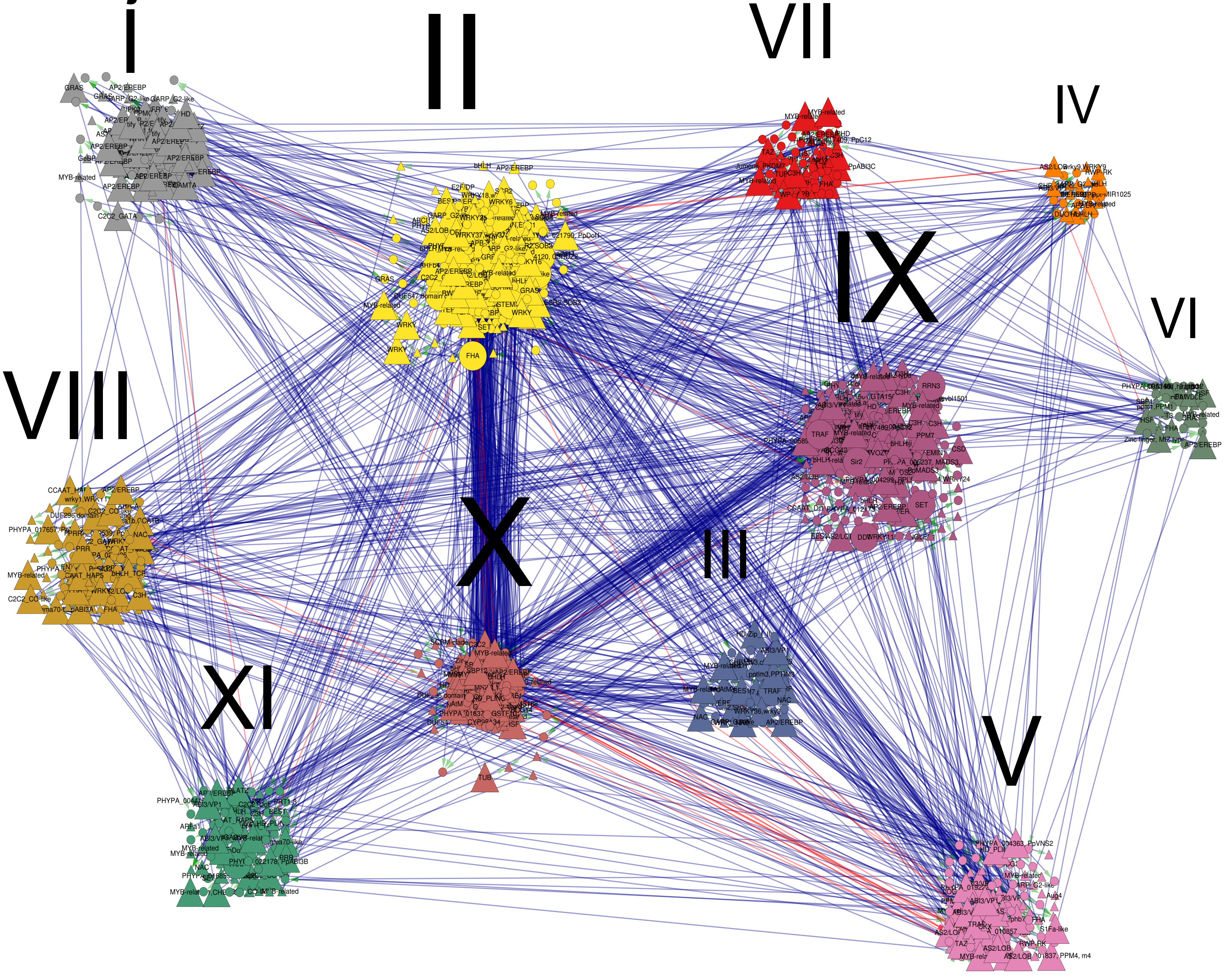

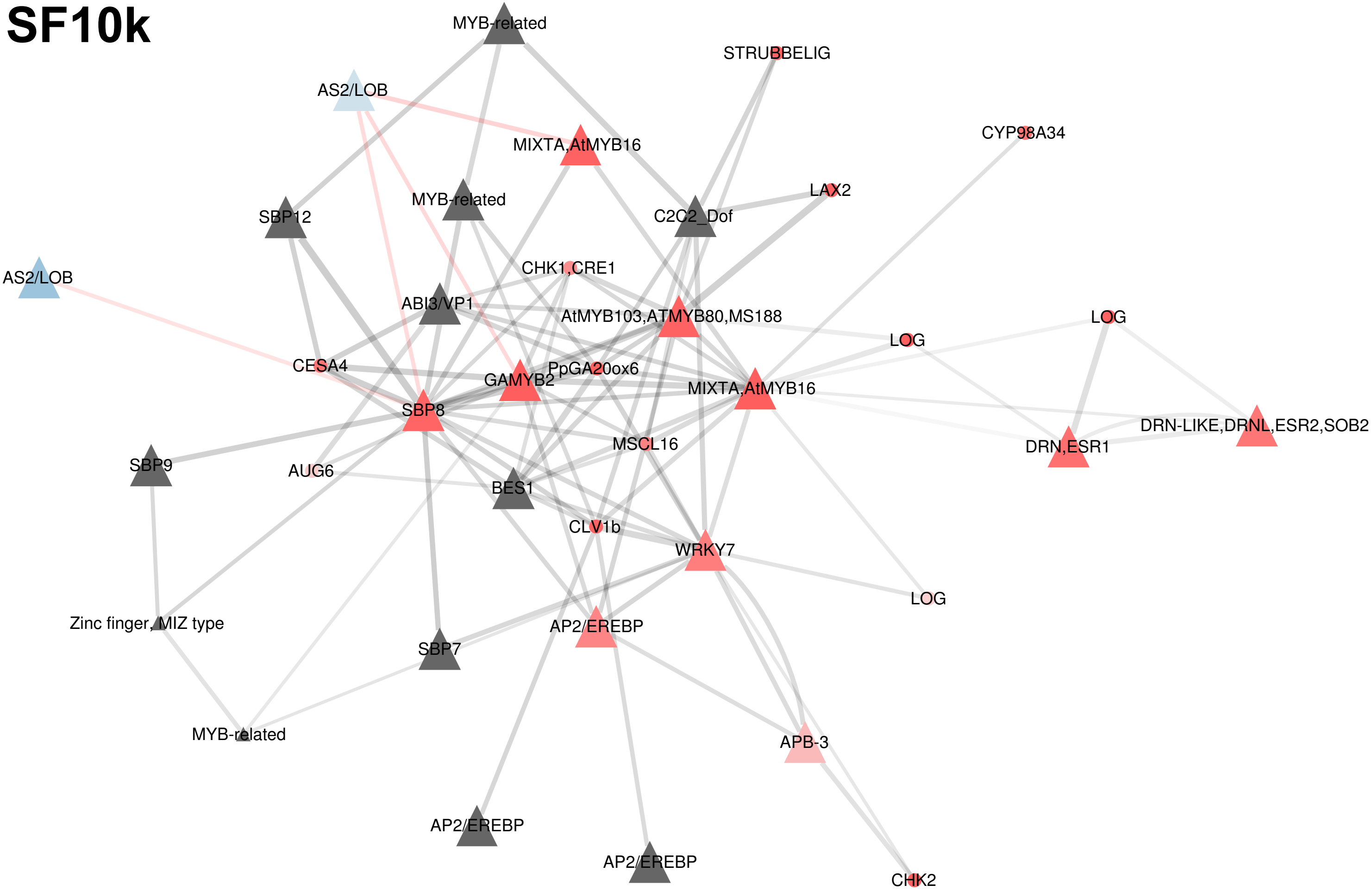
SF10k
MYB-related
STRUBBELIG
AS2/LOB
CYP98A34
MIXTA,AtMYB16
LAX2
MYB-related
SBP12
C2C2_Dof
CHK1,CRE1
AS2/LOB
ABI3/VP1
AtMYB103,ATMYB80,MS188
LOG
LOG
CESA4
PpGA20ox6
GAMYB2
MIXTA,AtMYB16
SBP8
DRN-LIKE,DRNL,ESR2,SOB2
DRN,ESR1
MSCL16
SBP9
AUG6
BES1
CLV1b
WRKY7
LOG
Zinc finger, MIZ type
AP2/EREBP
SBP7
MYB-related
APB-3
AP2/EREBP
AP2/EREBP
CHK2

SF10

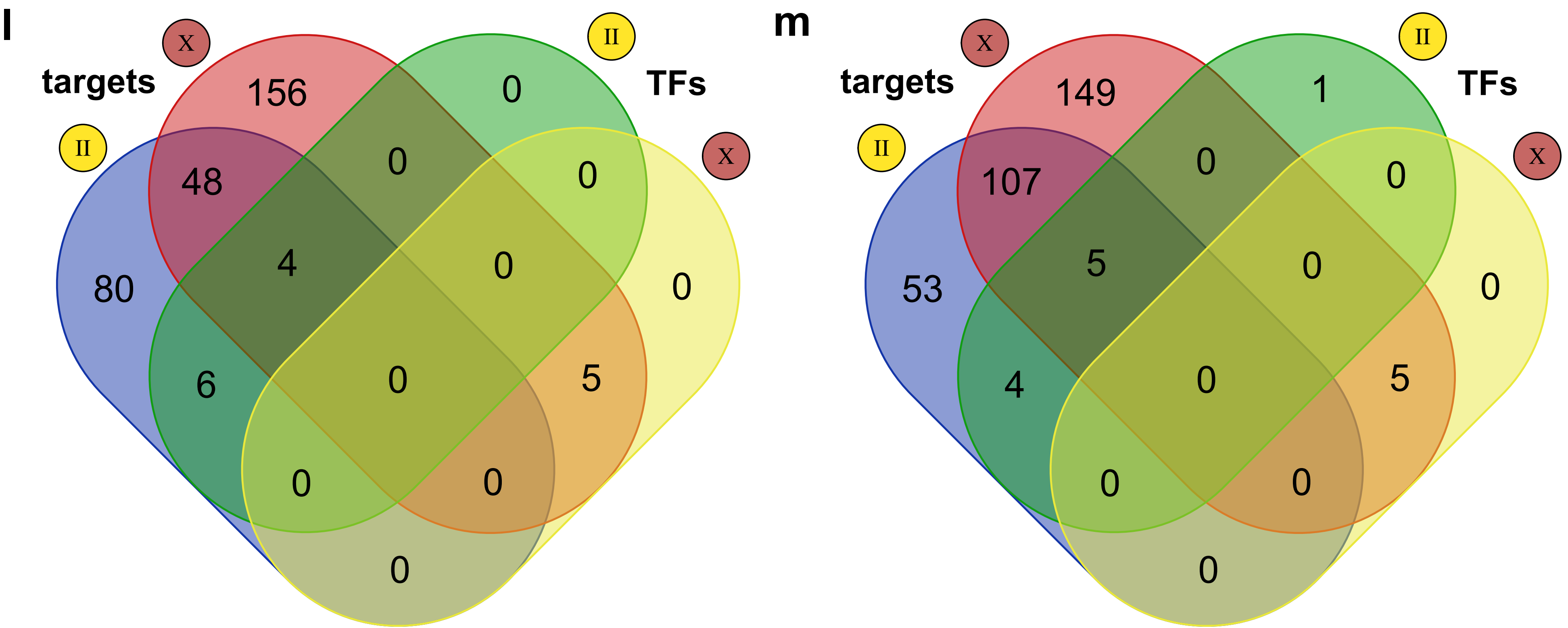
l
targets
X
II
TFs
X
II
156
0
48
0
0
80
4
0
0
6
0
5
0
0
0
m
targets
X
II
TFs
X
II
149
1
107
0
0
53
5
0
0
4
0
5
0
0
0

**SF11**

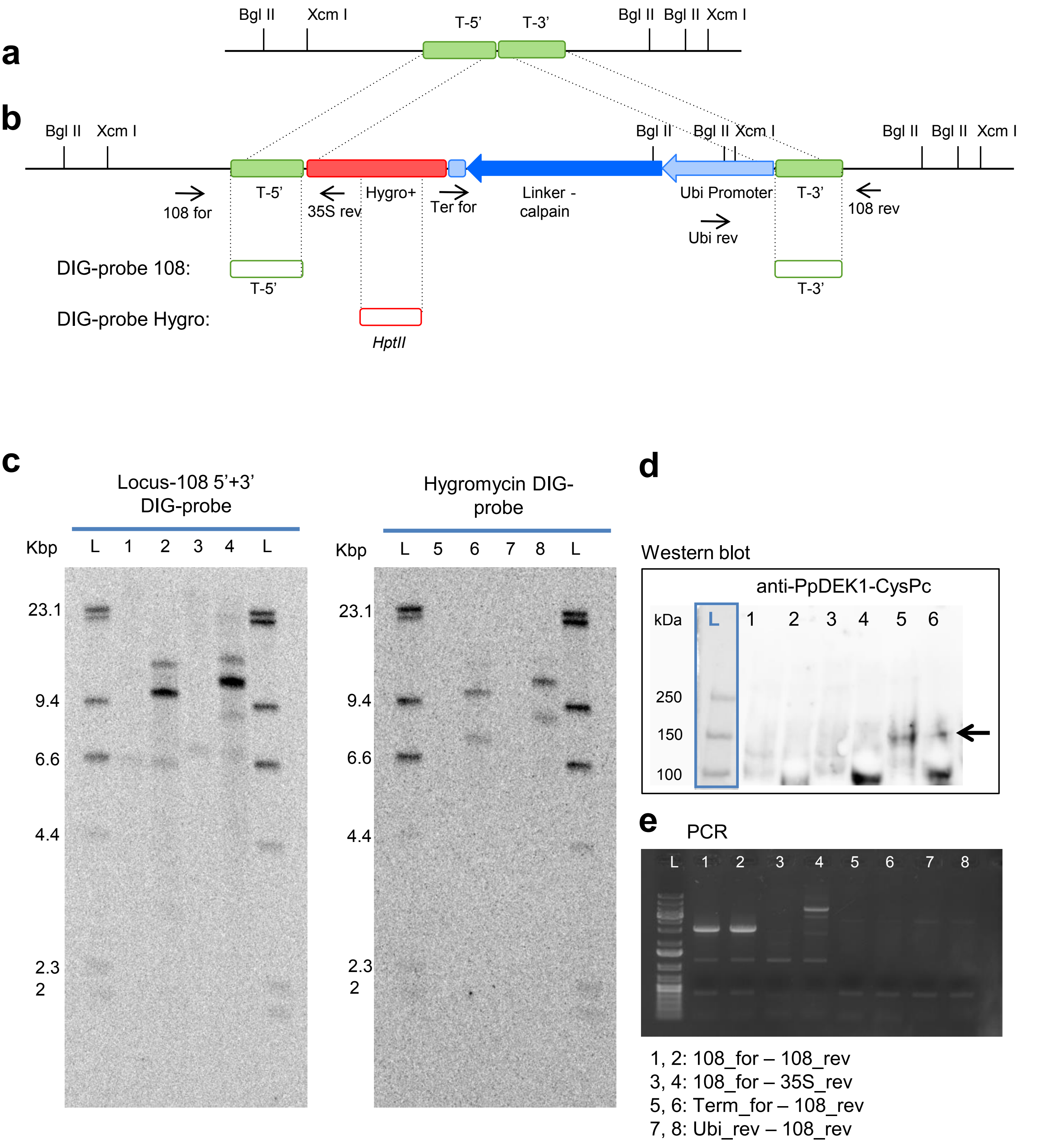
a
Bgl II
Xcm I
T-5'
T-3'
Bgl II
Bgl II
Xcm I
b
Bgl II
Xcm I
Bgl II
Bgl II
Xcm I
Bgl II
Bgl II
Xcm I
T-5'
108 for
35S rev
Hygro+
Ter for
Linker -
calpain
Ubi Promoter
Ubi rev
T-3'
108 rev
DIG-probe 108:
T-5'
T-3'
DIG-probe Hygro:
HptII
c
Locus-108 5'+3'
DIG-probe
Kbp
L 1 2 3 4 L
23.1
9.4
6.6
4.4
2.3
2
Hygromycin DIG-
probe
Kbp
L 5 6 7 8 L
23.1
9.4
6.6
4.4
2.3
2
d
Western blot
anti-PpDEK1-CysPc
kDa
L 1 2 3 4 5 6
250
150
100
e
PCR
L 1 2 3 4 5 6 7 8
1, 2: 108_for – 108_rev
3, 4: 108_for – 35S_rev
5, 6: Term_for – 108_rev
7, 8: Ubi_rev – 108_rev

**SF11**

**f**

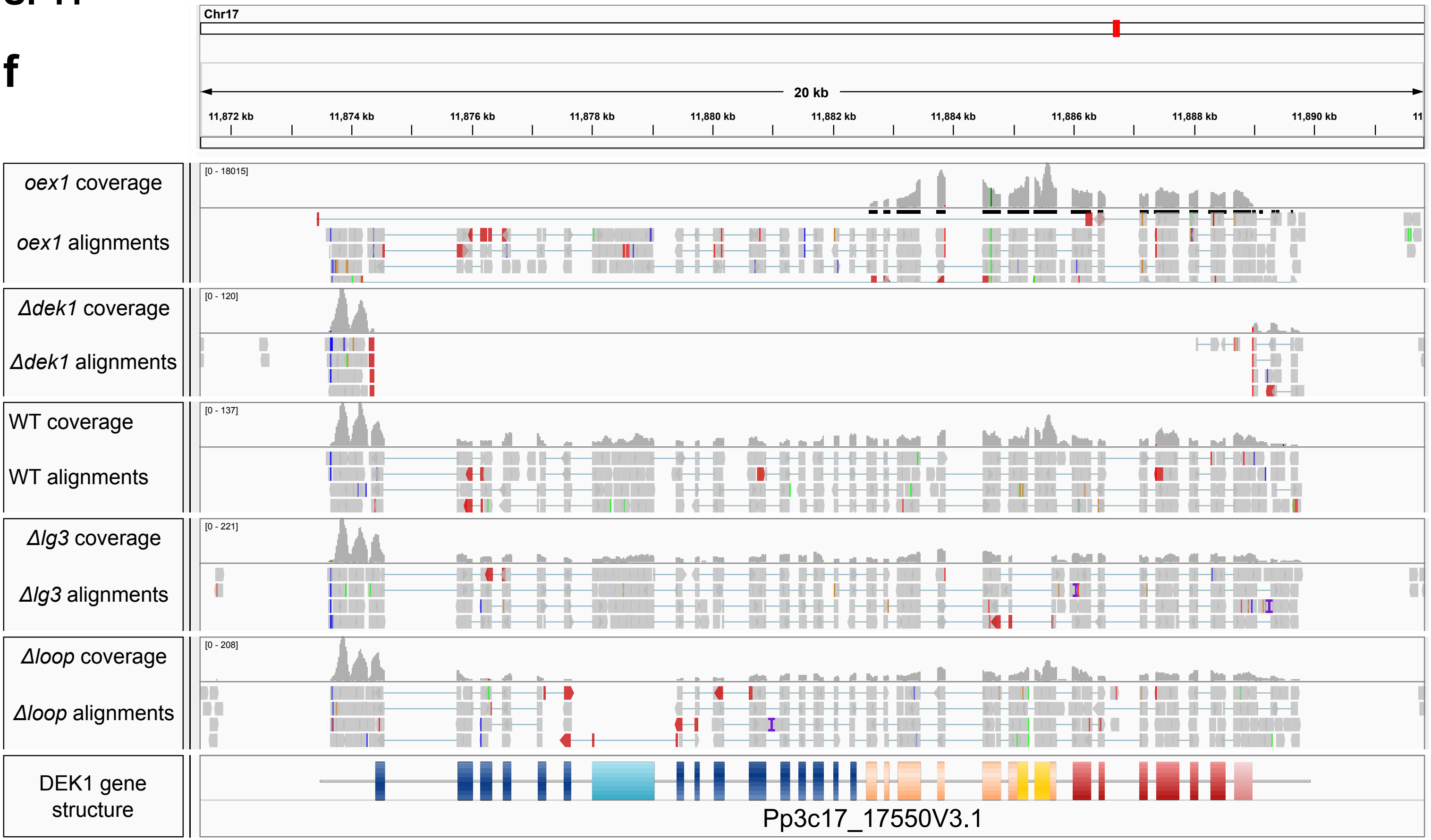
Chr17
20 kb
11,872 kb
11,874 kb
11,876 kb
11,878 kb
11,880 kb
11,882 kb
11,884 kb
11,886 kb
11,888 kb
11,890 kb
*oex1* coverage
*oex1* alignments
[0 - 18015]
*Δdek1* coverage
*Δdek1* alignments
[0 - 120]
WT coverage
WT alignments
[0 - 137]
*Δlg3* coverage
*Δlg3* alignments
[0 - 221]
*Δloop* coverage
*Δloop* alignments
[0 - 208]
DEK1 gene structure
Pp3c17_17550V3.1

**g**

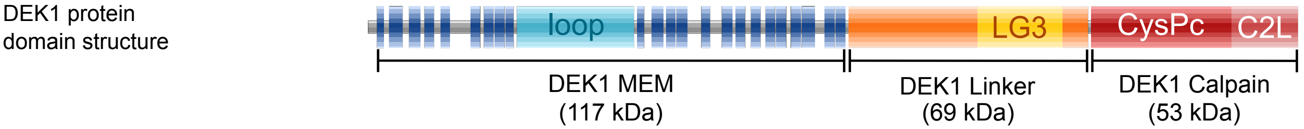
DEK1 protein domain structure
loop
LG3
CysPc
C2L
DEK1 MEM (117 kDa)
DEK1 Linker (69 kDa)
DEK1 Calpain (53 kDa)
